# Human-like sequential sound-to-meaning transfer drives artificial speech comprehension

**DOI:** 10.64898/2026.05.13.723203

**Authors:** Shenshen Zhang, Siqi Li, Ruolin Yang, Guanpeng Chen, Xing Tian, Qian Wang, Fang Fang

**Affiliations:** School of Psychological and Cognitive Sciences and Beijing Key Laboratory of Behavior and Mental Health, Peking University, 100871, Beijing, China; IDG/McGovern Institute for Brain Research, Peking University, 100871, Beijing, China; Peking-Tsinghua Center for Life Sciences, Academy for Advanced Interdisciplinary Studies, Peking University, 100871, Beijing, China; Key Laboratory of Machine Perception, Ministry of Education, Peking University, 100871, Beijing, China; New Cornerstone Science Laboratory, Center for Motor Control and Disease, Key Laboratory of Brain Functional Genomics, East China Normal University, Shanghai 200062, China; NYU-ECNU Institute of Brain and Cognitive Science at NYU Shanghai, 200062, Shanghai, China; Shanghai Key Laboratory of Brain Functional Genomics (Ministry of Education), School of Psychology and Cognitive Science, East China Normal University, 200062, Shanghai, China; Shanghai Frontiers Science Center of Artificial Intelligence and Deep Learning, Division of Arts and Sciences, New York University Shanghai, Shanghai, China; National Key Laboratory of General Artificial Intelligence, Peking University, 100871, Beijing, China

**Author notes:** Corresponding authors: Xing Tian, Qian Wang, Fang Fang.

## Abstract

Artificial intelligence has reached a pivotal threshold. Multimodal large models can approach human-level speech comprehension by rapidly transforming sound into meaning. However, whether this process relies on human-like mechanisms remains unknown. Here, we compared the human brain with twelve speech language models (SLMs) using a phonology–semantics confusion paradigm. Stereo-electroencephalography revealed two mechanisms of phonology-to-semantics (P2S) transfer in the human brain: a local sequential transformation within specific neuronal populations, and a global cross-regional hierarchy of P2S representations. Only brain–model alignment in the local sequential manner predicted model performance. Correspondingly, targeted lesioning of local sequential P2S-transfer model units markedly impaired comprehension performance, while activation steering of these units improved performance. In addition, such local sequential P2S-transfer model units were identified across languages. Together, this study establishes local sequential P2S transformation as a fundamental computational principle shared across biological and artificial intelligence, offering a mechanistic bridge for future brain-inspired speech systems.

## Introduction

Rapid and effortless speech comprehension is a hallmark of human cognition. In a fraction of a second, the brain converts fleeting acoustic waveforms into stable conceptual representations^1–4^. Recently, this biological monopoly has been challenged by the emergence of speech language models (SLMs). These systems, capable of processing raw audio inputs, have rapidly ascended to performance levels that rival human listeners^5–14^. This convergence offers a rare chance to tackle a core question about intelligence: do brains and machines reach meaning from sound by the same underlying computation, or do they merely converge on similar behavior through different routes?

Neuroscientific inquiry has long sought to map the trajectory of spoken language processing. Established models posit a division of labor, where phonological analysis relies on spectro-temporal decomposition in auditory cortices^15–24^, while semantic integration engages a distributed network of higher-order regions^4,21,25–27^. However, the specific mechanism bridging these two domains, phonology-to-semantics (P2S) transfer, remains a subject of intense debate. It is unclear whether this transfer is exclusively a product of slow, cumulative processing across a global cross-regional hierarchy, or if it also necessitates rapid, specialized computations within local neuronal populations.

Simultaneously, the “black box” nature of deep learning models presents a parallel challenge. Although extensive research has demonstrated that deep neural networks often develop internal representations analogous to those of the human brain in their hierarchical organization^28–55^, the functional relevance of these similarities is not fully understood^56^. Merely having a structure that correlates with the brain does not guarantee that a model operates like one. A critical, unanswered question is whether high-performing SLMs effectively replicate the fine-grained dynamics of human P2S transfer, and whether such mechanistic alignment is a prerequisite for superior comprehension capabilities.

To dissect these mechanisms, we leveraged a theoretical framework centered on the mental lexicon^57–62^. This interface binds acoustic structure to abstract meaning, creating a competitive environment where words can be confused based on either phonological similarity (e.g., son vs. sun) or semantic relatedness (e.g., son vs. nephew). By deploying an automatic phonology–semantics confusion paradigm, we can isolate the specific transformation stages at which sound is progressively mapped onto meaning.

In this study, we systematically aligned the internal processing of twelve state-of-the-art SLMs with human stereo-electroencephalography (sEEG) data, revealing two distinct P2S transfer mechanisms in the human brain: a global cross-regional hierarchy and a local sequential transformation within specific neuronal populations. While most SLMs successfully replicate the global hierarchy, we report a critical dissociation: this macro-scale similarity fails to predict model performance. Instead, functional speech comprehension depends on replicating the local sequential P2S dynamics. Causal perturbation experiments validate this finding: lesioning local P2S-transfer units impaired performance, while targeted activation steering improved performance. These results suggest that machine speech comprehension requires not merely structural depth, but precise sequential alignment with the biological mechanism of P2S transfer.

## Results

To compare P2S transfer between the human brain and SLMs, we analyzed their representational dynamics during speech comprehension using a parallel experimental framework (Fig. 1). **In the human brain experiment,** we recorded sEEG from 17 epilepsy patients (8 female) while they passively listened to Mandarin disyllabic words (Extended Data Fig. 1, Supplementary Table 1). Neural responses were segmented into 16 temporal sequence windows for sequence-resolved analysis (Fig. 1b). The stimulus set comprised 52 words from two semantic categories (26 edible, 26 inedible), organized into 26 phonologically similar pairs that shared the first syllable (e.g., *cǎo méi* “strawberry” vs. *cǎo píng* “lawn”; Supplementary Table 2). **In the SLM experiment,** we presented the same speech stimuli to 12 models (Table 1) and extracted sequence-resolved activations from all model neurons (Fig. 1b). **In the representational alignment experiment,** we systematically quantified brain–model alignment across three dimensions: hierarchical organization (brain regions vs. model layers), sequential dynamics (temporal sequence windows vs. sequence segments), and their relationship to model performance (Fig. 3).

**Fig. 1.**
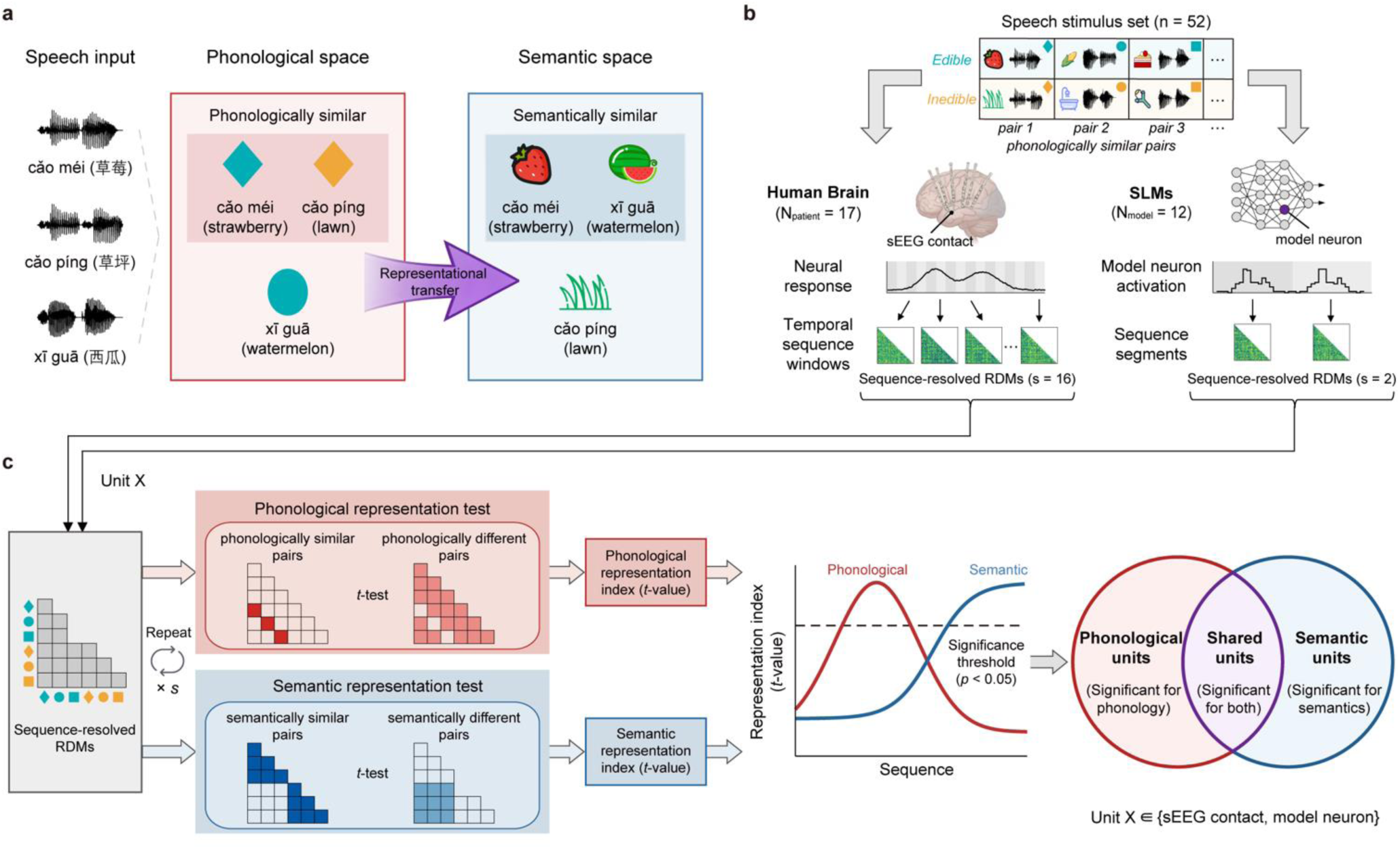
| Framework for comparing phonological-to-semantic representations in the human brain and speech language models (SLMs). **a**, Schematic of phonological-to-semantic representational transfer. Speech input (*left*) is first encoded into a phonological representational space (*middle*), in which phonologically similar words cluster together, and is then mapped via representational transfer into a semantic representational space (*right*), in which semantically similar words cluster together. **b,** Parallel experimental procedures in the human brain and SLMs. **Speech stimulus set:** 52 Mandarin disyllabic words from two semantic categories (26 edible and 26 inedible), organized into 26 phonologically similar pairs that shared the first syllable but differed in semantic category (e.g., *cǎo méi* “strawberry” vs. *cǎo píng* “lawn”). **Human brain experiment** (N_patient_ = 17; *left*): Stereo-electroencephalography (sEEG) recordings during passive listening were segmented into 16 temporal sequence windows, and pairwise decoding accuracies were used to construct 52 × 52 neural representational dissimilarity matrices (RDMs) for each sequence window and sEEG contact. **SLM experiment** (N_model_ = 12; *right*): Model activations were split into two sequence segments per word (*segment 1* and *segment 2*), and pairwise cosine distances were used to generate 52 × 52 model RDMs for each sequence segment and model neuron. **c,** Classification of representational preference for each unit. For each unit X (sEEG contact or model neuron), phonological and semantic representation indices were computed per sequence window or segment from the corresponding RDM by contrasting different versus similar word-pair distances (one-sided two-sample *t*-test). A window or segment was labeled phonological or semantic when the corresponding test was significant (*P* < 0.05). Units were categorized as phonological if only phonological tests were significant across windows/segments, semantic if only semantic tests were significant, or shared if both were significant.

**Fig. 2.**
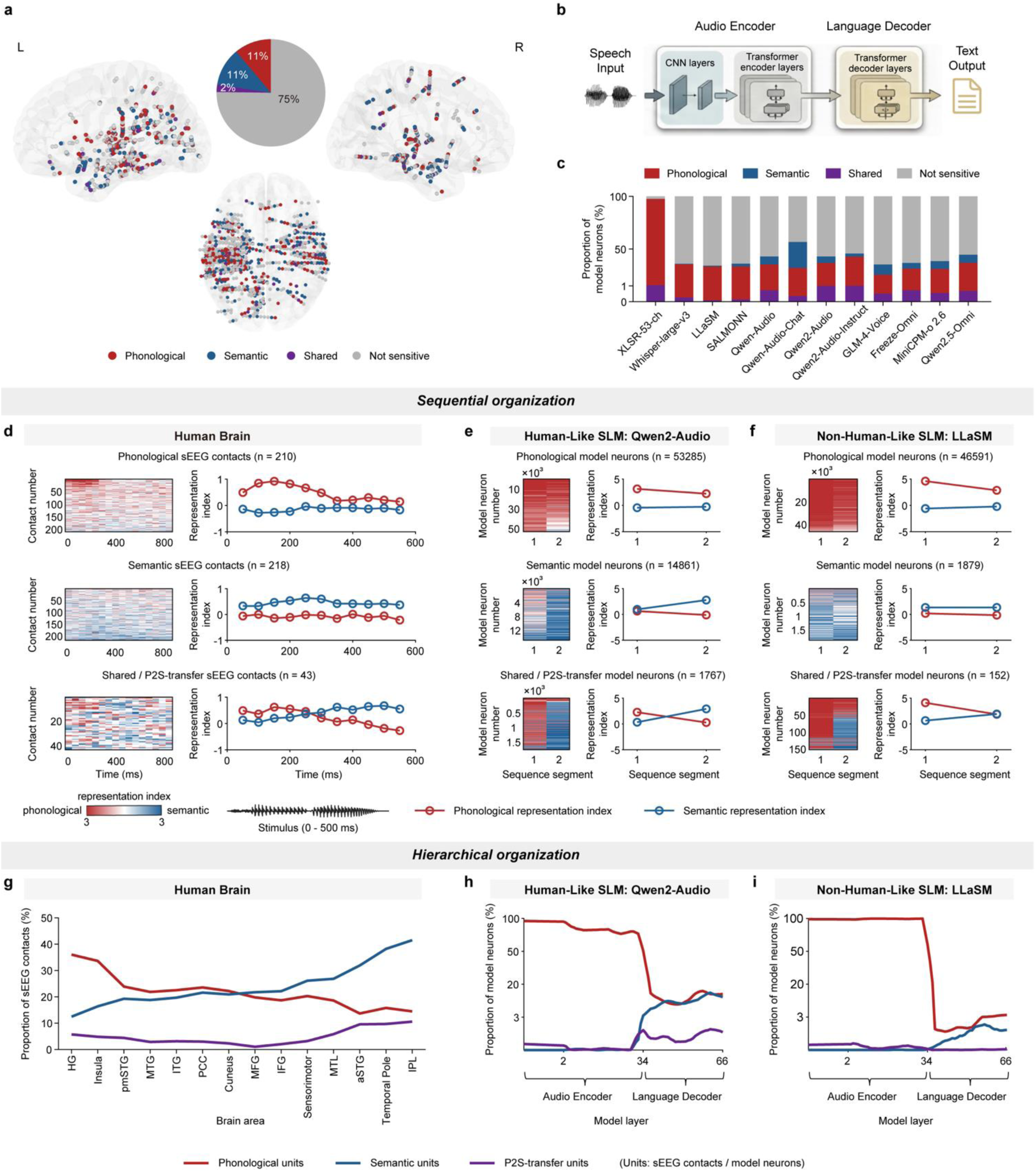
| Sequential and hierarchical organization of phonological, semantic and P2S-transfer representations in the human brain and SLMs. **a**, Spatial distribution of sEEG contacts classified as phonological (*red*), semantic (*blue*), shared (*purple*), or not sensitive (*gray*) in the left (L) and right (R) hemispheres, with overall proportions shown in the pie chart. **b,** Schematic of a generic SLM architecture, showing an audio encoder (CNN layers and transformer encoder layers) followed by a transformer-based language decoder. **c,** Proportions of model neurons in each class across SLMs (colors as in a). **d,** Sequential organization in the human brain. For phonological, semantic, and shared sEEG contacts (*top* to *bottom*), heatmaps show the dominant representation index (defined as the larger of the phonological and semantic index) across successive temporal sequence windows for individual contacts, sorted by dominant representation index. Line plots show mean phonological (*red*) and semantic (blue) representation indices over the temporal sequence. Notably, shared contacts exhibit a transition from phonological to semantic representation along the temporal sequence; these are hereafter termed P2S-transfer contacts. **e,** Sequential organization in a human-like SLM (Qwen2-Audio). For phonological, semantic, and shared model neurons (*top* to *bottom*), heatmaps show sequence-resolved dominant representation index for sequence segments 1 and 2, and line plots show mean phonological and semantic representation indices across segments. Similar to the human brain, these shared model neurons also exhibit a transition from phonological to semantic representation across sequence segments and are thus termed P2S-transfer neurons. **f,** Sequential organization in a non-human-like SLM (LLaSM), shown as in **e. g,** Hierarchical organization in the human brain. Proportions of phonological (*red*), semantic (*blue*), and P2S-transfer (*purple*) sEEG contacts across brain areas (HG, Heschl’s gyrus; pmSTG, posterior/middle superior temporal gyrus; MTG, middle temporal gyrus; ITG, inferior temporal gyrus; PCC, posterior cingulate cortex; MFG, middle frontal gyrus; IFG, inferior frontal gyrus; MTL, medial temporal lobe; IPL, inferior parietal lobule). **h,** Hierarchical organization in a human-like SLM (Qwen2-Audio). Proportions of phonological, semantic, and P2S-transfer model neurons across model layers, spanning the audio encoder and language decoder (module boundaries indicated). **i,** Hierarchical organization in a non-human-like SLM (LLaSM), shown as in **h**.

**Fig. 3.**
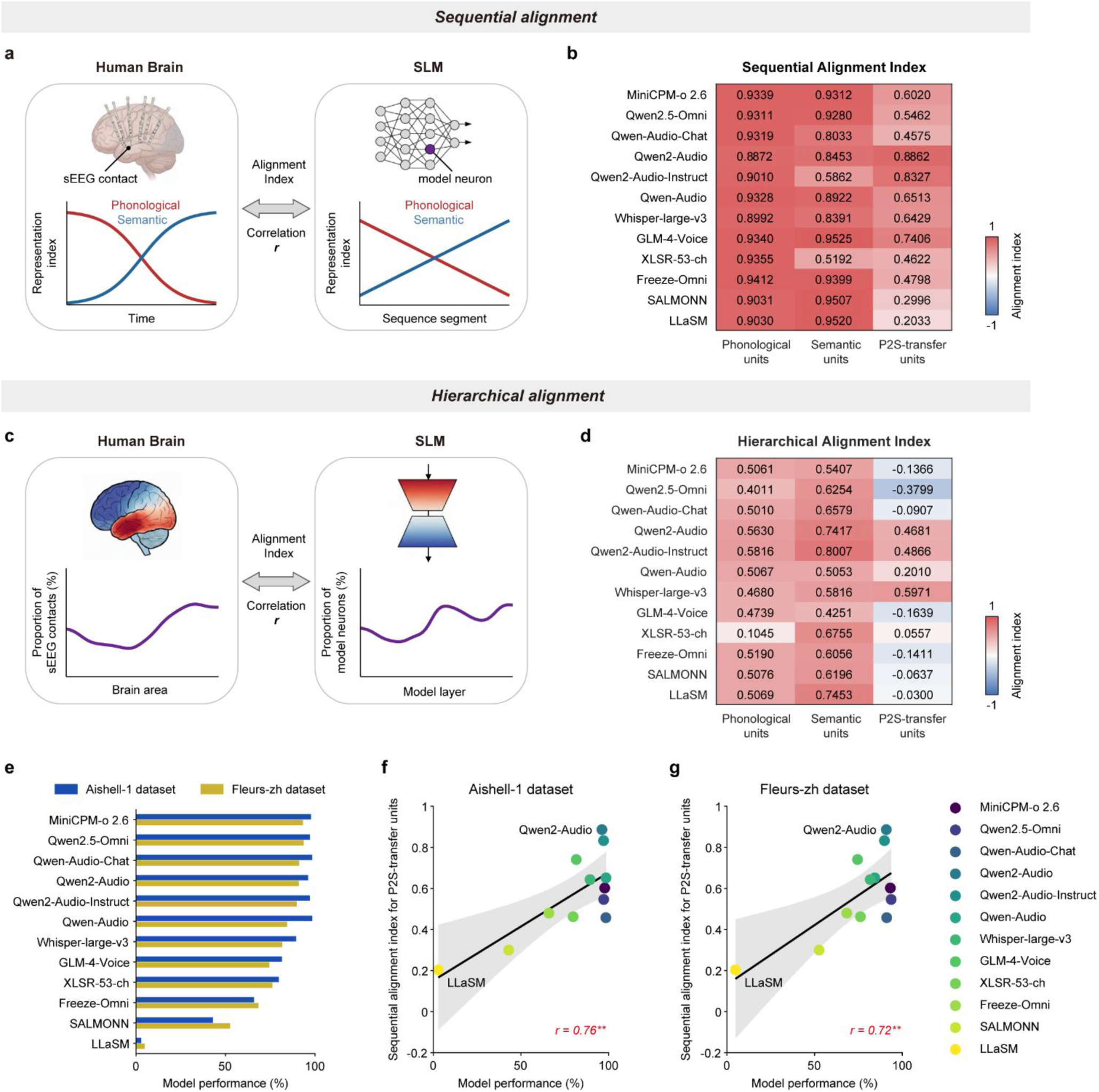
| Sequential and hierarchical brain–SLM alignment identifies P2S-transfer units predictive of model performance. **a**, *Sequential alignment.* For each unit class (phonological, semantic, and P2S-transfer), a sequential alignment index was defined as the Pearson correlation (*r*) between the sequential dynamics of phonological and semantic representation indices in the human brain (across temporal sequence windows in sEEG contacts) and in SLMs (across sequence segments in model neurons). **b,** Sequential alignment indices across SLMs for phonological, semantic, and P2S-transfer units. Models are ranked by mean performance (averaged across two benchmarks; see **e**) (highest to lowest); color scale indicates alignment strength. **c,** *Hierarchical alignment.* For each unit class, a hierarchical alignment index was defined as the Pearson correlation (*r*) between the distribution of unit proportions across brain areas in humans and across model layers in SLMs. **d,** Hierarchical alignment indices across SLMs for phonological, semantic, and P2S-transfer units. Models are ranked by mean performance (as in **b**) (highest to lowest); color scale indicates alignment strength. **e,** Performance of evaluated SLMs on two Mandarin speech recognition benchmarks (Aishell-1 and Fleurs-zh). Mean performance across these benchmarks is used to rank models in **b** and **d**. **f, g,** Relationship between SLM performance and sequential alignment index for P2S-transfer units on the Aishell-1 dataset (f) and the Fleurs-zh dataset (g). Each point represents one SLM, color-coded by model identity (see legend). Linear regression line with shaded 95% confidence interval is shown; Pearson correlation coefficient (*r*) indicates significant positive correlation (\*\**P <* 0.01).

**Table 1.**
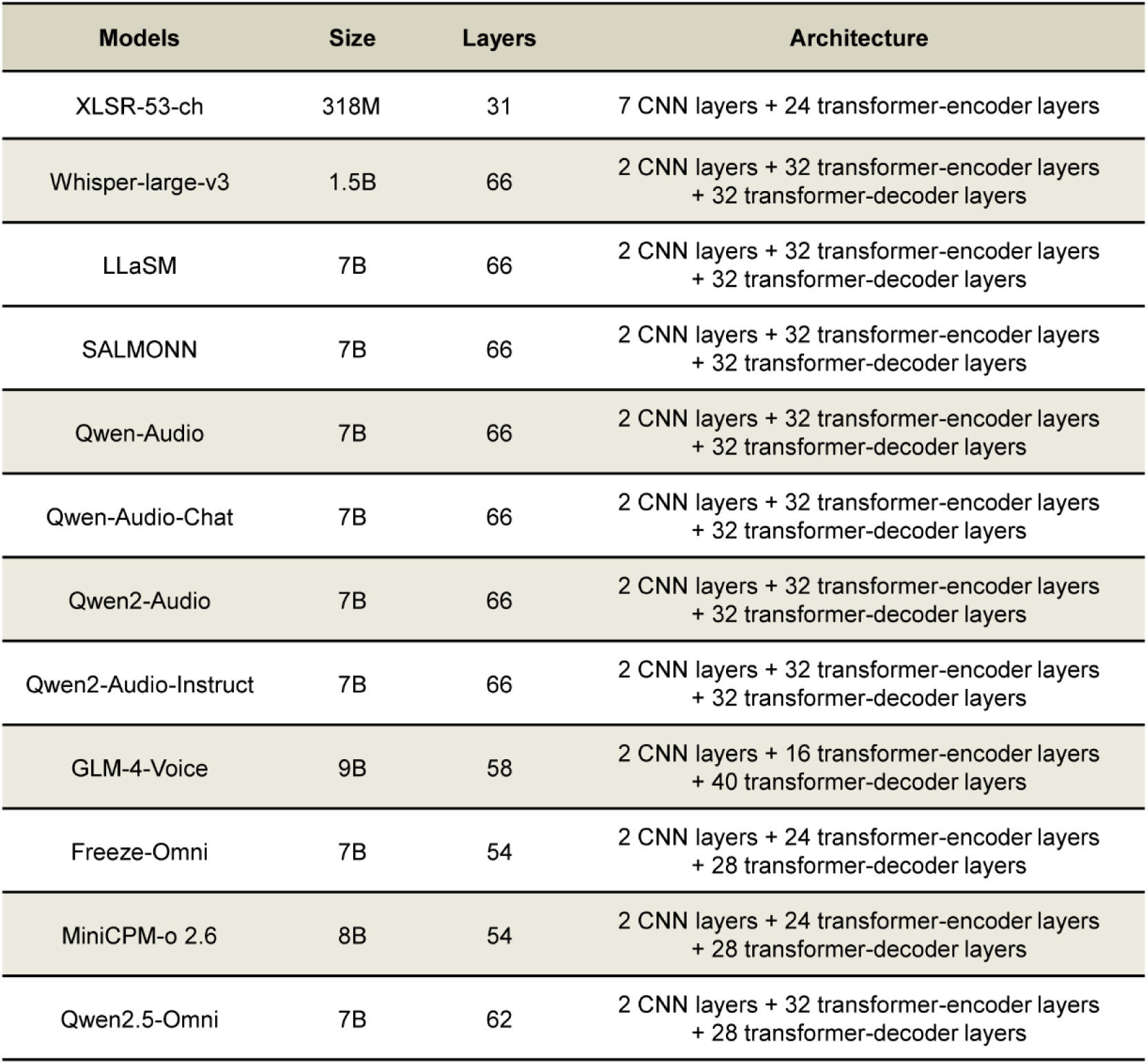
| Summary of speech language model parameters and architectures.

| Models | Size | Layers | Architecture |
| --- | --- | --- | --- |
| XLSR-53-ch | 318M | 31 | 7 CNN layers + 24 transformer-encoder layers |
| Whisper-large-v3 | 1.5B | 66 | 2 CNN layers + 32 transformer-encoder layers<br>+ 32 transformer-decoder layers |
| LLaSM | 7B | 66 | 2 CNN layers + 32 transformer-encoder layers<br>+ 32 transformer-decoder layers |
| SALMONN | 7B | 66 | 2 CNN layers + 32 transformer-encoder layers<br>+ 32 transformer-decoder layers |
| Qwen-Audio | 7B | 66 | 2 CNN layers + 32 transformer-encoder layers<br>+ 32 transformer-decoder layers |
| Qwen-Audio-Chat | 7B | 66 | 2 CNN layers + 32 transformer-encoder layers<br>+ 32 transformer-decoder layers |
| Qwen2-Audio | 7B | 66 | 2 CNN layers + 32 transformer-encoder layers<br>+ 32 transformer-decoder layers |
| Qwen2-Audio-Instruct | 7B | 66 | 2 CNN layers + 32 transformer-encoder layers<br>+ 32 transformer-decoder layers |
| GLM-4-Voice | 9B | 58 | 2 CNN layers + 16 transformer-encoder layers<br>+ 40 transformer-decoder layers |
| Freeze-Omni | 7B | 54 | 2 CNN layers + 24 transformer-encoder layers<br>+ 28 transformer-decoder layers |
| MiniCPM-o 2.6 | 8B | 54 | 2 CNN layers + 24 transformer-encoder layers<br>+ 28 transformer-decoder layers |
| Qwen2.5-Omni | 7B | 62 | 2 CNN layers + 32 transformer-encoder layers<br>+ 28 transformer-decoder layers |

To establish a commensurable representational space between human and SLM data, we implemented a unified unit-level framework treating sEEG contacts and model neurons as equivalent computational units (Fig. 1c). For each unit, representational dissimilarity matrices (RDMs) were constructed from pairwise word comparisons across sequence windows (human) or segments (SLMs). Units were classified as phonological, semantic, or shared based on the significance of representational sensitivity tests within each dimension (see Methods). Representation strength was quantified by *t*-values and defined as phonological and semantic representation indices. Shared units, which exhibited a sequential transition from phonological to semantic representation, are hereafter termed P2S-transfer units.

### Dual neural mechanisms govern human phonology-to-semantics transfer

We identified 962 speech-responsive sEEG contacts from 1,909 recording sites across 17 patients (see Methods; Extended Data Fig. 1). For each responsive contact, we performed pairwise decoding across temporal sequence windows and constructed a 52 × 52 RDM per window from decoding accuracies (see Methods). We then quantified phonological and semantic representation indices from these RDMs and classified contacts as phonological, semantic, or shared (Fig. 2a). Across the 962 responsive contacts, 210 (21.8%) were classified as phonological, 218 (22.7%) as semantic, and 43 (4.5%) as shared (which exhibited significant representation in both dimensions across the temporal sequence windows) (Fig. 2a, d).

To test whether shared contacts mediate a directional transformation, we aggregated their representation indices across windows. These contacts exhibited a clear transition from phonological to semantic representation: phonological representation peaked early (50–250 ms) and declined as semantic representation emerged later (250–550 ms) (Fig. 2d, bottom). High-γ activity (60–150 Hz) further supported this transition, displaying temporal courses intermediate between purely phonological and purely semantic contacts (Extended Data Fig. 1e–g). Accordingly, we termed these shared contacts P2S-transfer contacts. By contrast, phonological contacts showed strong phonological representation in early temporal sequence windows with virtually no semantic representation (Fig. 2d, top), whereas semantic contacts exhibited sustained semantic representation from mid-to-late windows with virtually no phonological representation (Fig. 2d, middle).

The spatial distribution of these three contact classes revealed a hierarchical gradient (Fig. 2a, g). Phonological contacts were enriched in early auditory regions (Heschl’s gyrus, insula, posterior–middle superior temporal gyrus) and declined toward higher-order language regions, whereas semantic contacts showed the opposite gradient, increasing toward anterior temporal and parietal regions (Fig. 2g; detailed regional distributions in Extended Data Fig. 2). P2S-transfer contacts were distributed at relatively uniform proportions (3–6%) across all regions (Fig. 2g), consistent with a distributed transformation network rather than a localized interface. The overall proportions of the three classes did not differ between hemispheres (*χ*² = 3.136, *P* = 0.208; Extended Data Fig. 2o; detailed regional distributions in Extended Data Fig. 2a–n).

To quantify the temporal precedence of phonological over semantic representations, we estimated encoding latencies for each dimension. Phonological latency was computed from phonological and P2S-transfer contacts (*n* = 253), and semantic latency from semantic and P2S-transfer contacts (*n* = 261). Phonological representations emerged significantly earlier than semantic representations by approximately 40 ms (phonological: 183 ± 8 ms; semantic: 223 ± 9 ms; *t*(437) = −2.35, *P* = 0.019, two-sided *t*-test; Supplementary Fig. 1). These findings reveal that human P2S transfer is governed by dual mechanisms: a sparse yet distributed network of contacts executing rapid local sequential transformations, which is entirely embedded within a broader global cortical hierarchy.

### Human-like dual mechanisms emerge in some artificial speech models

In parallel, we analyzed 12 state-of-the-art, open-source SLMs spanning distinct architectures and training paradigms (Table 1), including speech recognition models (XLSR-53-ch^5^, Whisper-large-v3^6^), audio-language models (LLaSM^7^, SALMONN^8^, Qwen-Audio^9^, Qwen-Audio-Chat^9^, Qwen2-Audio^10^, Qwen2-Audio-Instruct^10^, GLM-4-Voice^12^, Freeze-Omni^11^), and multimodal omni models (MiniCPM-o 2.6^13^, Qwen2.5-Omni^14^). Despite these differences, all SLMs shared a hierarchical architecture with an audio encoder and, in most cases, a transformer-based language decoder (Fig. 2b); XLSR-53-ch was an encoder-only exception. We presented the same 52-word speech stimulus set to each model and extracted sequence-resolved activations from all model neurons (see Methods). To accommodate variable sequence lengths, each word’s activation sequence was bisected into two segments (segment 1 and segment 2), and for each segment we computed pairwise cosine distances across words to construct a 52 × 52 model RDM per model neuron (Fig. 1b).

Applying the same unit-level framework, we quantified phonological and semantic representation indices and classified model neurons as phonological, semantic, or shared. Across the 12 models, the proportions of these classes varied substantially (Fig. 2c), indicating that phonological and semantic selectivity—and their co-occurrence—are not fixed outcomes of speech-language training but depend on model-specific factors.

To determine whether shared model neurons mediate directional P2S transformation, we examined their sequential dynamics in two representative models: Qwen2-Audio, which exhibited human-like P2S patterns, and LLaSM, which showed divergent dynamics. In Qwen2-Audio, we identified 53,285 phonological neurons (30.5%), 14,861 semantic neurons (8.5%), and 1,767 shared neurons (1.0%); in LLaSM, we identified 46,591 phonological (26.7%), 1,879 semantic (1.1%), and 152 shared neurons (0.1%) (Fig. 2e, f).

In Qwen2-Audio, P2S-transfer neurons showed a clear sequential transition: phonological representation dominated in segment 1 and declined as semantic representation strengthened in segment 2, while phonological and semantic neurons retained their respective profiles (Fig. 2e). This was accompanied by a layer-wise hierarchy: phonological neurons dominated early encoder layers, semantic neurons emerged in later decoder layers, and P2S-transfer neurons concentrated at the encoder-decoder boundary before maintaining consistent presence throughout the decoder (Fig. 2h). By contrast, in LLaSM, shared neurons showed minimal modulation across segments, lacking the pronounced P2S transition (Fig. 2f). Although a qualitatively similar layer-wise gradient was present, LLaSM exhibited reduced shared and semantic neuron populations and weaker semantic enrichment in later layers (Fig. 2i).

Extending the analysis across all 12 SLMs revealed that human-like P2S transformation is not uniformly expressed across models. Using the simultaneous presence of human-like hierarchical and sequential organization as the operational criterion, we classified Qwen2-Audio (Fig. 2e, h) and eight other SLMs (Extended Data Fig. 3) as human-like models. These models reproduced the two core features observed in the human brain: a layer-wise global hierarchy of P2S representations, and a local sequential P2S transformation within shared/P2S-transfer neurons. By contrast, LLaSM (Fig. 2f, i), XLSR-53-ch, and SALMONN (Extended Data Fig. 4) were identified as non-human-like.

### Local sequential alignment uniquely dictates artificial speech comprehension

Having established that only a subset of SLMs exhibit human-like P2S transformations, we next asked whether brain–model alignment predicts model performance, and which component of alignment is most informative^40^ (Fig. 3a, c). We quantified brain–model alignment using two complementary approaches, each applied separately to phonological, semantic, and P2S-transfer units. First, sequential alignment was defined as the Pearson correlation between representation indices across temporal sequence windows (human brain) and across sequence segments (SLMs), computed separately for phonological, semantic, and P2S-transfer units (Fig. 3a, b; Supplementary Table 3; see Methods). Second, hierarchical alignment was defined as the correlation between unit proportions across brain regions (human brain) and across model layers (SLMs) for each unit class (Fig. 3c, d; Supplementary Table 3, see Methods).

We evaluated automatic speech recognition (ASR) performance on two Mandarin speech recognition benchmarks, Aishell-1^63^ and Fleurs-zh^64^, following established protocols^6,9,10,14^ (see Methods and Supplementary Table 3; Fig. 3e). Our analysis revealed a striking dissociation: local sequential alignment uniquely dictates model performance. Models whose P2S-transfer units more faithfully reproduced the human sequential trajectory, phonological encoding transitioning to semantic encoding, achieved higher accuracy on both benchmarks (Aishell-1: *r* = 0.76, *P* = 4.03 × 10^-3^; Fleurs-zh: *r* = 0.72, *P* = 7.93 × 10^-3^; Fig. 3f, g). For example, Qwen2-Audio showed high P2S sequential alignment together with strong ASR performance, whereas LLaSM exhibited weak P2S sequential alignment and poor performance (Fig. 3b, e–g). Family-stratified analyses indicated that the pooled P2S-performance correlation was primarily driven by the audio-language model family (Supplementary Fig. 2b). In the other two model families (speech recognition models and multimodal omni models), small sample sizes (*N* = 2 each) limited statistical power to detect family-specific trends (Supplementary Fig. 2a, c).

Crucially, alignment with the brain’s global hierarchical processing for any unit class did not predict model performance (Extended Data Figs. 5 and 6), nor did the sequential alignment of purely phonological or purely semantic units (Extended Data Fig. 5). These findings demonstrate that model efficacy is dictated exclusively by the capacity to replicate the fine-grained local temporal dynamics of biological P2S transfer, while macro-scale structural similarities offer no comparable predictive power.

### Local P2S-transfer model neurons form a causal bottleneck for artificial comprehension

To verify whether these local P2S-transfer model neurons are causally necessary for speech comprehension, we performed targeted lesioning in three representative high-performing SLMs (Qwen-Audio^9^, Qwen2-Audio^10^ and Qwen2-Audio-Instruct^10^). Within each model, we ranked neurons in each functional class by representation indices and progressively lesioned them in blocks of 100 until reaching the smallest class size^54,65,66^ (Fig. 4a; see Methods). After each lesioning step, we re-evaluated ASR performance on the Fleurs-zh benchmark (Fig. 4c).

**Fig. 4.**
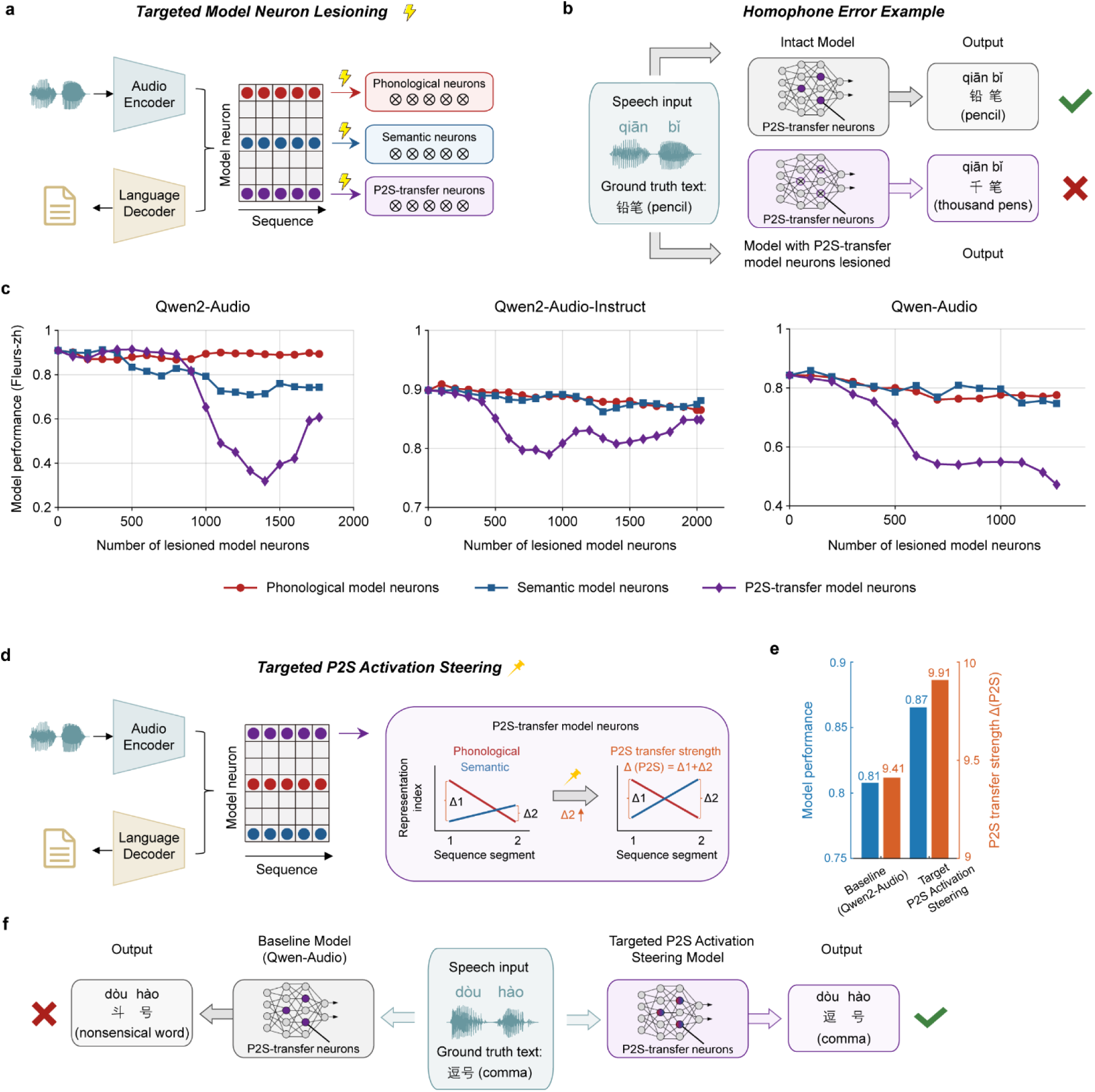
| Bidirectional perturbation of local P2S-transfer model neurons establishes their causal role in phonology-to-semantics transformation within SLMs. **a**, Schematic of targeted model neuron lesioning. For each SLM, model neurons classified as phonological, semantic, or P2S-transfer were selectively lesioned during speech processing to test their causal contributions to phonology-to-semantics transformation. **b,** Example of homophone error caused by disrupting P2S transfer. Given the speech input *qiān bǐ*, the intact model produces the semantically correct output ‘铅笔’ (pencil), whereas the model with lesioned P2S-transfer model neurons produces a homophone error ‘千笔’ (thousand pens), demonstrating preserved phonological recognition but impaired semantic mapping. **c,** Effect of targeted lesioning on model performance. Performance on the Fleurs-zh benchmark is plotted as a function of the number of lesioned neurons for three SLMs (Qwen2-Audio, Qwen2-Audio-Instruct, and Qwen-Audio), with separate curves for lesioning phonological (*red*), semantic (*blue*), and P2S-transfer (*purple*) model neurons. **d,** Schematic of targeted P2S activation steering. During inference, identified P2S-transfer neurons were selectively steered to strengthen their sequential phonology-to-semantics transition. Conceptually, the intervention increases P2S transfer strength by shifting the representation trajectory across sequence segments, quantified as Δ(P2S) = Δ1 + Δ2 (as defined in Methods). Here, Δ1 denotes the phonological-over-semantic dominance of the targeted P2S-transfer neurons in sequence segment 1, whereas Δ2 denotes their semantic-over-phonological dominance in sequence segment 2; larger Δ(P2S) values therefore indicate a stronger sequential phonology-to-semantics transition. **e,** Effect of targeted steering on model performance and P2S-transfer strength in Qwen-Audio. Compared with the baseline model, targeted steering increased both model performance (*blue*) and Δ(P2S) (*orange*). **f,** Example of corrected transcription after P2S activation steering. Given the speech input *dòu hào*, the baseline Qwen-Audio model produced an incorrect nonsensical transcription, whereas the model with steered P2S-transfer neurons generated the correct output ‘逗号’ (comma), illustrating improved semantic mapping after strengthening P2S transfer.

Across all three models, perturbing the local sequential transformation triggered a collapse in model capability. Lesioning P2S-transfer neurons produced disproportionately severe performance deficits compared with lesioning equivalent numbers of phonological or semantic neurons (Fig. 4c). In Qwen2-Audio, lesioning the first 1,000 P2S-transfer neurons reduced accuracy from 95% to 65%, whereas lesioning an equivalent number of purely phonological or semantic neurons caused minor impairments of less than 5 percentage points (Fig. 4c, left). This selective vulnerability was consistent across all three models: the P2S lesioning curve showed the steepest decline and diverged sharply from the shallower curves for phonological and semantic neurons (Fig. 4c). Similar effects were observed in additional human-like SLMs (for example, Qwen-Audio-Chat), with full results reported in Extended Data Figs. 7 and 8.

Qualitative inspection revealed a distinctive error pattern after P2S-transfer lesioning: systematic homophone substitutions (Fig. 4b; Supplementary Table 4). For a speech input with the pronunciation *qiān bǐ*, the intact model produced the semantically appropriate text “铅笔” (pencil), whereas lesioning P2S-transfer neurons yielded the homophonic but semantically inappropriate output “千笔” (thousand pens) (Fig. 4b). These errors indicate preserved access to acoustic-phonological information but impaired recruitment of semantic constraints needed to disambiguate homophones. By contrast, lesioning phonological neurons produced phonetic confusions, whereas lesioning semantic neurons caused broader accuracy decline without a systematic homophone pattern (Supplementary Table 4). These lesioning results definitively establish that local P2S-transfer neurons, despite comprising only ∼1% of the network, form a critical causal bottleneck for advanced artificial speech comprehension.

We then asked whether artificially strengthening local P2S transfer could enhance speech comprehension in SLMs. We introduced a gain-of-function intervention^67^, termed targeted P2S activation steering, in Qwen-Audio, a representative medium-performing human-like SLM, by selectively steering identified P2S-transfer model neurons to enhance their sequential phonology-to-semantics transformation during inference (Fig. 4d). This manipulation increased model performance by enhancing P2S transfer strength (Fig. 4e, f). These results further support a causal link between local P2S-transfer units and speech comprehension in SLMs.

To test cross-linguistic generalizability, we applied the same analysis to an English word dataset (26 edible–inedible pairs matched for first syllable and lexical frequency)^68^. Qwen2-Audio showed a similar sequential and hierarchical organization of P2S-transfer neurons (Extended Data Fig. 9), and targeted lesioning again produced the strongest comprehension impairment relative to phonological or semantic neuron lesioning (Extended Data Fig. 9b). These results indicate that local P2S-transfer dynamics reflect a general computational motif across languages.

### Dynamic geometric reconfiguration underlies local sequential transformations

To characterize how P2S-transfer units transform sound into meaning, we examined the sequential evolution of representational geometry using RDM-based analyses^69^ and multidimensional scaling (MDS)^70^ across temporal sequence windows (human brain) and sequence segments (SLMs)^48^.

In human P2S-transfer contacts, representational geometry shifted systematically: early windows (50–250 ms) showed stronger phonological organization, while later windows (250–450 ms) showed strengthened semantic and weakened phonological organization (Fig. 5a, d). MDS trajectories confirmed progressive reconfiguration from phonological to semantic clustering (Fig. 5g, Supplementary Fig. 4). Phonological and semantic contacts showed stable geometries across windows, confirming this dynamic evolution is specific to P2S-transfer units (Extended Data Fig. 10a, d). In Qwen2-Audio, P2S-transfer neurons showed analogous cross-segment geometric evolution (Fig. 5b, e, h), whereas LLaSM showed an attenuated transition without the pronounced P2S shift (Fig. 5c, f, i). Across all 12 SLMs, human-like models (*N* = 9) exhibited progressive phonological-to-semantic geometric evolution, while non-human-like models (*N* = 3) lacked this signature (Supplementary Figs. 6–17). These findings reveal the computational mechanism underlying P2S transformation: a dynamic reconfiguration of representational space that progressively separates phonological neighbors into distinct semantic categories.

**Fig. 5.**
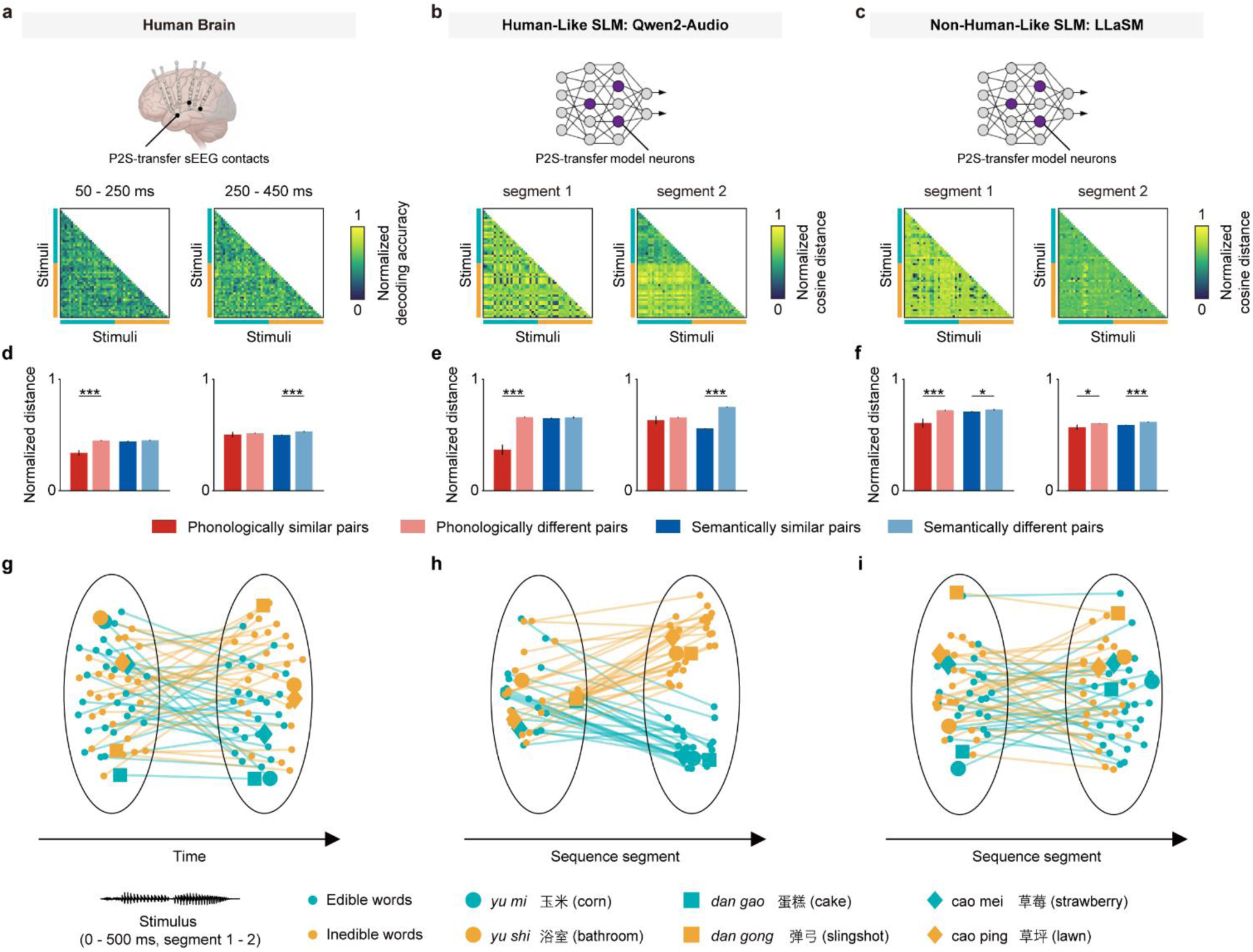
| P2S-transfer units exhibit a structured transformation from phonological to semantic representational geometry in both the human brain and human-like SLMs. **a**, Human brain P2S-transfer sEEG contacts. RDMs (based on pairwise decoding accuracy) are shown for early (50–250 ms) and late (250–450 ms) temporal sequence windows, demonstrating a transition from phonological to semantic representational structure. **b,** Human-like SLM (Qwen2-Audio) P2S-transfer model neurons. Sequence-resolved RDMs (based on cosine distance) are shown for sequence segments 1 and 2. **c,** Non-human-like SLM (LLaSM) P2S-transfer model neurons, shown as in **b**. **d–f,** Quantification of representational distances between word pairs grouped by phonological and semantic similarity. Bars show normalized distances for phonologically similar versus phonologically different pairs and semantically similar versus semantically different pairs (colors as indicated) in the human brain (**d**), Qwen2-Audio (**e**), and LLaSM (**f**). Asterisks denote significance (\**P* < 0.05; \*\*\**P* < 0.001; one-sided two-sample *t*-test). **g–i,** Two-dimensional embeddings via multidimensional scaling (MDS) of the representational transition within P2S-transfer units. Each point represents one word, colored by semantic category. Three phonologically similar word pairs are highlighted with matching distinct shapes to illustrate the phonology-to-semantics transformation; lines connect the same word across temporal sequence windows (**g**) or sequence segments (**h, i**). Trajectories reveal structured phonology-to-semantics reorganization in the human brain and Qwen2-Audio, but attenuated geometric reconfiguration in LLaSM.

## Discussion

The present study reveals that advanced speech comprehension in both biological and artificial systems converges on a shared computational solution: local sequential P2S transformation within a small population of specialized units. Using a phonology–semantics confusion paradigm combined with sEEG recordings and systematic analysis of twelve SLMs, we identified two mechanisms of P2S transfer in the human brain and demonstrated that only local sequential alignment predicts model performance. Bidirectional causal manipulation further confirmed the functional necessity of these units, as their lesioning impaired and their activation steering improved comprehension. The cross-linguistic consistency of this finding suggests it reflects a fundamental rather than language-specific principle. Together, these results bridge cognitive neuroscience and artificial intelligence, establishing local sequential P2S transformation as a core computational principle for speech comprehension.

Our identification of three functionally distinct unit types refines the established hierarchical framework of speech processing. Prior work demonstrates a clear anatomical progression from primary auditory cortex to higher-order language areas^3,4,21,26^, with early regions encoding acoustic-phonetic features^15,19,20,24^ and later regions representing abstract semantic information^21,25,26^. Our spatial analysis corroborates this global hierarchy: phonological units dominate primary auditory regions, semantic units predominate in higher-order areas, and P2S-transfer units are distributed throughout to bridge this processing gap. Remarkably, SLMs spontaneously develop similar hierarchical organizations without explicit neuroscientific constraints. Phonological units dominate early encoder layers and semantic units emerge in later decoder layers, mirroring the cortical progression. In some SLMs, P2S-transfer units concentrate at the encoder-decoder interface, serving as a structural pivot for the shift from phonological to semantic representations^71^. This parallel organization across brains and models suggests that hierarchical processing is a structural necessity for speech comprehension networks.

While global hierarchies provide structural scaffolding, local sequential P2S transformation delivers the critical computational solution for speech comprehension. Phonology alone produces homophone confusions (sun vs. son), while pure semantics conflates acoustically distinct but related words (doctor vs. nurse); efficient comprehension requires simultaneous integration of both dimensions^57,60,61,72,73^. P2S-transfer units accomplish this through rapid local sequential transitions from phonological to semantic representations, enabling resolution of complementary confusions. Causal evidence from our lesioning experiments confirms this mechanism: selective removal of P2S-transfer units produced characteristic homophone substitution errors, demonstrating that intact local sequential transformation is essential for accurate word recognition. These dynamics are consistent with mental lexicon theory^60,61,74,75^, which proposes that word recognition involves multidimensional feature integration. P2S-transfer units serve as the computational interface enabling this process, dynamically weighting phonological and semantic evidence as acoustic input unfolds over time^72,76^. Their consistency across high-performing SLMs establishes local sequential P2S transformation as a fundamental computational principle.

The relationship between brain–model alignment and behavioral performance is a central question in computational neuroscience. Prior work in vision, language, and auditory processing consistently shows that greater brain-like alignment predicts better model performance^28,31,32,44,52,53,69,77^. However, which specific aspects of alignment most strongly predict functional outcomes has remained unclear. Building on recent work examining hierarchical alignment^40^, we systematically compared global hierarchical and local sequential alignment indices across SLMs. This revealed a striking dissociation: only local sequential alignment of P2S-transfer units significantly predicted model performance, while global hierarchical alignment, despite showing robust brain–model correspondence, showed no significant correlation with functional outcomes. This finding aligns with evidence from vision that accurate brain prediction does not require spatial compositional hierarchies^78^, and demonstrates that functional speech comprehension critically depends on implementing the precise local sequential dynamics of P2S transformation.

Applying neuroscience methodologies to artificial neural networks has proven powerful for uncovering computational principles, from vision model units analogous to visual cortex neurons^79,80^, to language model neurons encoding specific linguistic features^81–84^. Recent advances have further introduced causal interventions: targeted lesioning of model units reveals specific functional deficits^65,66,85,86^. Our study extends this framework by combining unit-level representational geometry with bidirectional causal validation, revealing a previously unidentified computational bottleneck in SLMs. Local P2S-transfer units, characterized by a sequential phonological-to-semantic transition, emerge spontaneously across diverse architectures. Despite comprising approximately 1% of all units, their lesioning caused catastrophic comprehension degradation, while targeted activation steering of the same units improved performance, demonstrating that precise manipulation of these local units selectively governs model behavior.

In sum, our findings reveal that local sequential P2S transformation represents a universal computational principle shared across biological and artificial intelligence. The critical finding that local dynamics, rather than global organization, uniquely predict functional performance challenges conventional wisdom and establishes a new paradigm for evaluating speech comprehension systems. These results illuminate deep computational constraints governing intelligent language processing, with broad implications for both cognitive neuroscience and artificial intelligence.

## Methods

### Human brain

#### Subjects

Seventeen human patients (9 males, all right-handed, all native-Mandarin speakers, 13 to 42 years old) with drug-resistant epilepsy who underwent invasive sEEG monitoring for potential surgical treatments at the Sanbo Brain Hospital of Capital Medical University (Beijing, China) were included in the study. All patients had self-reported normal hearing and provided written informed consent for their participation. The experimental procedure was approved by the Ethics Committee of the Sanbo Brain Hospital of Capital Medical University and the Human Subject Review Committee of Peking University. Demographic information and implantation details are listed in Supplementary Table 1.

#### Experiment and stimuli

We used 52 Mandarin disyllabic speech stimuli recorded from a female native speaker (average fundamental frequency, 254 Hz). The speech stimuli were equally divided into two semantic categories (26 edible, 26 inedible) and organized into 26 phonologically matched pairs (Supplementary Table 2). Within each pair, items shared the same initial syllable (phonologically matched) but differed in semantic category, such as *cǎo méi* (strawberry; edible) versus *cǎo píng* (lawn; inedible). Mean word frequency was matched between the two semantic categories^87^.

Stimuli were delivered binaurally via insert earphones (ER-3, Etymotic Research) connected to a calibrated sound card (SB X-Fi Surround 5.1 Pro, Creative Technology). Intensity was adjusted to ∼70 dB SPL in the ear canal, as verified with a precision sound level meter (AUDit, Larson Davis System 824).

Participants were instructed to passively listen to the sequence of the speech stimuli (Extended Data Fig. 1). The inter-stimulus interval varied randomly between 2 and 4 s. The experiment consisted of four blocks, each containing 156 randomized trials, yielding a total of 624 trials with each stimulus repeated 12 times. Block order was counterbalanced across participants. Recordings were performed while patients were awake and seizure-free for at least 4 hours prior to the experiment.

#### Localization of sEEG electrode contacts

All patients were implanted with stereo-electrodes (Huake Hengsheng Medical Technology Co. Ltd., Beijing, China). Each stereo-electrode (0.8 mm diameter) has 8–16 recording sites (2 mm length, spaced 3.5 mm apart). All electrode implantations were determined solely based on clinical reasons.

To localize the electrode contacts, we integrated the anatomic information of the brain from preoperative magnetic resonance imaging (MRI) and the position information of the electrodes from postoperative computed tomography (CT). For each patient, we first co-registered the post-implant CT with the pre-implant anatomic T1-weighted MRI using the SPM12 software^88^ (https://www.fil.ion.ucl.ac.uk/spm/software/spm12/). Next, we identified electrode traces in the aligned CT images and calculated the coordinates of contacts using the Brainstorm toolbox^89^ (http://neuroimage.usc.edu/brainstorm). To assign the anatomic label to each contact, we performed subcortical and cortical segmentations based on individual preoperative T1 MRI using FreeSurfer version 6.0^90^ and registered the data to the University of Southern California (USC) Brain atlas^91^. To identify cortical contacts, we projected each contact to the nearest vertex on the individual cortical surface using the MATLAB function *dsearchn*, and assigned the contact to a cortical area based on the projected vertex. To identify subcortical contacts, we used the “*iEEG atlas labels*” function in Brainstorm to localize electrode contacts in subcortical structures such as the amygdala and hippocampus. For illustration purposes, the coordinates of contacts were normalized to the MNI space and visualized on the template brain *ICBM152_2023b*.

#### Electrophysiological recording

Intracranial electrophysiological signals from each contact were amplified using a Nicolet video-EEG monitoring system (Thermo Nicolet Corp., USA) without any online filtering. sEEG data at each electrode contact site were sampled at 512 Hz for all patients. Both the reference and ground electrodes were placed on the forehead of the patients. The electrode impedances were kept below 50 kΩ throughout the recording. All further processing was performed offline.

#### Identifying speech-responsive sEEG contacts

We defined speech-responsive sEEG contacts based on significant high-γ (60–150 Hz) activity evoked by speech stimuli. All the intracranial sEEG signals were pre-processed using the EEGLAB toolbox^92^ in the MATLAB environment. To extract high-γ band amplitude, we first filtered the continuous sEEG signals with a zero-phase finite impulse response (FIR) band-pass filter (“*pop_eegfiltnew*” function from the EEGLAB toolbox). We then obtained the amplitude envelope of the high-γ band signals using the Hilbert transform^93^. The high-γ amplitude signals were segmented into epochs from −200 to 1200 ms around sound onset and baseline-corrected using the pre-stimulus interval (−200 to 0 ms). Epochs containing interictal epileptic spikes or recording artifacts were visually inspected and excluded from the analysis^94^.

Speech-responsive sites were identified by comparing high-γ activity during the post-stimulus interval (0–600 ms) with that during the pre-stimulus baseline interval (−200–0 ms)^95^. Specifically, for each sEEG contact, we evaluated responses to each of the 52 speech stimuli individually. Cohen’s *d* effect size was calculated at each time point using trial-wise paired comparisons between response signals and baseline means. A contact was considered responsive to a given speech stimulus if at least 10 consecutive time points (approximately 20 ms at the sampling rate of 512 Hz) exhibited *d* > 0.5 (medium effect size). Finally, an sEEG contact was classified as speech-responsive if it met this criterion for at least five speech stimuli to reduce false positives from stimulus-specific transients.

#### Sequence-resolved representational dissimilarity matrices

For each speech-responsive sEEG contact, we characterized its dynamic representational geometry using a time-resolved spectra-temporal decoding framework^96^. This approach enabled us to track how neural representations evolved across time within each individual sEEG contact. The analysis pipeline consisted of four main steps:

##### Signal preprocessing

The broadband signal (0.5–200 Hz) of the sEEG data was extracted using a zero-phase FIR band-pass filter (“*pop_eegfiltnew*” function from the EEGLAB toolbox) and then segmented into epochs from −200 to 1200 ms relative to stimulus onset^97^. Epochs containing interictal epileptic spikes or recording artifacts (as identified in the speech-responsive contact analysis) were excluded^94^. We then performed time-frequency decomposition using a continuous Morlet wavelet transform (MATLAB function “*cwt*”)^98^. For each contact, trial-wise time-frequency maps spanning 0.5–200 Hz and −200 to 1200 ms relative to stimulus onset were computed and subsequently z-scored for normalization.

##### Sequence-resolved feature extraction

To capture the sequential dynamics of representational geometry, we applied a sliding-window analysis across multiple time epochs. For each contact, we extracted features from 16 consecutive temporal sequence windows (ranging from −50 to 900 ms relative to stimulus onset, 200-ms window, 50-ms step). Within each temporal sequence window, high-γ (60–150 Hz) power was segmented into non-overlapping 100-ms bins, generating 26 spectra-temporal features (13 frequency bins × 2 temporal bins) per window^98^. This sequence-resolved approach allowed us to probe how representational structure changed dynamically following stimulus onset.

##### Classification and cross-validation

For each stimulus pair, dissimilarity was quantified as the cross-validated pairwise classification accuracy using a support vector machine^96,99^ (SVM; LIBSVM implementation). Trials were randomly partitioned into four splits (three trials per split), with one split used for testing and three for training at each iteration. Binary decoding was performed for all 1,326 (*C*^2^) pairwise stimulus comparisons. The SVM employed an RBF kernel, with hyperparameters (Gamma and C) optimized via grid search over exponentially scaled values (2⁻¹⁰ to 2¹⁰, step 2⁰·⁵) using nested four-fold cross-validation^97,98,100^. This procedure was repeated 10 times with different random splits, and accuracies were averaged.

##### Dynamic representational dissimilarity matrix construction

For each sEEG contact, this procedure generated 16 sequence-resolved representational dissimilarity matrices (RDMs). This time series of RDMs captured the evolving representational geometry of speech-evoked high-γ activity, revealing how neural similarity structures transformed dynamically across the post-stimulus period within each electrode.

### Speech language models

We analyzed 12 pretrained speech language models (SLMs) (Table 1), including speech recognition models (XLSR-53-ch^5^, Whisper-large-v3^6^), audio-language models (LLaSM^7^, SALMONN^8^, Qwen-Audio^9^, Qwen-Audio-Chat^9^, Qwen2-Audio^10^, Qwen2-Audio-Instruct^10^, GLM-4-Voice^12^, Freeze-Omni^11^), and multimodal omni models (MiniCPM-o 2.6^13^, Qwen2.5-Omni^14^). The selected SLMs spanned a broad range of open-source architectures, from 318M to 9B parameters and from 31 to 66 layers.

#### Extracting model unit activations

To extract the activations of the SLMs for each speech stimulus, we fed the same 52 speech stimuli to the models and recorded layer-wise activations using PyTorch forward hooks. The evaluated SLMs broadly followed a hierarchical speech-processing pipeline, with an audio encoder front-end and, in most models, a transformer-based language decoder for text generation (Fig. 2b); XLSR-53-ch is an encoder-only exception lacking a language decoder. We applied a unified approach to extract model unit (“model neuron”) activations from each architectural component where present.

For convolutional layers in audio encoders, we registered hooks to capture their output tensors. Individual channels were treated as model neurons, and each neuron’s complete temporal response was preserved across all time steps^101^. For convolutional layers with output shape (*C*, *T*), where *C* represents the number of channels and *T* denotes the temporal sequence length, we obtained *C* model neurons, each with a *T*-dimensional temporal response profile representing the neuron’s activation pattern over time.

For transformer encoder layers in audio encoders, we extracted hidden states from each transformer encoder block and defined individual hidden dimensions as model neurons^86,102^. For each encoder layer with output shape (*T*, *D*), where *T* represents the temporal sequence length and *D* denotes the hidden dimension, we obtained *D* model neurons, each with a *T*-dimensional temporal response.

For transformer decoder layers in language models, we extracted the hidden states from each transformer decoder block, defining individual hidden dimensions as model neurons^86,102^. To isolate model responses to the speech input, we removed prompt-related activations by excluding all activations preceding the first token generated in response to the speech input. For each decoder layer with output shape (*M*, *D*), where *M* represents the number of generated tokens and *D* denotes the hidden dimension, we obtained *D* model neurons, each with an *M*-dimensional temporal response.

#### Sequence-resolved representational dissimilarity matrix

##### Stimuli and models

All 12 models were presented with 52 Mandarin disyllabic speech items (identical to those used in the human experiment). For each item, hidden activations were extracted from the target model units (“model neurons”) across temporal steps or generated tokens, depending on the model architecture.

##### Activation extraction (sequence-resolved)

For each model neuron, activations were recorded sequentially across time steps (for audio processing layers) or tokens (for language generation layers). To accommodate differences in sequence length among words, each word’s sequence was bisected into two contiguous segments, denoted segment 1 and segment 2. All subsequent analyses were conducted separately for segment 1 and segment 2. No additional normalization was applied beyond the model’s native scaling unless otherwise specified.

##### Pairwise distance computation

Within each sequence segment, we characterized representational geometry by computing pairwise cosine distances between speech-evoked activation vectors^103–105^. For words *i* and *j*, with activation vectors *x*_*i*_ and *x*_*j*_ from the same model neuron and sequence segment, cosine distance was defined as:

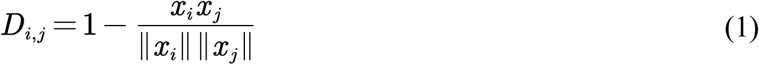

yielding values in [0, 2]. For all 52 stimuli, this resulted in 1,326 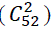 pairwise stimulus comparisons per model neuron per sequence segment.

##### RDM construction

Distances were assembled into an RDM for each model neuron and each sequence segment. Each model neuron thus yielded two RDMs, one for segment 1 and one for segment 2.

#### Assessment of model performance

We evaluated the speech recognition capability of each SLM using a standard automatic speech recognition (ASR) task^6,9,10,14^. The models were tested on two publicly available Mandarin speech datasets: (1) Aishell-1^63^ (a large-scale read-speech corpus containing approximately 178 hours of speech from 400 speakers), and (2) Fleurs-zh^64^ (the Mandarin Chinese subset of the Fleurs multilingual benchmark).

Each model was provided with the raw speech audio as input and generated text transcriptions using its built-in speech recognition interface. Decoding settings followed the default inference interface of each model unless otherwise specified. Model outputs were compared to ground-truth transcripts at the character level, consistent with standard Mandarin ASR evaluation practices. The evaluation metric was Recognition Accuracy, defined as one minus the character error rate (CER):

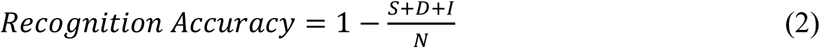

where *S, D*, and *I* denote the numbers of *substitution*, *deletion*, and *insertion* errors, respectively, and *N* is the total number of characters in the reference transcription. To avoid negative per-utterance scores, Recognition Accuracy was clipped to the range [0, 1] for each utterance and then averaged across the dataset. Higher values indicate better model performance (Supplementary Table 3).

### Brain–model alignment

#### Defining three classes of units

In this study, a unit refers to an sEEG contact in the human brain or a model neuron in the SLMs. To enable direct comparability between biological and artificial systems, we applied an identical representational classification procedure to the RDMs derived from each unit (Fig. 1c).

For each RDM, we computed representational distances between word pairs that were either phonologically different or phonologically similar, and likewise between semantically different and semantically similar pairs. Distances for the two conditions were compared using one-sided two-sample *t*-tests (*P* < 0.05) to assess whether different pairs were represented farther apart in the unit’s representational space. Specifically, for phonological representation:

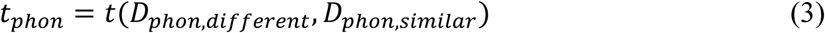

and for semantic representation:

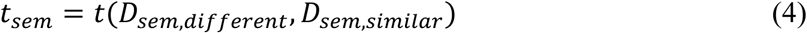

Where *D_phon,different_* and *D_phon,similar_* denote the sets of distances between phonologically different and similar word pairs, respectively; *D_sem,different_* and *D_sem,similar_* denote the corresponding semantic distances. Positive *t*-values indicate stronger representational separation for different pairs compared to similar pairs. The resulting *t*-values were defined as the phonological and semantic representation indices for each RDM.

Each unit yielded multiple RDMs corresponding to temporal sequence windows (for human sEEG contacts) or sequence segments (for SLM model neurons). Units were labeled phonological if at least one RDM showed a significant phonological effect (*P* < 0.05) and none showed a significant semantic effect across temporal windows or sequence segments; units were labeled semantic if at least one RDM showed a significant semantic effect and none showed a significant phonological effect; units were labeled shared if at least one RDM showed a significant effect in each dimension.

This unified classification framework provided a common representational metric for systematically comparing phonological and semantic organization between the human brain and SLMs.

#### Sequence-resolved multidimensional scaling (MDS)

To visualize the representational geometry of both the human brain and SLMs, we applied multidimensional scaling^70^ (MDS) to the sequence-resolved RDMs. This projected the 52 words onto a common two-dimensional space, providing a low-dimensional visualization of the underlying neural or model representations (Fig. 5).

For each unit type in both the human brain and SLMs, RDMs were averaged across all units (sEEG contacts or model neurons) within each temporal sequence window (human brain) or sequence segment (SLMs), yielding a population-level 52 × 52 RDM per window/segment. For each population-level RDM, we computed two-dimensional MDS projections using the *‘mdscale’* function in MATLAB with stress criterion minimization. To enable comparison across temporal windows or sequence segments, all MDS solutions were normalized to unit radius (maximum distance from origin = 1) and sequentially aligned using Procrustes analysis without scaling (allowing only rotation and reflection). This chain alignment procedure ensured that geometric evolution reflected genuine representational changes rather than arbitrary rotations between time points. This approach yielded a sequence-resolved visualization of representational dynamics, enabling direct comparison of how representational geometry evolved over time between biological and artificial systems.

To quantify clustering structure, we computed representational distances between phonologically different versus similar word pairs and between semantically different versus similar pairs within each MDS projection. Phonologically similar pairs were defined as words sharing the first syllable (26 pairs in the stimulus set), whereas semantically similar pairs were defined as words within the same category (edible or inedible). The degree of phonological or semantic clustering was quantified as the difference between mean between-class distances and mean within-class distances (larger values indicate stronger clustering). This metric was computed separately for each temporal sequence window or sequence segment to characterize the evolution of phonological and semantic organization.

#### Estimating hierarchical alignment between the human brain and SLMs

To evaluate the hierarchical alignment between the human brain and SLMs, we computed a hierarchical alignment index for each unit type (phonological, semantic, and P2S-transfer) by comparing their proportional distributions across brain areas and model layers.

For each unit type, the proportional distributions were normalized to a common 0–1 range:

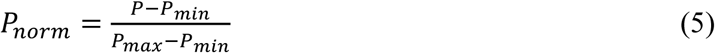

where *P* denotes the proportion of a given unit type within a brain area or model layer, and *P*_*min*_ and *P*_*max*_ are the minimum and maximum proportions across all unit types and all brain areas/layers. We used a shared normalization range across unit types, rather than normalizing each type separately, so that the resulting alignment index retained information about differences in overall unit prevalence as well as differences in hierarchical distribution.

The normalized layer-wise proportions from SLMs were then linearly interpolated to match the number of brain areas. We next computed the Pearson correlation coefficient *r* between the normalized proportional distributions in the human brain and in the SLMs:

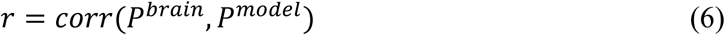

where *P*^*brain*^ and *P*^*model*^ denote the vectors of normalized proportions (*P*_*norm*_) for a given unit type across brain areas and interpolated model layers, respectively. For each unit type, the correlation was calculated across corresponding hierarchical positions. Larger *r* values indicate stronger alignment between the human brain and SLMs in the hierarchical distribution of that unit type.

#### Estimating sequential alignment between the human brain and SLMs

To evaluate the sequential alignment between representational dynamics in the human brain and SLMs, we computed a sequential alignment index for each unit type by comparing the sequential evolution of phonological and semantic representation indices across brain temporal sequence windows and model sequence segments.

For each unit type, the representation indices were first normalized to a common 0–1 range:

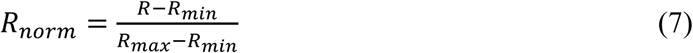

where *R* denotes the phonological or semantic representation index, and *R*_*min*_ and *R*_*max*_ are the minimum and maximum values computed jointly across all unit types and all temporal sequence windows/sequence segments. This shared normalization preserved the relative magnitude of representation indices across unit types.

The normalized sequence-segment trajectories from the SLMs were then linearly interpolated to match the number of temporal sequence windows in the brain. For each unit type, we next computed the Pearson correlation coefficient *r* between the normalized representational trajectories from the brain and model data:

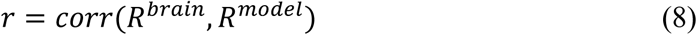

where *R*^*brain*^ and *R*^*model*^ denote the concatenated vectors of normalized phonological and semantic representation indices (*R*_*norm*_) for a given unit type across brain temporal sequence windows and interpolated model sequence segments, respectively. Thus, 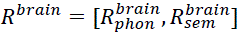 and 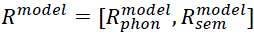. For each unit type, the correlation was calculated across corresponding temporal positions. Larger *r* values indicate stronger sequential alignment in representational dynamics between the human brain and SLMs.

#### In silico lesion experiment

To examine the functional contribution of distinct model neuron types within the SLMs, we conducted a series of *in silico* lesion analyses targeting three predefined categories: phonological, semantic, and P2S-transfer model neurons^65^. The lesion experiment comprised three steps:

##### Baseline performance

We first evaluated each model’s Recognition Accuracy on the Fleurs-zh dataset^64^ to establish the baseline for all subsequent lesion analyses.

##### Lesion procedure

Model neurons were lesioned using a rank-based cumulative deactivation approach^65,66,96^. Specifically, model neurons of each type were first ranked according to their corresponding representation index: phonological, semantic, or a combined index (the sum of both) for P2S-transfer neurons. Lesioning was then applied sequentially in blocks of 100 model neurons, beginning with the highest-ranked ones, until reaching the smallest population size among the three types. During each lesioning step, activations of the targeted model neurons were set to zero throughout the forward pass, effectively removing their contribution to model computations.

##### Evaluation of lesion effects

After each lesioning step, the model’s ASR accuracy on the Fleurs-zh dataset was re-evaluated to quantify the functional impact of progressive deactivation. The resulting performance degradation curves characterized how selectively silencing phonological, semantic, and P2S-transfer model neurons affected overall model performance, thereby providing a direct estimate of the model’s functional dependence on each representational subspace.

#### In silico steering experiment

To further close the loop on our findings and test whether the identified P2S-transfer mechanism can positively inform SLM optimization, we performed an in silico steering experiment in a representative medium-performing model, Qwen-Audio, by selectively modulating the activations of P2S-transfer model neurons during inference^67^. Unlike the lesion analysis, which removed the contribution of targeted neurons, the steering experiment was designed to strengthen the sequential P2S transformation within a predefined P2S subspace while preserving the remaining model computations. All steering analyses were conducted on the Mandarin disyllabic speech stimulus set listed in Supplementary Table 2. The procedure comprised three steps.

##### Baseline pass

We first ran the intact model on the full 52-word stimulus set without intervention to obtain baseline hidden representations for each model neuron. For each intervention layer, these baseline activations were then used to define a steering axis within the subset of direction-consistent P2S-transfer model neurons. Specifically, we computed the mean second-segment activations for edible and inedible words, took the difference between the two category means as the steering direction, and masked this direction so that only dimensions corresponding to the selected P2S-transfer neurons were retained for intervention.

##### Targeted steering of P2S-transfer neurons

Steering was restricted to direction-consistent P2S-transfer model neurons, defined as neurons showing higher phonological than semantic representation index in sequence segment 1 and higher semantic than phonological representation index in sequence segment 2. For neuron *u*, these two directional components were defined as

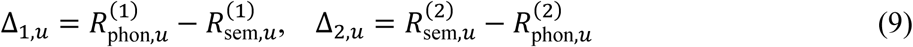

Only neurons satisfying Δ_1,*u*_ > 0, Δ_2,*u*_ > 0 were retained as steering candidates. These neurons were ranked by a consistency score:

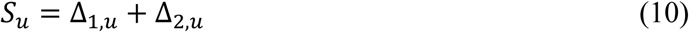

The top 1,000 P2S-transfer neurons were selected as steering targets. During inference, forward hooks were inserted at the selected layers, and intervention was applied only to the second sequence segment of the hidden state. At each position in this segment, the hidden activation was biased along the masked steering direction, and the magnitude of this modulation was controlled by a steering coefficient α. This procedure selectively reinforced the phonology-to-semantics transition within the targeted P2S subspace without globally perturbing the full hidden state.

##### Evaluation of steering effects

After intervention, model performance was re-evaluated on the same 52-word stimulus set across a range of steering strengths. In parallel, we quantified P2S transfer strength as:

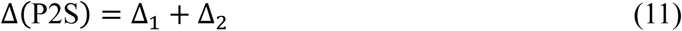

where

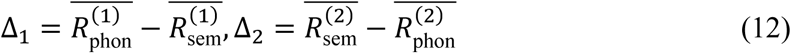

Here, 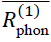 and 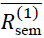 denote the average phonological and semantic representation indices of the targeted neurons in sequence segment 1, and 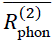 and 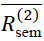 denote the corresponding averages in sequence segment 2. Larger Δ(P2S) values indicate a stronger sequential P2S transition. This allowed us to compare the effects of different steering coefficients on both recognition performance and representational reorganization. Finally, for descriptive reporting, we retained the results obtained with the steering coefficient that produced the largest performance improvement.

## Data and code availability

Processed source data and custom code are openly available at https://github.com/purityss/p2s-slm-brain-alignment. Due to patient privacy and institutional regulations, raw sEEG recordings are restricted but can be made available from the corresponding authors upon reasonable request and completion of a data transfer agreement.

## Acknowledgments

This work was supported by the National Science and Technology Innovation 2030 Major Program (2022ZD0204802) (FF), (2022ZD0204804) (QW), the National Natural Science Foundation of China (T2421004) (FF), (31930053) (FF), (32171039) (QW), and Xiaomi Foundation (QW).

## Extended Data

**Extended Data Fig. 1.**
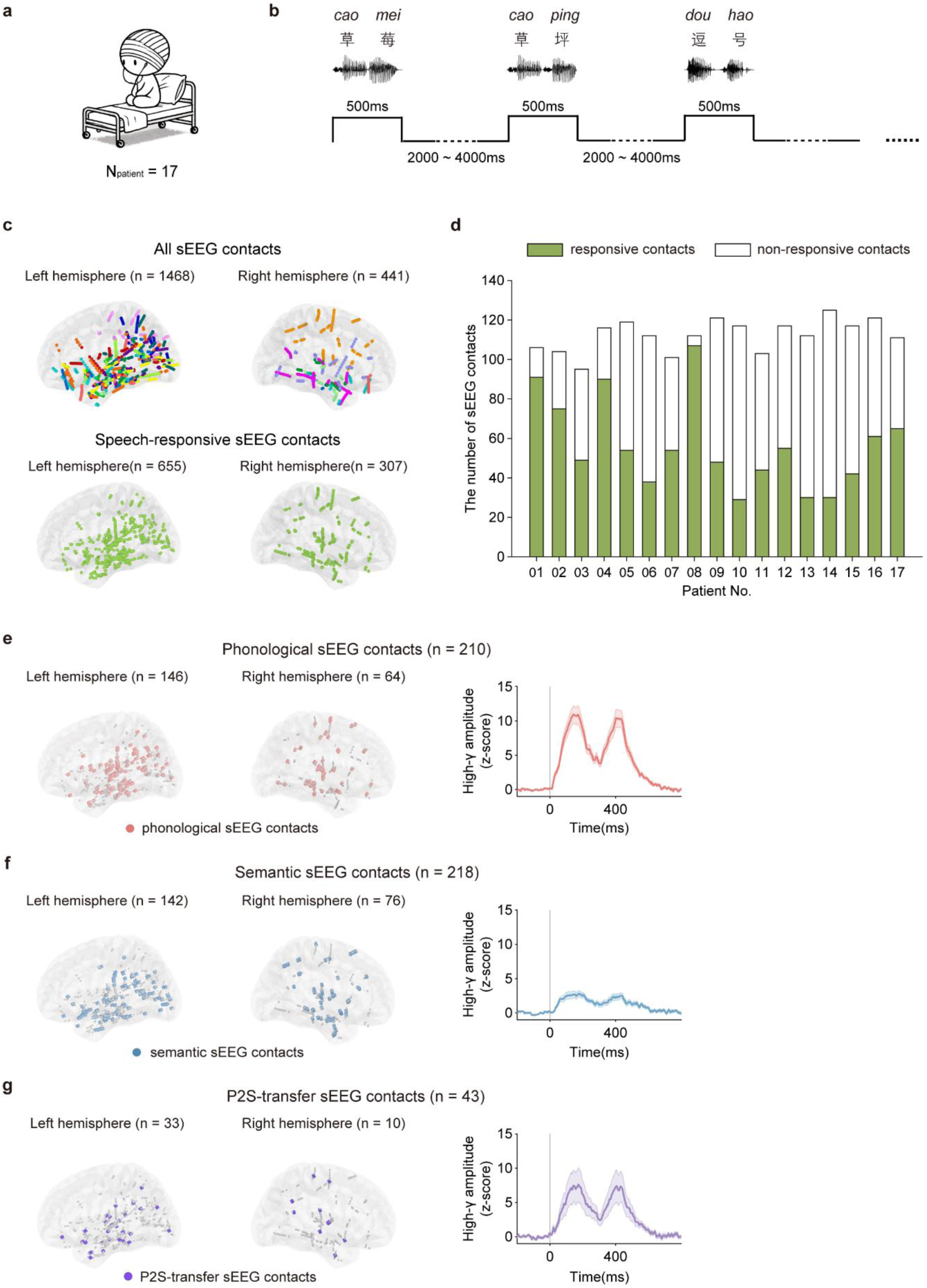
| Experimental paradigm, anatomical localization, and temporal dynamics of speech-responsive sEEG contacts in the human brain. **a**, Schematic illustration of the sEEG recording paradigm. Seventeen epilepsy patients participated in the study. **b,** The experimental procedure. Mandarin disyllabic speech stimuli were presented in a random order. Each stimulus was displayed for 500 ms, followed by an inter-stimulus interval (ISI) randomly jittered between 2000 and 4000 ms (Methods). **c,** Anatomical distribution of all sEEG contacts (*top*) and speech-responsive contacts (*bottom*). Contacts are projected onto the ICBM152 template brain for visualization. **d,** Patient-wise distribution of speech-responsive contacts. Green bars indicate the number of speech-responsive contacts; white bars indicate non-responsive contacts for each patient. **e–g,** Spatial distribution (*left*) and averaged high-γ amplitude (*z*-score) time courses (*right*) for phonological (**e**, *n* = 210), semantic (**f**, *n* = 218), and P2S-transfer (**g**, *n* = 43) sEEG contacts. Each dot represents one contact. Numbers in parentheses indicate contact counts per hemisphere. Shaded areas, SEM.

**Extended Data Fig. 2.**
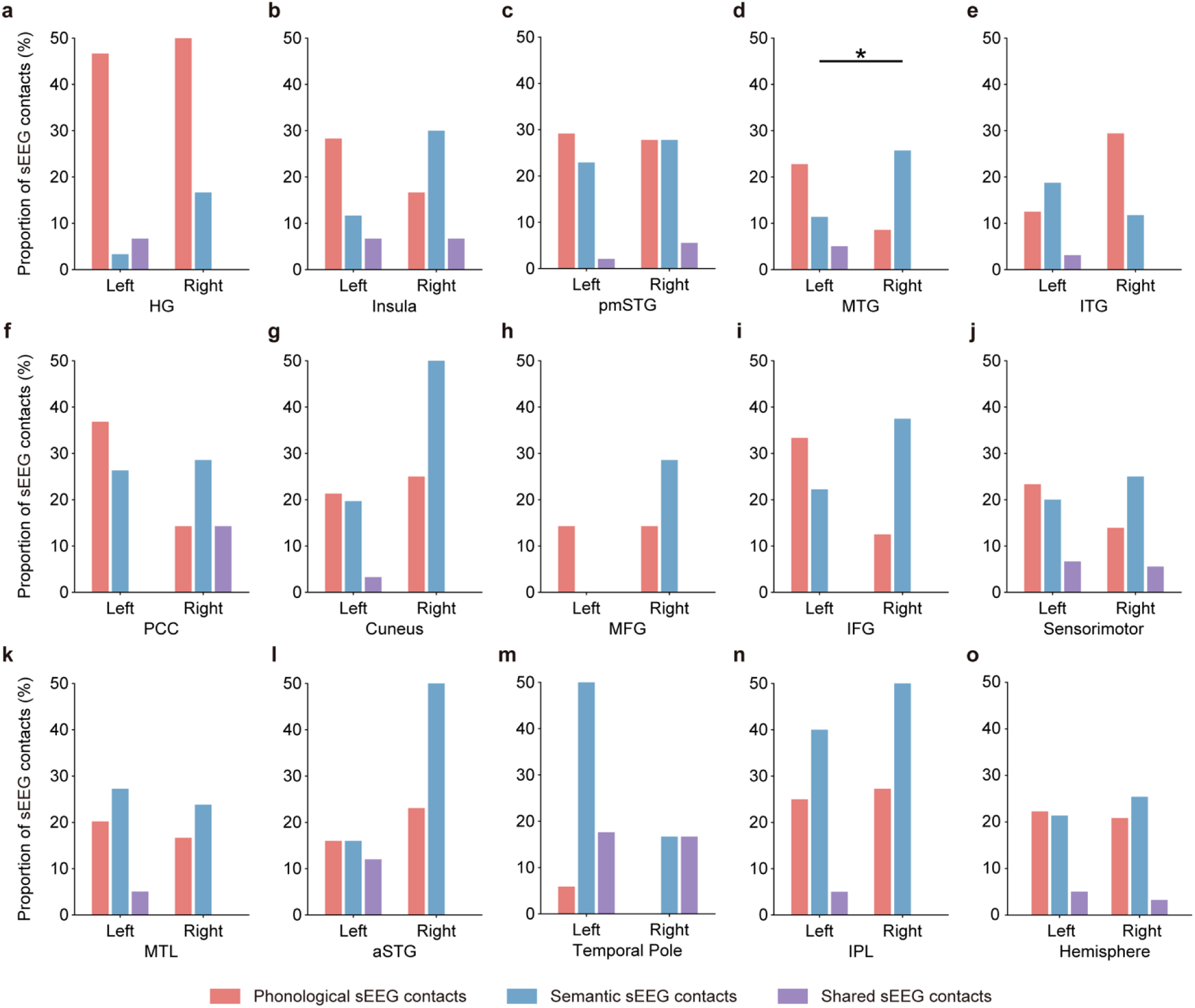
| Hemispheric and regional distribution of phonological, semantic, and shared sEEG contacts. Bar plots show the proportion of sEEG contacts classified as phonological (*red*), semantic (*blue*), and shared/P2S-transfer (*purple*) in each brain region, separated by hemisphere (left, right). Brain regions: *HG: Heschl’s gyrus; pmSTG: posterior and middle superior temporal gyrus; MTG: middle temporal gyrus; ITG: inferior temporal gyrus; PCC: posterior cingulate cortex; Cuneus: cuneus and precuneus; MFG: middle frontal gyrus; IFG: inferior frontal gyrus; Sensorimotor: sensorimotor cortex; MTL: medial temporal lobe; aSTG: anterior superior temporal gyrus; IPL: inferior parietal lobule.* Asterisk denotes significant hemispheric difference (chi-square test, \**P* < 0.05).

**Extended Data Fig. 3.**
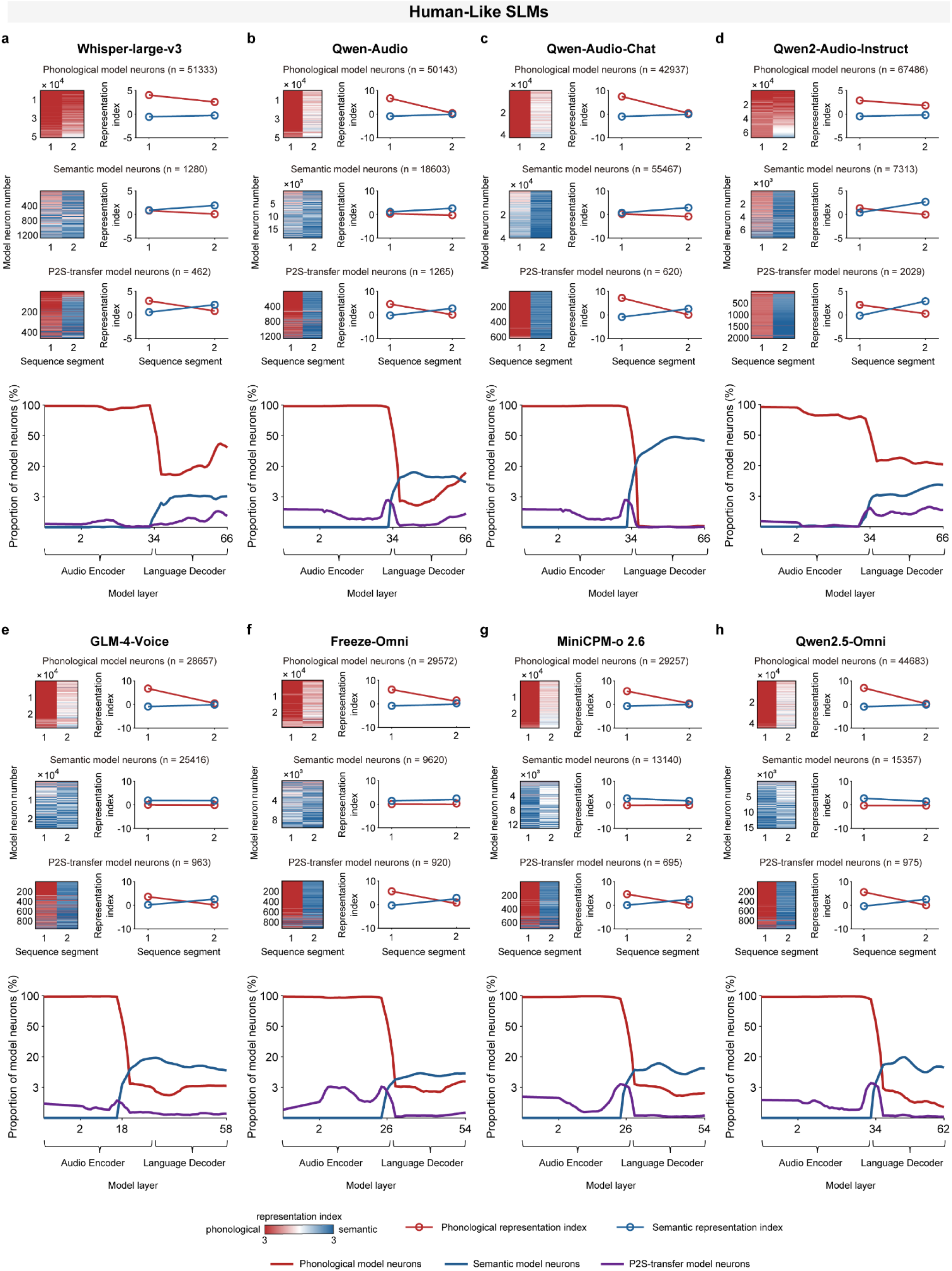
| Sequential and hierarchical organization of phonological, semantic and P2S-transfer model neurons in additional human-like SLMs. **a–h**, Sequential and hierarchical organization in Whisper-large-v3 (a), Qwen-Audio (b), Qwen-Audio-Chat (c), Qwen2-Audio-Instruct (d), GLM-4-Voice (e), Freeze-Omni (f), MiniCPM-o 2.6 (g), and Qwen2.5-Omni (h). For each model, the top three rows show the sequential organization of phonological, semantic, and P2S-transfer model neurons, respectively. Heatmaps show the sequence-resolved dominant representation index for sequence segments 1 and 2 in individual neurons, sorted by dominant representation index, and line plots show mean phonological (*red*) and semantic (*blue*) representation indices across sequence segments. As in the human brain and Qwen2-Audio (Fig. 2), P2S-transfer neurons exhibit a transition from phonological to semantic representation across the sequence. The bottom row shows hierarchical organization for each model, with the proportions of phonological (*red*), semantic (*blue*), and P2S-transfer (*purple*) model neurons plotted across model layers spanning the audio encoder and language decoder (module boundaries indicated). Consistent with a human-like organizational pattern, phonological neurons were enriched in the audio encoder and early decoder layers, whereas semantic neurons became more prominent in later decoder layers.

**Extended Data Fig. 4.**
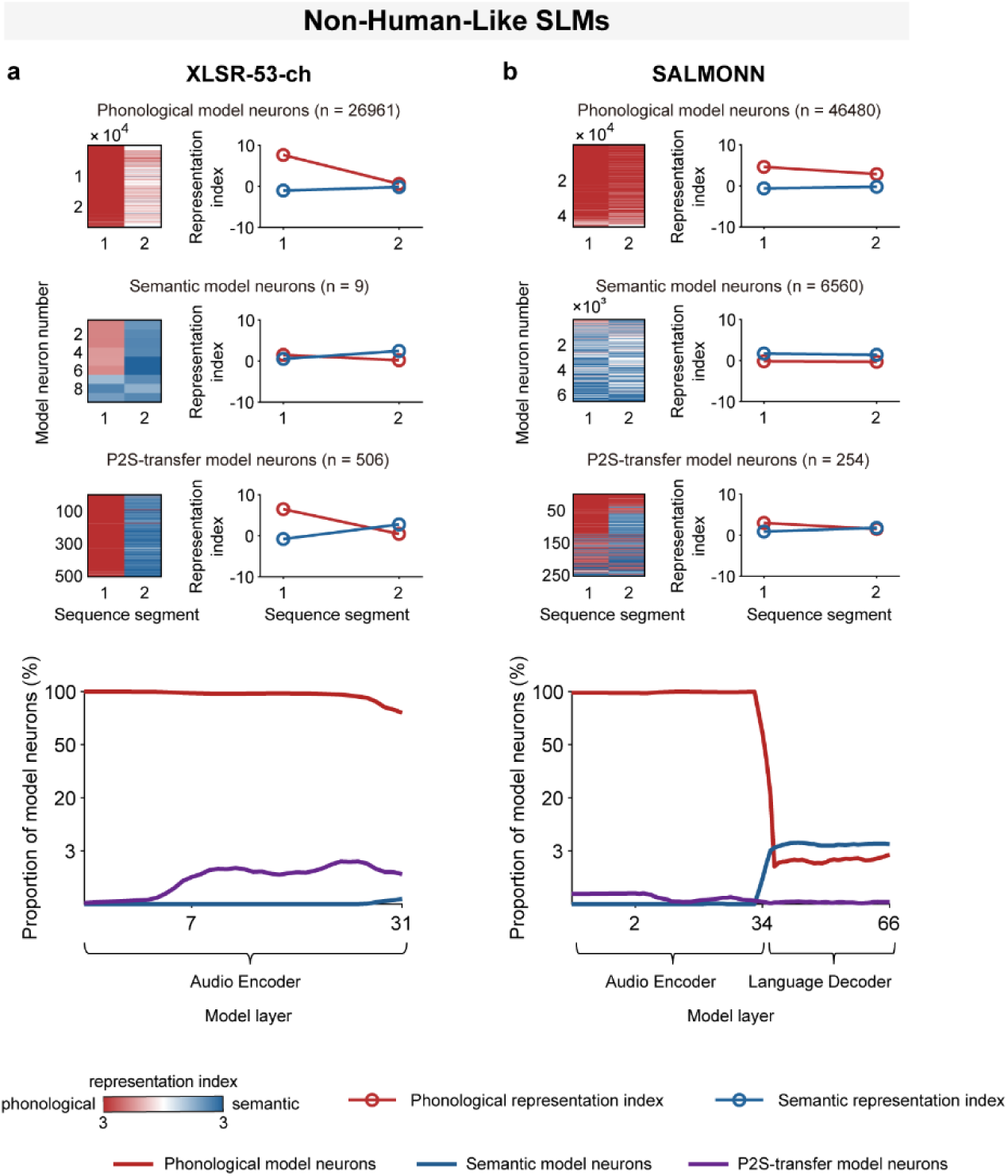
| Sequential and hierarchical organization of phonological, semantic and P2S-transfer model neurons in additional non-human-like SLMs. **a,b**, Sequential and hierarchical organization in XLSR-53-ch (a) and SALMONN (b). For each model, the top three rows show the sequential organization of phonological, semantic, and P2S-transfer model neurons, respectively. Heatmaps show the sequence-resolved dominant representation index for sequence segments 1 and 2 in individual neurons, sorted by dominant representation index, and line plots show mean phonological (*red*) and semantic (*blue*) representation indices across sequence segments. Unlike human-like SLMs (Fig. 2 and Extended Data Fig. 3), these models showed weaker or less distinct phonology-to-semantics transition profiles in P2S-transfer neurons, together with reduced sequential separation between phonological and semantic representations. The bottom row shows hierarchical organization for each model, with the proportions of phonological (*red*), semantic (*blue*), and P2S-transfer (*purple*) neurons plotted across model layers spanning the audio encoder and language decoder, where applicable (module boundaries indicated).

**Extended Data Fig. 5.**
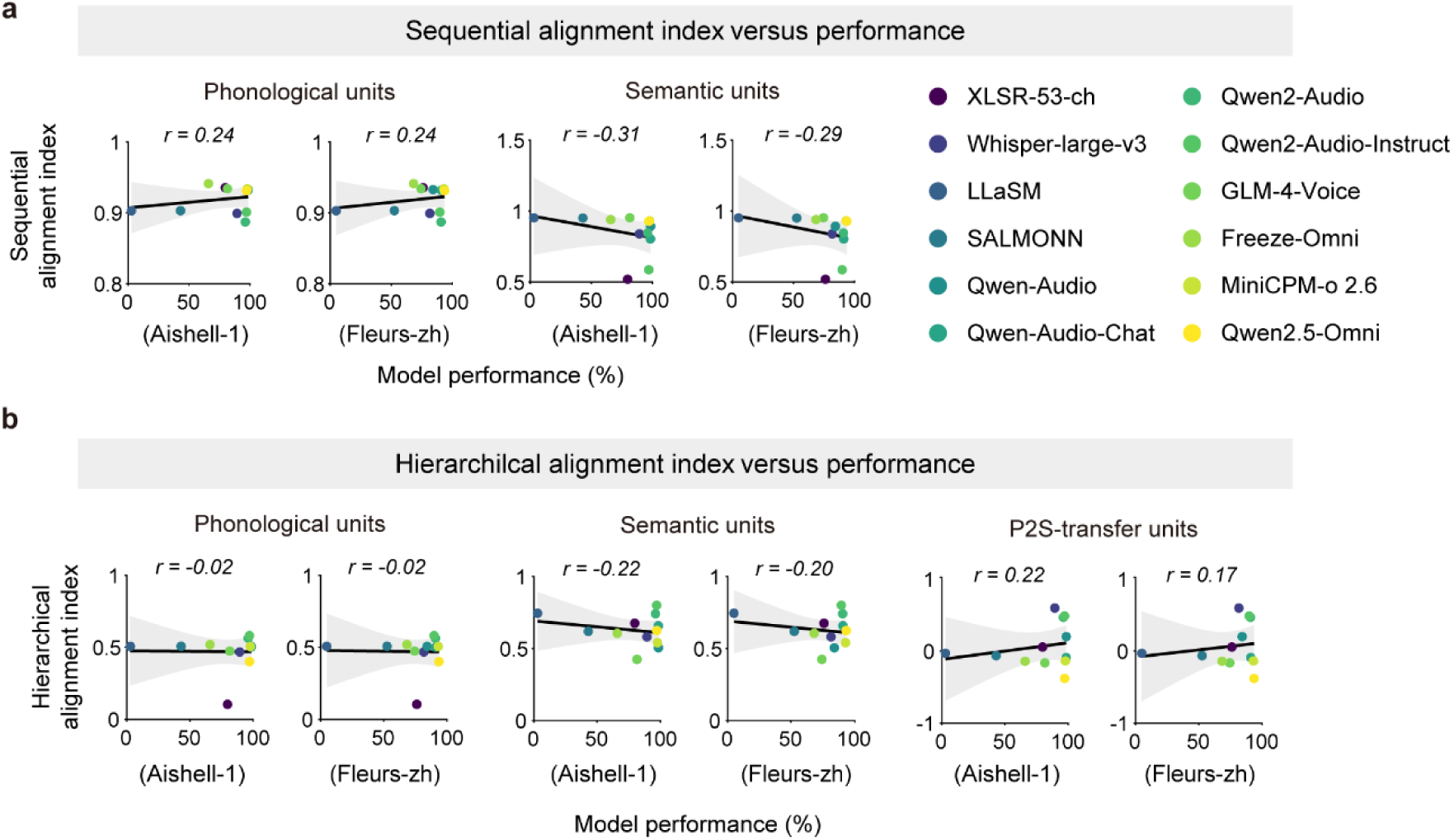
| Relationship between brain–SLM alignment and model performance for phonological, semantic, and P2S-transfer units. **a**, Sequential alignment index as a function of model performance on two Mandarin speech recognition benchmarks (Aishell-1 and Fleurs-zh) for phonological units (*left*) and semantic units (*right*). Each point represents one SLM, color-coded by model identity (see legend). Linear regression lines with shaded 95% confidence intervals are shown; Pearson correlation coefficient (*r*) is indicated for each plot. **b,** Hierarchical alignment index as a function of model performance on Aishell-1 and Fleurs-zh benchmarks for phonological units (*left*), semantic units (*middle*), and P2S-transfer units (*right*). Visualization format as in **a**.

**Extended Data Fig. 6.**
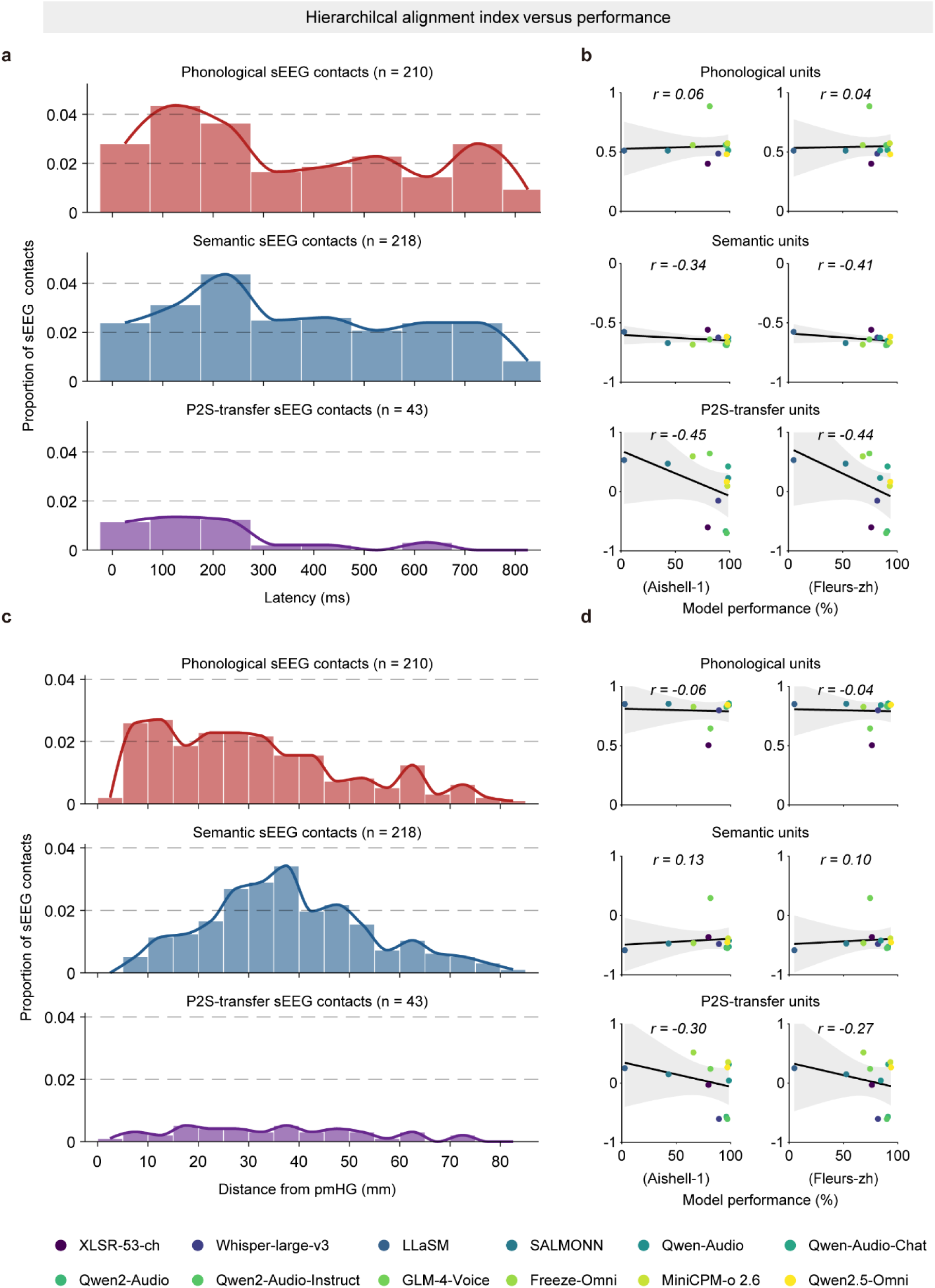
| Brain–SLM hierarchical alignment based on alternative definitions of human hierarchical organization. **a**, Human hierarchical organization quantified as the distribution of phonological (*red*), semantic (*blue*), and P2S-transfer (*purple*) sEEG contacts across representation onset latency. Histograms show the proportion of contacts across latency bins, with overlaid density curves. **b,** Relationship between model performance and the hierarchical alignment index computed using the latency-based human distributions in **a** for phonological units (*top*), semantic units (*middle*), and P2S-transfer units (*bottom*), evaluated on the Aishell-1 (*left*) and Fleurs-zh (*right*) benchmarks. **c,** Human hierarchical organization quantified as the distribution of phonological, semantic, and P2S-transfer sEEG contacts as a function of distance from posteromedial Heschl’s gyrus (pmHG). Histograms show the proportion of contacts across distance bins, with overlaid density curves. **d,** Relationship between model performance and the hierarchical alignment index computed using the distance-from-pmHG human distributions in **c** for phonological units (*top*), semantic units (*middle*), and P2S-transfer units (*bottom*), shown as in **b**. For **b** and **d**, the hierarchical alignment index was defined as the Pearson correlation (r) between the human distributions in **a** or **c** and the distribution of corresponding unit proportions across model layers in SLMs. Each point represents one SLM, color-coded by model identity (see legend). Linear regression lines with shaded 95% confidence intervals are shown; Pearson correlation coefficient (*r*) is indicated for each plot. No significant correlations were observed.

**Extended Data Fig. 7.**
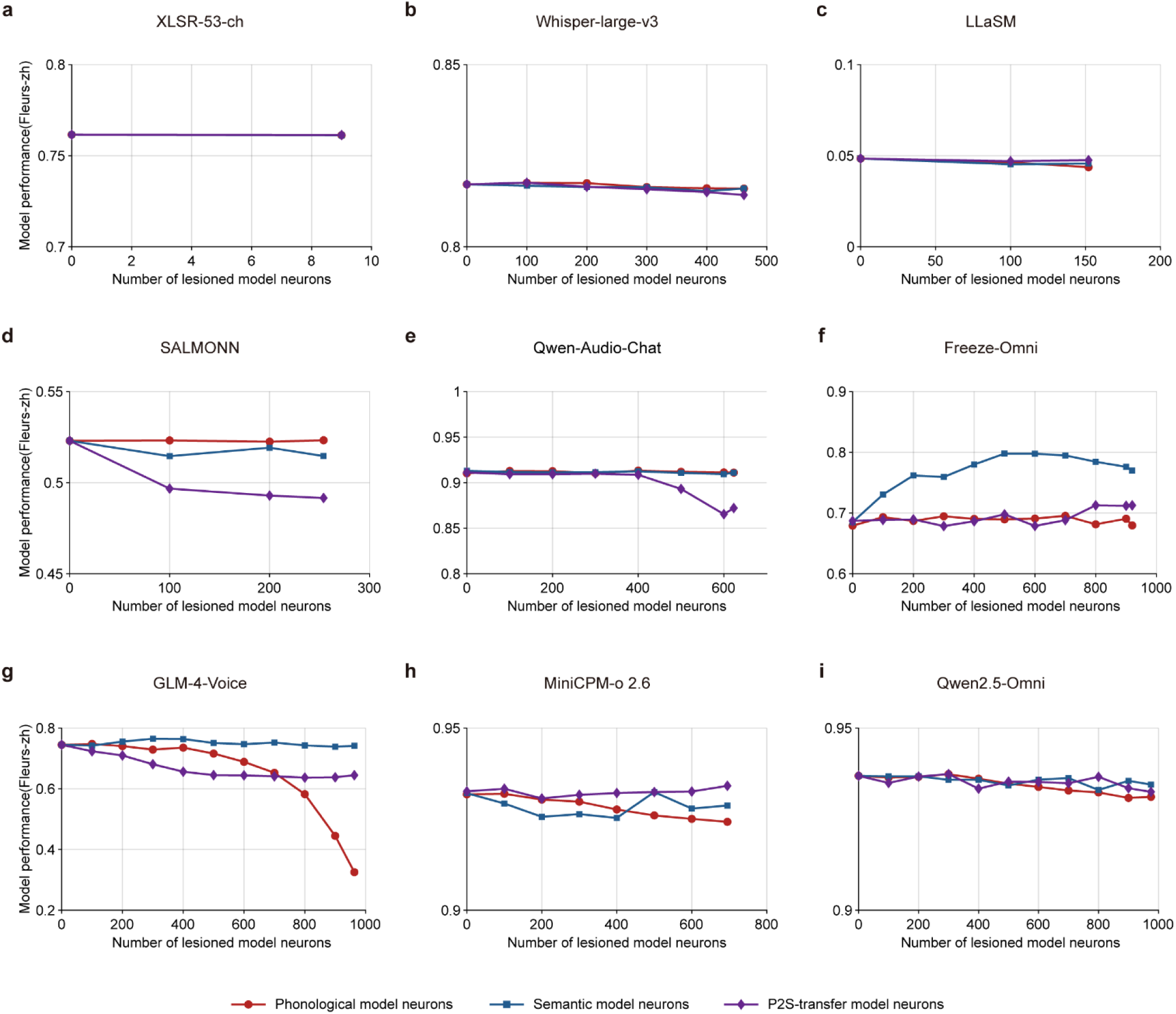
| Effect of targeted lesioning on model performance across nine SLMs. **a–i**, Model performance on the Fleurs-zh benchmark as a function of the number of lesioned neurons for XLSR-53-ch (**a**), Whisper-large-v3 (**b**), LLaSM (**c**), SALMONN (**d**), Qwen-Audio-Chat (**e**), Freeze-Omni (**f**), GLM-4-Voice (**g**), MiniCPM-o 2.6 (**h**), and Qwen2.5-Omni (**i**). Separate curves show lesioning of phonological (*red*), semantic (*blue*), and P2S-transfer (*purple*) neurons. Lesioning paradigm is the same as in Fig. 4.

**Extended Data Fig. 8.**
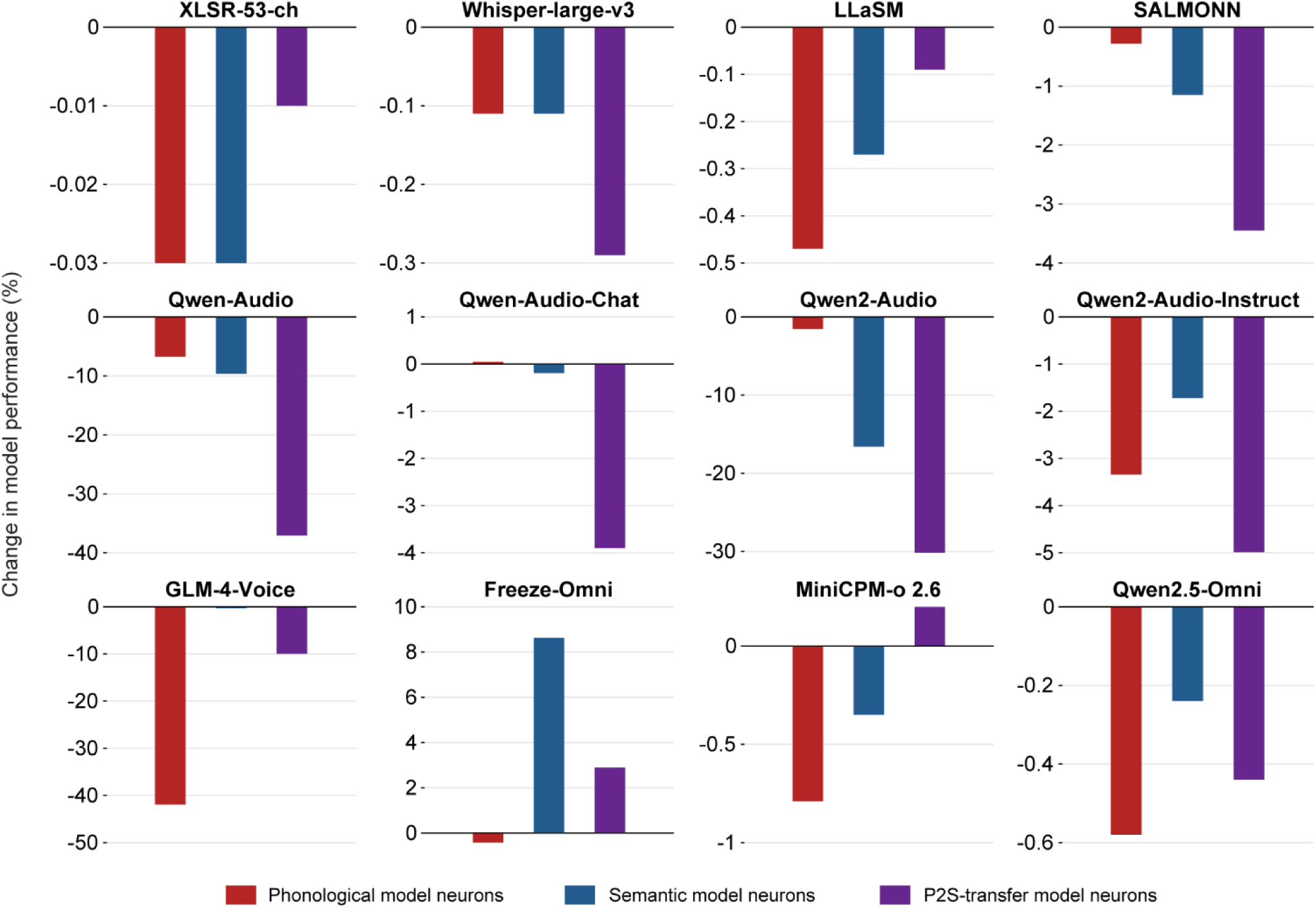
| Change in model performance after matched targeted lesioning across 12 SLMs. For each SLM (XLSR-53-ch, Whisper-large-v3, LLaSM, SALMONN, Qwen-Audio, Qwen-Audio-Chat, Qwen2-Audio, Qwen2-Audio-Instruct, GLM-4-Voice, Freeze-Omni, MiniCPM-o 2.6, and Qwen2.5-Omni), bars show the change in model performance (%) on the Fleurs-zh benchmark after lesioning phonological (*red*), semantic (*blue*), or P2S-transfer (*purple*) model neurons. To enable matched comparison across unit classes within each model, the number of lesioned neurons was fixed to the maximum common lesion size, defined as the smallest unit count among the three classes in that model. Negative values indicate performance impairment relative to the intact model, whereas positive values indicate improved performance. The lesioning paradigm is the same as in Fig. 4 and Extended Data Fig. 7.

**Extended Data Fig. 9.**
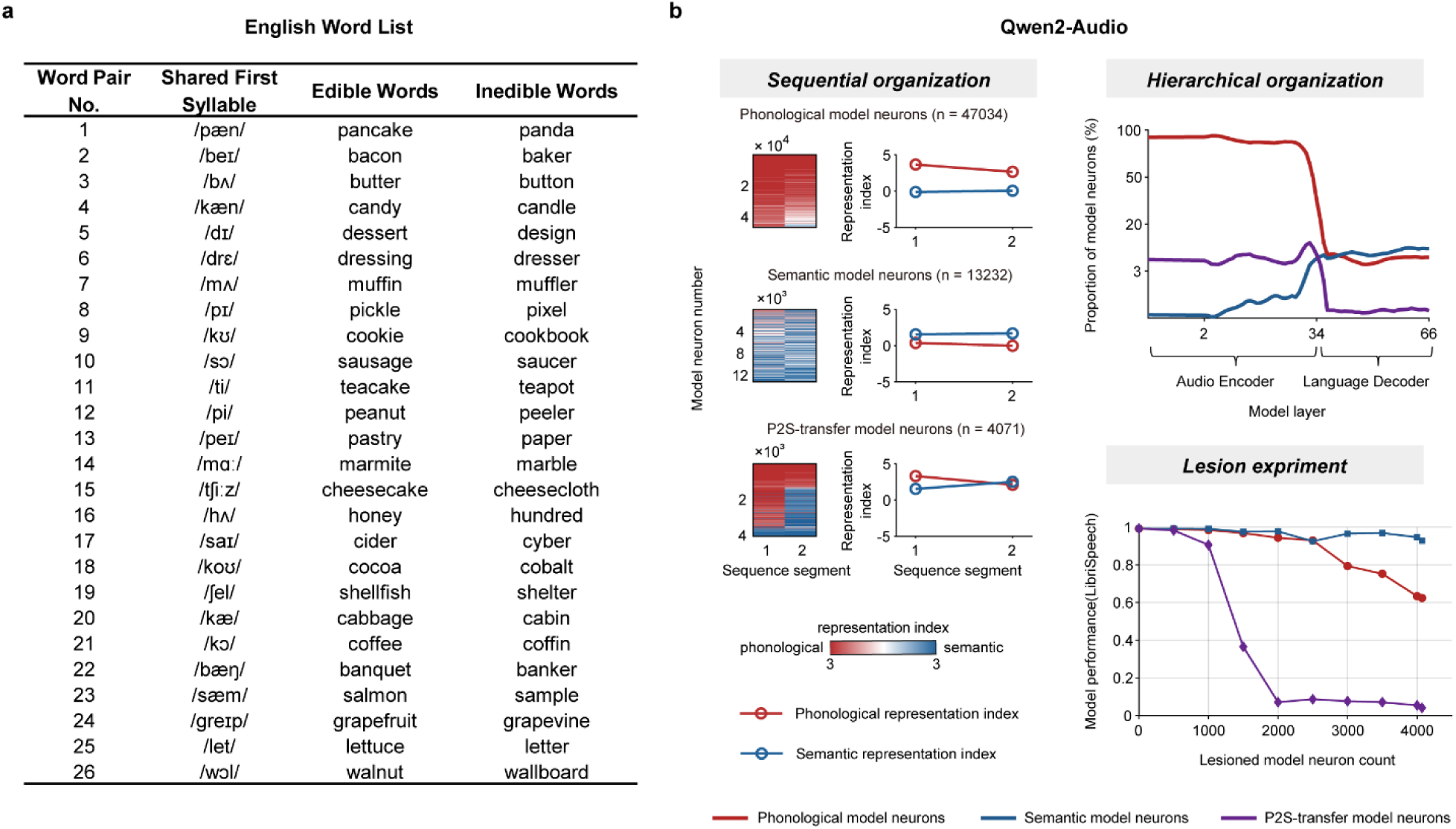
| Sequential organization, hierarchical organization and lesion effects in Qwen2-Audio on an English word dataset. **a**, English word list used for analysis. The dataset comprised 26 word pairs, each sharing the same first syllable and consisting of one edible word and one inedible word. **b,** Sequential organization, hierarchical organization and lesioning experiment results for Qwen2-Audio on the English dataset. In the sequential organization analysis (*left*), phonological, semantic, and P2S-transfer model neurons are shown from top to bottom. Heatmaps show the sequence-resolved dominant representation index for sequence segments 1 and 2 in individual neurons, sorted by dominant representation index, and line plots show mean phonological (*red*) and semantic (*blue*) representation indices across sequence segments. As in the Mandarin dataset, P2S-transfer neurons exhibited a transition from phonological to semantic representation across the sequence. In the hierarchical organization analysis (*top right*), the proportions of phonological (*red*), semantic (*blue*), and P2S-transfer (*purple*) model neurons are plotted across model layers spanning the audio encoder and language decoder. In the lesion experiment (*bottom right*), model performance on the LibriSpeech benchmark is plotted as a function of the number of lesioned neurons for phonological (*red*), semantic (*blue*), and P2S-transfer (*purple*) neuron classes. Targeted lesioning of P2S-transfer neurons caused the strongest performance impairment, consistent with their critical role in phonology-to-semantics transformation.

**Extended Data Fig. 10.**
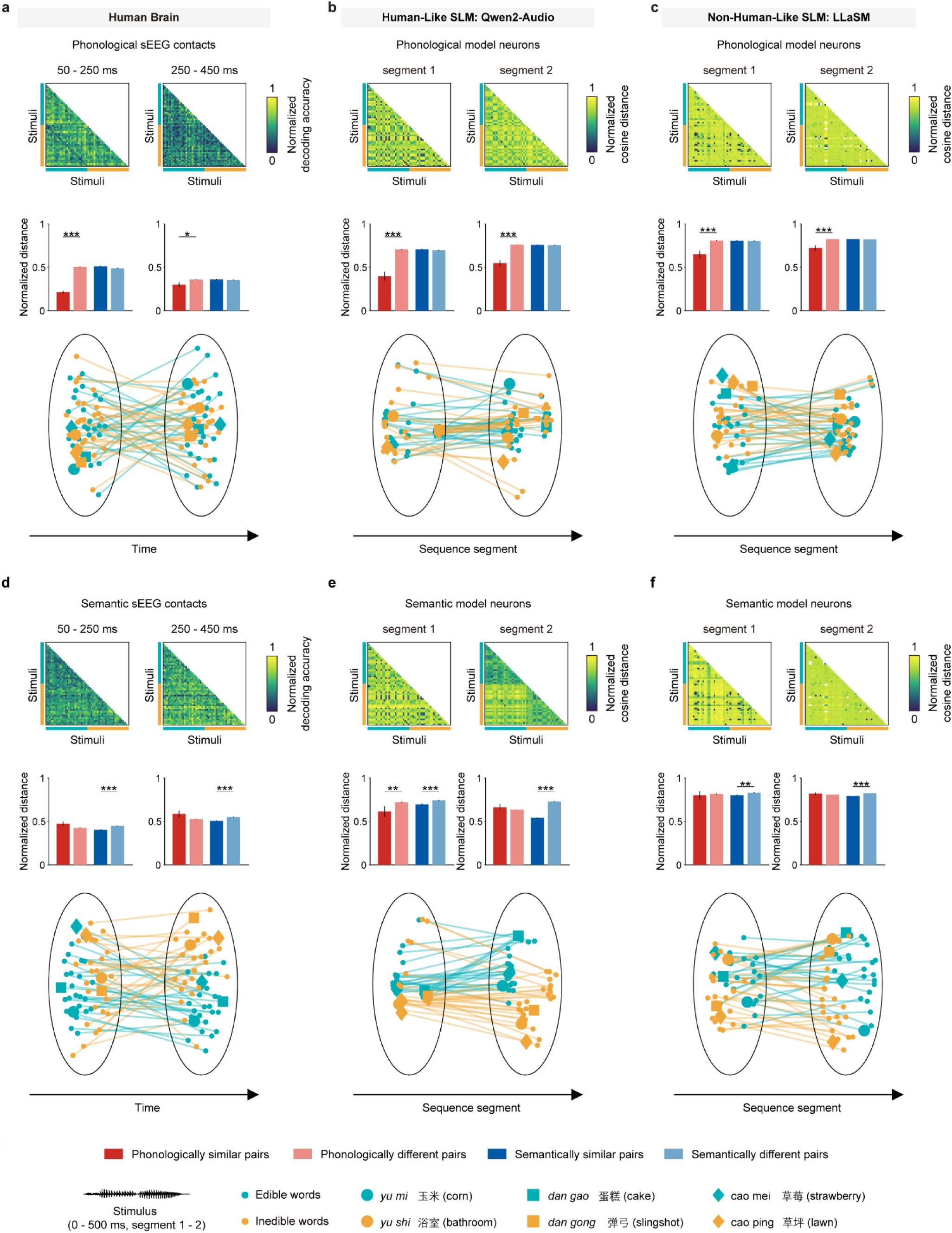
| Phonological and semantic units maintain dimension-specific representational geometry in both the human brain and SLMs without pronounced phonology-to-semantics reconfiguration. **a–c**, Representational geometry of phonological units in the human brain (**a**), the human-like SLM Qwen2-Audio (**b**), and the non-human-like SLM LLaSM (**c**). **Top,** RDMs are shown for early (50–250 ms) and late (250–450 ms) temporal sequence windows in the human brain, and for sequence segments 1 and 2 in SLMs. In the human brain, RDMs are based on pairwise decoding accuracy; in SLMs, RDMs are based on cosine distance. **Middle,** quantification of representational distances between word pairs grouped by phonological and semantic similarity. Bars show normalized distances for phonologically similar versus phonologically different pairs and semantically similar versus semantically different pairs (colors as indicated). **Bottom,** two-dimensional embeddings via multidimensional scaling (MDS) of the representational transition across temporal windows or sequence segments. Each point represents one word, colored by semantic category. Three phonologically similar word pairs are highlighted with matching distinct shapes, and lines connect the same word across temporal sequence windows (**a**) or sequence segments (**b, c**). Phonological units preserve predominantly phonological geometry across the sequence, with limited semantic reconfiguration. **d–f,** Representational geometry of semantic units in the human brain (**d**), Qwen2-Audio (**e**), and LLaSM (**f**), shown as in **a–c**. Semantic units maintain predominantly semantic geometry across temporal windows or sequence segments, with minimal phonological-to-semantic transformation. Across both biological and artificial systems, these results indicate that stable dimension-specific geometry in phonological and semantic units contrasts with the dynamic geometric reconfiguration observed in P2S-transfer units. Asterisks denote significance (\**P* < 0.05; \*\**P* < 0.01; \*\*\**P* < 0.001; one-sided two-sample t-test).

