## Supplementary figures and tables for "Human-like sequential sound-to-meaning transfer drives artificial speech comprehension"

[#] Corresponding authors:

#### Supplementary figures

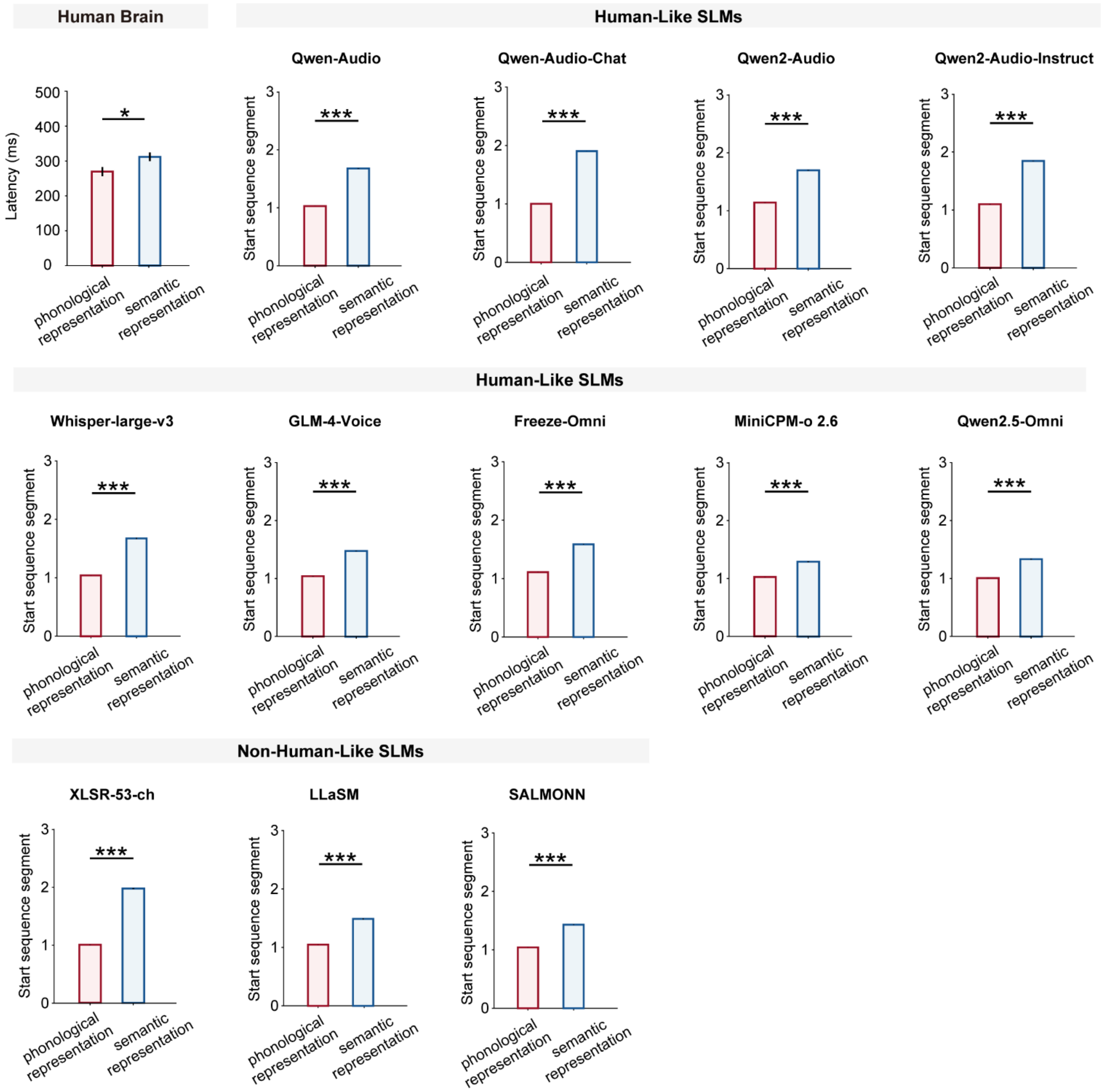


**Supplementary Fig. 1 | Sequential onset of phonological and semantic representations in the human brain and SLMs.** Latency of phonological (*red*) and semantic (*blue*) representations in the human brain, and the onset sequence segment of phonological and semantic representations in human-like SLMs (Qwen-Audio, Qwen-Audio-Chat, Qwen2-Audio, Qwen2-Audio-Instruct, Whisper-large-v3, GLM-4-Voice, Freeze-Omni, MiniCPM-o 2.6, and Qwen2.5-Omni) and non-human-like SLMs (XLSR-53-ch, LLaSM, and SALMONN). Error bars indicate SEM. Asterisks indicate significant differences (two-sided two-sample *t*-test; *$P < 0.05$, ***$P < 0.001$).

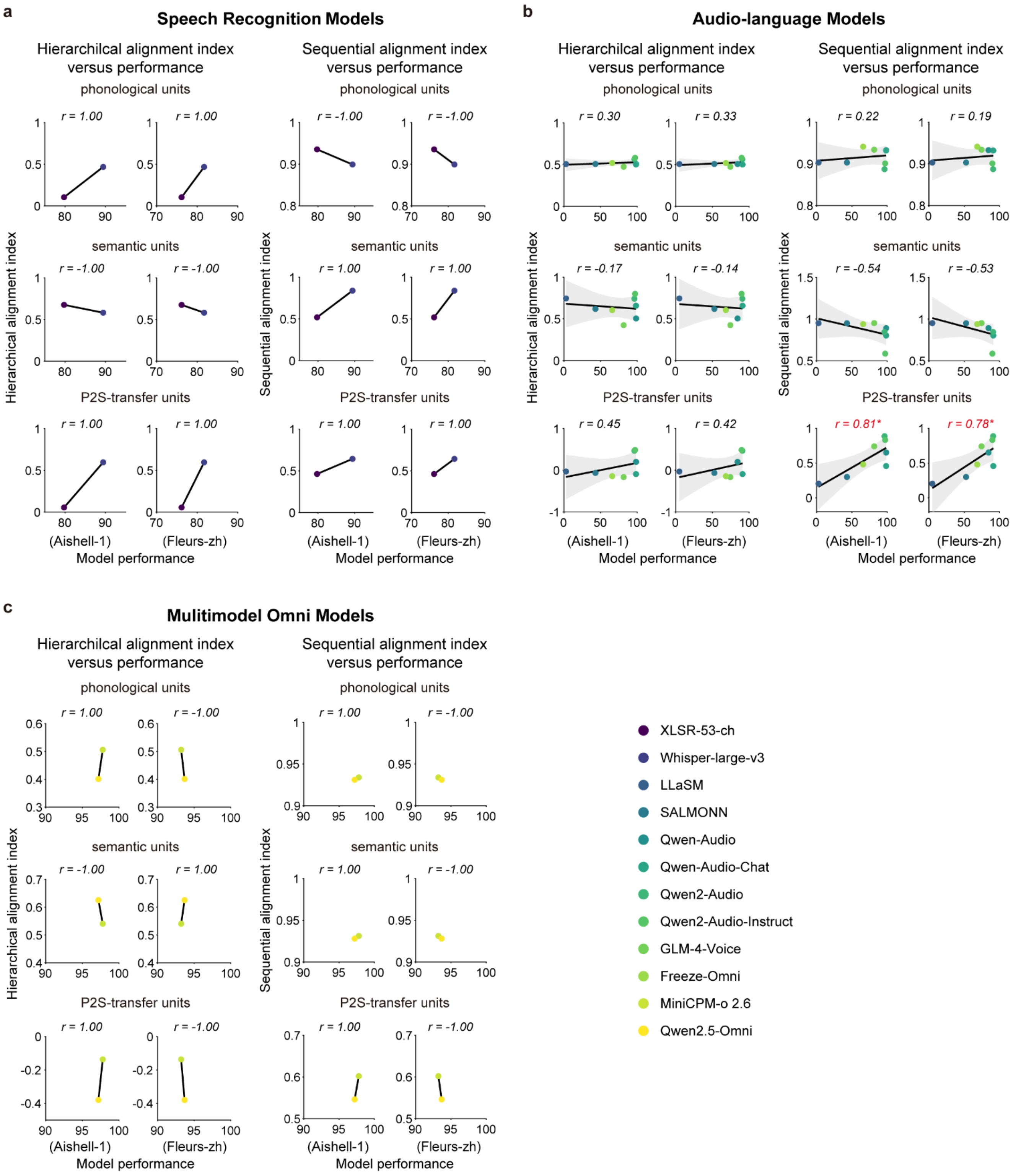


**Supplementary Fig. 2** | Hierarchical and sequential alignment versus model performance across three categories of SLMs. Scatter plots show relationships between alignment indices and model performance for individual SLMs (each dot represents one model, color-coded by model identity; see legend). Linear regression lines with shaded 95% confidence intervals are shown; $r$ values denote Pearson correlation coefficients. Significant correlations are highlighted in color (*$P < 0.05$, *red*). Family-specific analyses for model families with $N = 2$ are presented for descriptive purposes only. **a,** Speech recognition models (Whisper-large-v3, XLSR-53-ch). Hierarchical (*left*) and sequential (*right*) alignment indices for phonological, semantic, and P2S-transfer units are plotted against model performance on the Aishell-1 (*left*) and Fleurs-zh (*right*) datasets. **b,** Audio–language models (Qwen-Audio, Qwen-Audio-Chat, Qwen2-Audio, Qwen2-Audio-Instruct, LLaSM, GLM-4-Voice, Freeze-Omni). Analyses follow the same format as in **a,** showing relationships between alignment indices and model performance. **c,** Multimodal omni models (Qwen2.5-Omni, MiniCPM-o 2.6). Corresponding hierarchical and sequential alignment indices are plotted against model performance.

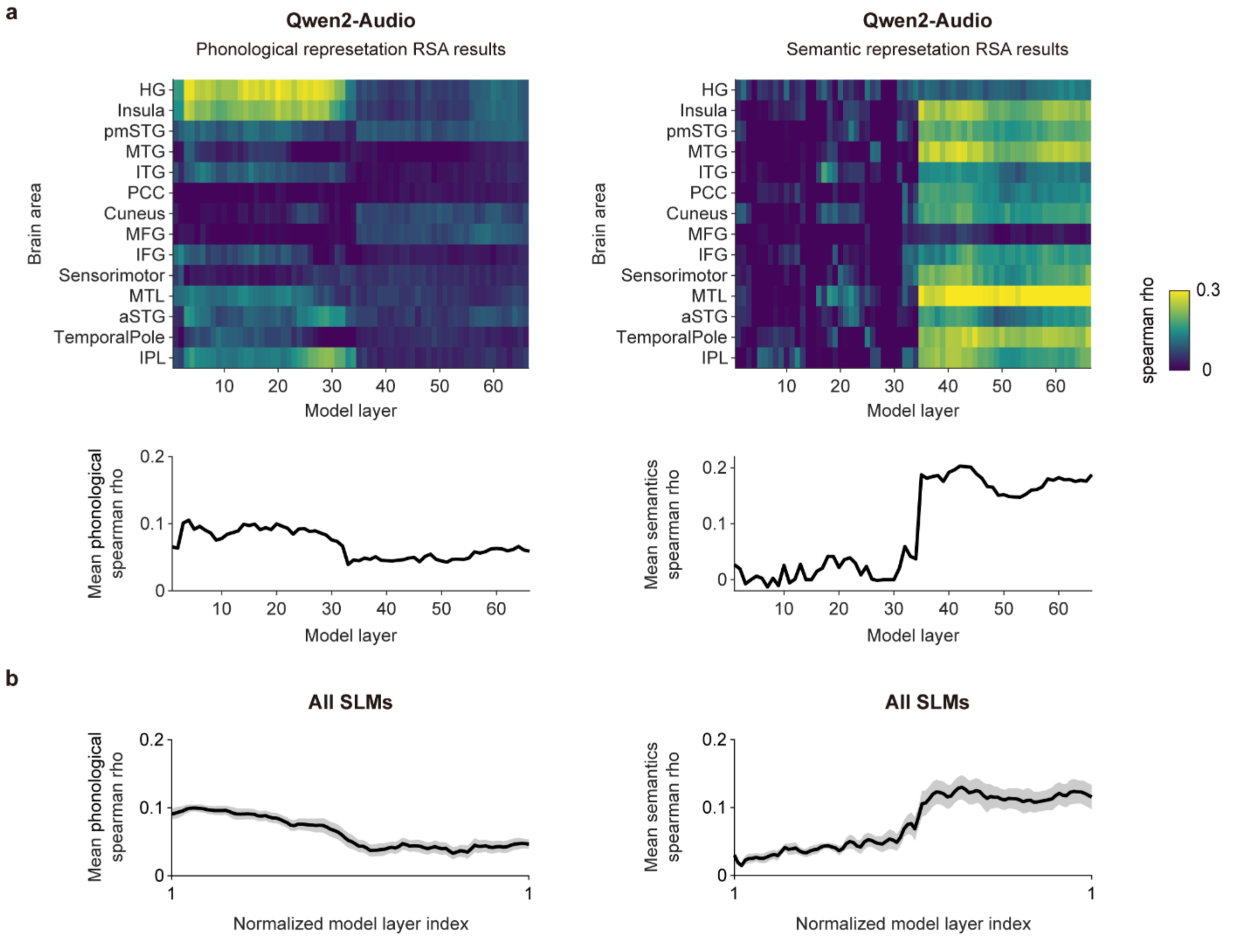


**Supplementary Fig. 3 | Brain–model representational similarity analysis (RSA) of phonological and semantic representations across brain areas and model layers. a,** Phonological (*left*) and semantic (*right*) RSA results for Qwen2-Audio. Heatmaps (*top*) show the Spearman correlation coefficients (*r*) between the average phonological or semantic RDMs of brain areas and those of model layers. The line plots (*bottom*) show the mean Spearman correlation for phonological and semantic representations, illustrating that shallow layers exhibit greater phonological representation alignment with the human brain, while deeper layers show greater semantic representation alignment. **b,** Grand-average phonological (*left*) and semantic (*right*) RSA results across all SLMs after normalizing model layer indices to the range [0, 1]. Solid lines denote the mean Spearman correlation across models, and shaded regions indicate SEM across models. Consistent with panel **a,** the across-model average shows that shallower layers tend to align more strongly with phonological representations in the human brain, while deeper layers demonstrate greater alignment with semantic representations.

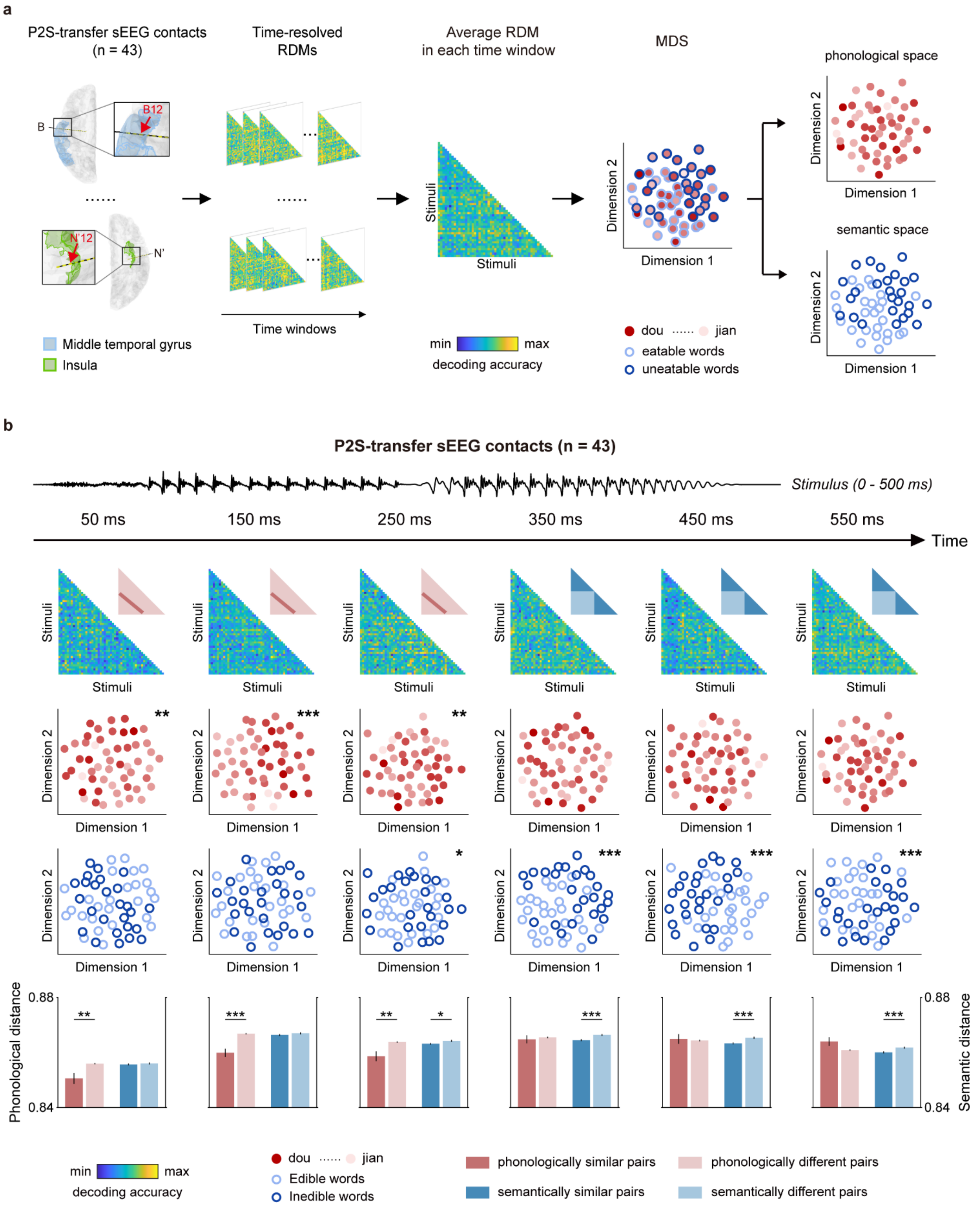


**Supplementary Fig. 4 | Representational geometry of P2S-transfer sEEG contacts in the human brain. a,** Schematic of the representational geometry analysis in P2S-transfer sEEG contacts. Sequence-resolved RDMs (*left*) were constructed from pairwise decoding accuracies among 52 Mandarin words and averaged within each temporal sequence window, projected into two-dimensional spaces via multidimensional scaling (MDS; *middle*). Each circle indicates one word. In the phonological space (*top right*), the circle faces span a continuous red gradient from light to dark, encoding 26 word pairs that share the same first syllable. In the semantic space (*bottom right*), the circle edges are light or dark blue to denote edible or inedible semantic categories. **b**, Visualizing phonological-to-semantic representations in P2S-transfer sEEG contacts (n = 43). At each time window centered on 50, 150, 250, 350, 450 and 550 ms (±100 ms), RDMs (*top rows*), MDS projections in phonological and semantic spaces (*middle rows*), and bar plots of distances between similar and different word pairs (*bottom rows*) are shown. P2S-transfer sEEG contacts exhibit a progressive shift in representational geometry from phonological to semantic across successive time windows. Statistical significance was assessed by one-sided two-sample *t*-tests (*$P < 0.05$, **$P < 0.01$, ***$P < 0.001$). Error bars indicate SEM.

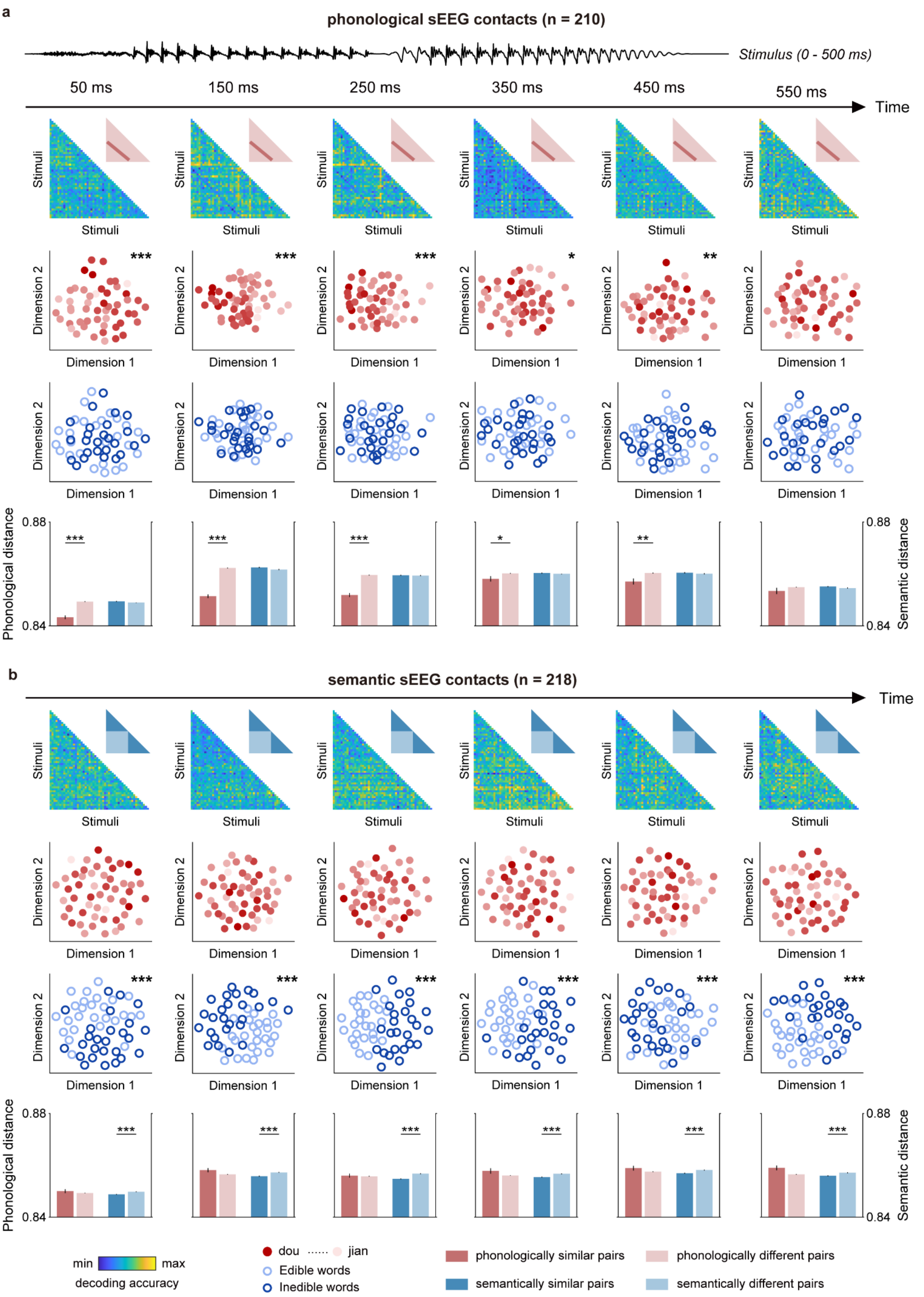


**Supplementary Fig. 5 | Representational geometry of phonological and semantic sEEG contacts in the**

**human brain. a,** Visualizing representations in phonological sEEG contacts ($n = 210$) across time. At each temporal sequence window centered on 50, 150, 250, 350, 450 and 550 ms (±100 ms), RDMs (*top rows*), MDS projections in phonological and semantic spaces (*middle rows*), and bar plots of distances between similar and different word pairs (*bottom rows*) are shown. Statistical significance was assessed by one-sided two-sample $t$-tests (*$P < 0.05$, **$P < 0.01$, ***$P < 0.001$). Error bars indicate SEM. **b,** Visualizing representations in semantic sEEG contacts ($n = 218$) across time. At each time window centered on 50, 150, 250, 350, 450 and 550 ms (±100 ms), RDMs (*top rows*), MDS projections in phonological and semantic spaces (*middle rows*), and bar plots of distances between similar and different word pairs (*bottom rows*) are shown. Statistical significance was assessed by one-sided two-sample $t$-tests (***$P < 0.001$). Error bars indicate SEM.

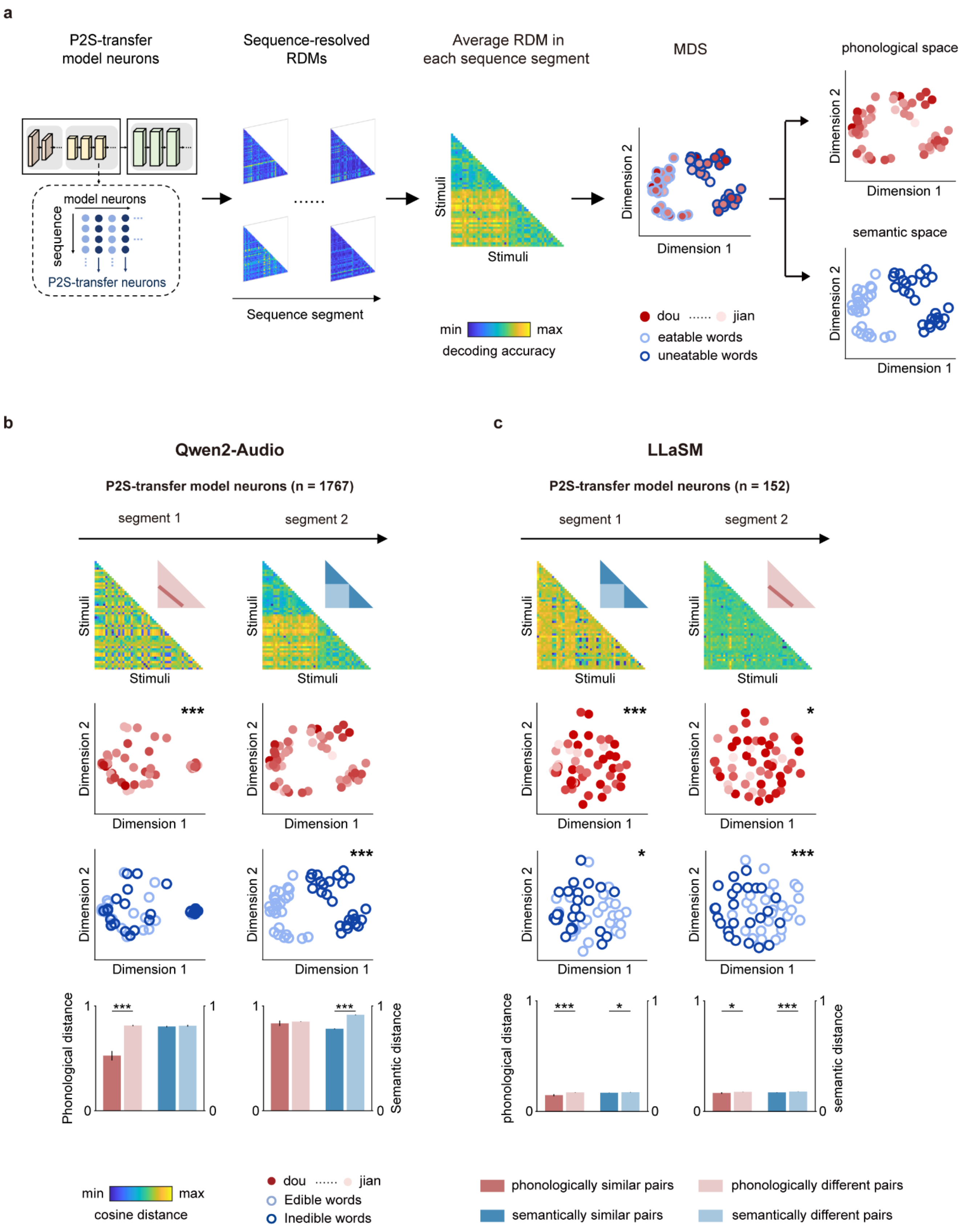


**Supplementary Fig. 6 | Representational geometry of P2S-transfer model neurons in the SLMs. a,** Schematic of the representational geometry analysis in P2S-transfer model neurons. Sequence-resolved RDMs (*left*) were computed from pairwise cosine distances among 52 Mandarin words and averaged within each sequence segment, projected into two-dimensional spaces via MDS (*middle*). Each circle indicates one word. In the phonological space (*top right*), the circle faces span a continuous red gradient from light to dark, encoding 26 word pairs that share the same first syllable. In the semantic space (*bottom right*), the circle edges are light or dark blue to denote edible or inedible semantic categories. **b**, Visualizing phonological-to-semantic representations in P2S-transfer model neurons of Qwen2-Audio ($n$ = 1,767). For segments 1 and 2, RDMs (*top*), MDS projections in phonological and semantic spaces (*middle*) and bar plots of distances between similar and different word pairs (*bottom*) are shown. P2S-transfer model neurons of Qwen2-Audio exhibit a progressive shift in representational geometry from phonological to semantic along the sequence segments. Statistical significance was assessed by one-sided two-sample $t$-tests (***$P$ < 0.001). Error bars indicate SEM. **c**, Visualizing phonological and semantic representations in P2S-transfer model neurons of LLaSM ($n$ = 152), plotted as in **b**; no systematic shift in representational geometry from phonological to semantic along the sequence segments is observed. Statistical significance was assessed by one-sided two-sample $t$-tests (*$P$ < 0.05, ***$P$ < 0.001). Error bars indicate SEM.

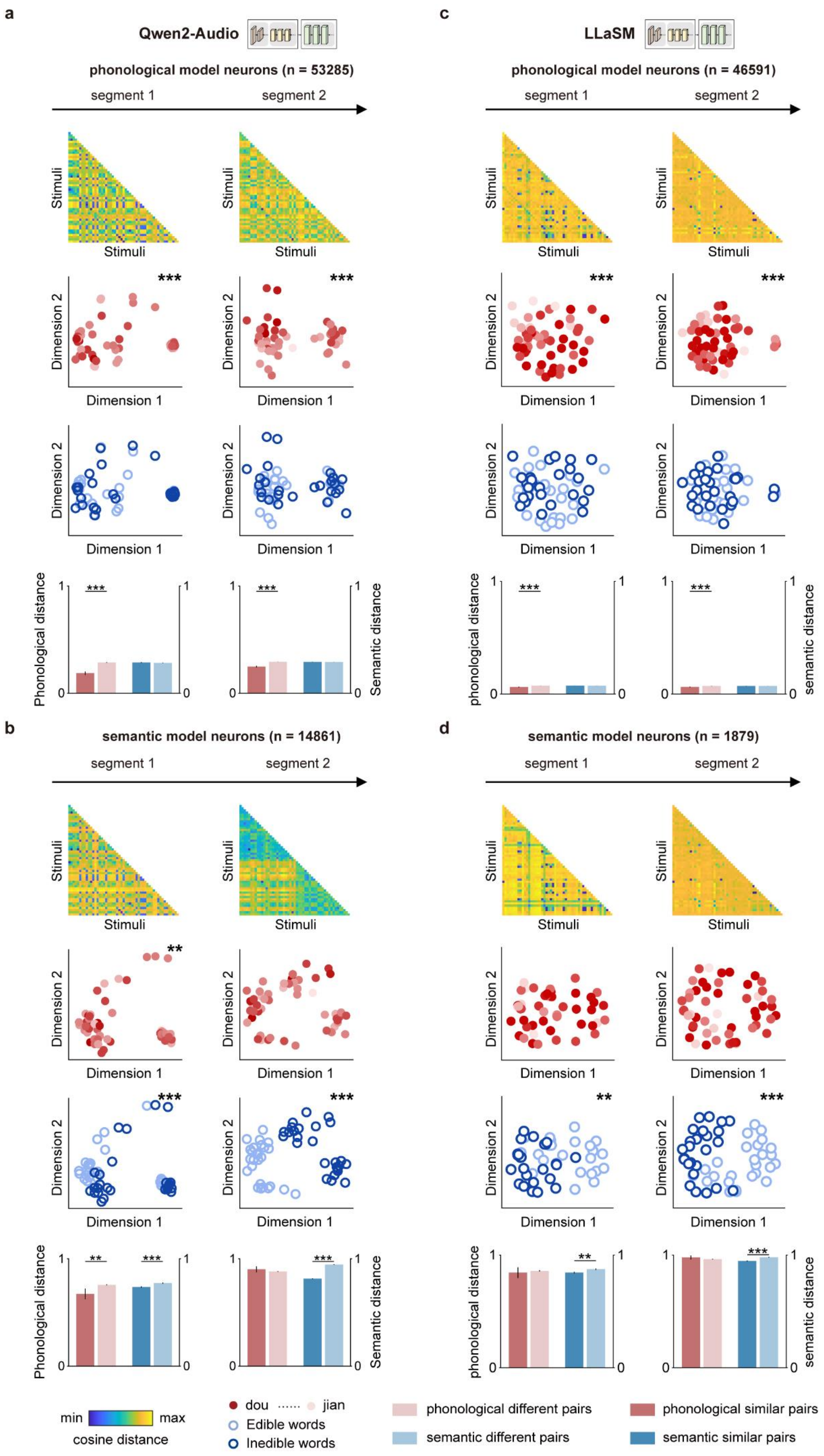

a
Qwen2-Audio
phonological model neurons (n = 53285)
segment 1
segment 2
Stimuli
Dimension 1
Dimension 2
Phonological distance
Semantic distance
***
b
semantic model neurons (n = 14861)
**
c
LLaSM
phonological model neurons (n = 46591)
phonological distance
semantic distance
d
semantic model neurons (n = 1879)
min
max
cosine distance
dou
jian
Edible words
Inedible words
phonological different pairs
phonological similar pairs
semantic different pairs
semantic similar pairs

**Supplementary Fig. 7 | Representational geometry of phonological and semantic model neurons in the SLMs.** **a,** Visualizing representations of phonological model neurons in Qwen2-Audio ($n$ = 53,285) across segment 1 and segment 2. For each sequence segment, RDMs (*top*), MDS projections in phonological and semantic spaces (*middle*), and bar plots of distances between phonologically similar and different pairs, and semantically similar and different pairs (*bottom*) are shown. Statistical significance was assessed by one-sided two-sample $t$-tests (**$P$ < 0.01, ***$P$ < 0.001). Error bars indicate SEM. **b,** Visualizing representations of semantic model neurons in Qwen2-Audio ($n$ = 14,861) across segment 1 and segment *2*. **c,d,** Equivalent analysis for phonological (**c,** $n$ = 46,591) and semantic (**d,** $n$ = 1,879) model neurons in LLaSM. Statistical significance was assessed by one-sided two-sample $t$-tests (*$P$ < 0.05, ***$P$ < 0.001). Error bars indicate SEM.

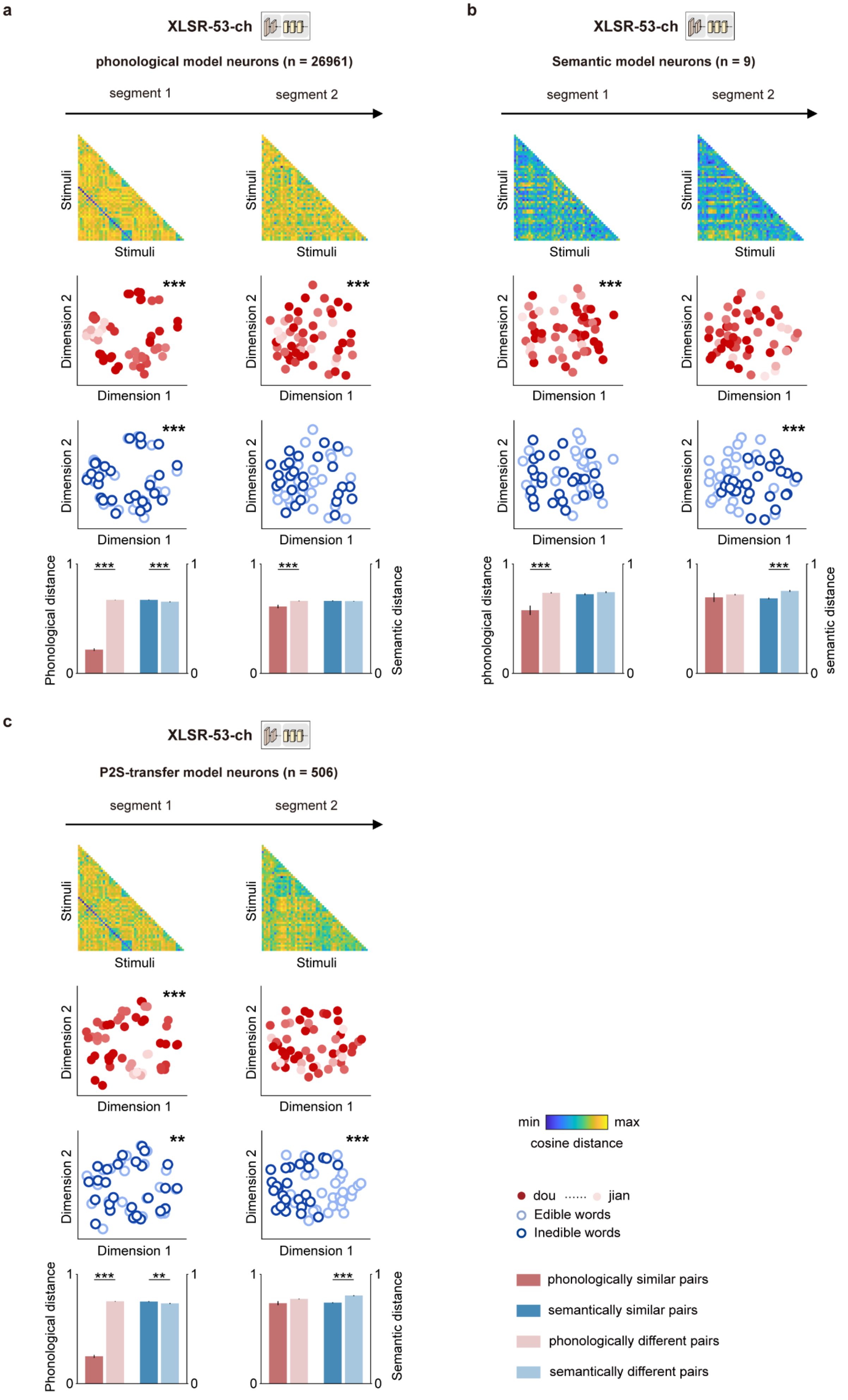
a
XLSR-53-ch
phonological model neurons (n = 26961)
segment 1
segment 2
Stimuli
Dimension 1
Dimension 2
***
Phonological distance
Semantic distance
b
XLSR-53-ch
Semantic model neurons (n = 9)
segment 1
segment 2
phonological distance
semantic distance
c
XLSR-53-ch
P2S-transfer model neurons (n = 506)
segment 1
segment 2
**
min
max
cosine distance
dou
jian
Edible words
Inedible words
phonologically similar pairs
semantically similar pairs
phonologically different pairs
semantically different pairs

**Supplementary Fig. 8 | Representational geometry of phonological, semantic and P2S-transfer model neurons in XLSR-53-ch.** a, Visualizing representations of phonological model neurons in XLSR-53-ch ($n$ = 26,961) across segment 1 and segment 2. For each sequence segment, RDMs (*top*), MDS projections in phonological and semantic spaces (*middle*), and bar plots of distances between phonologically similar and different pairs, and semantically similar and different pairs (*bottom*) are shown. Statistical significance was assessed by one-sided two-sample *t*-tests (**$P < 0.01$, ***$P < 0.001$). Error bars indicate SEM. b**,** Visualizing representations of semantic model neurons in XLSR-53-ch ($n$ = 9) across segment 1 and segment 2, plotted as in a. c**,** Visualizing representations of P2S-transfer model neurons in XLSR-53-ch ($n$ = 506) across segment 1 and segment 2, plotted as in a.

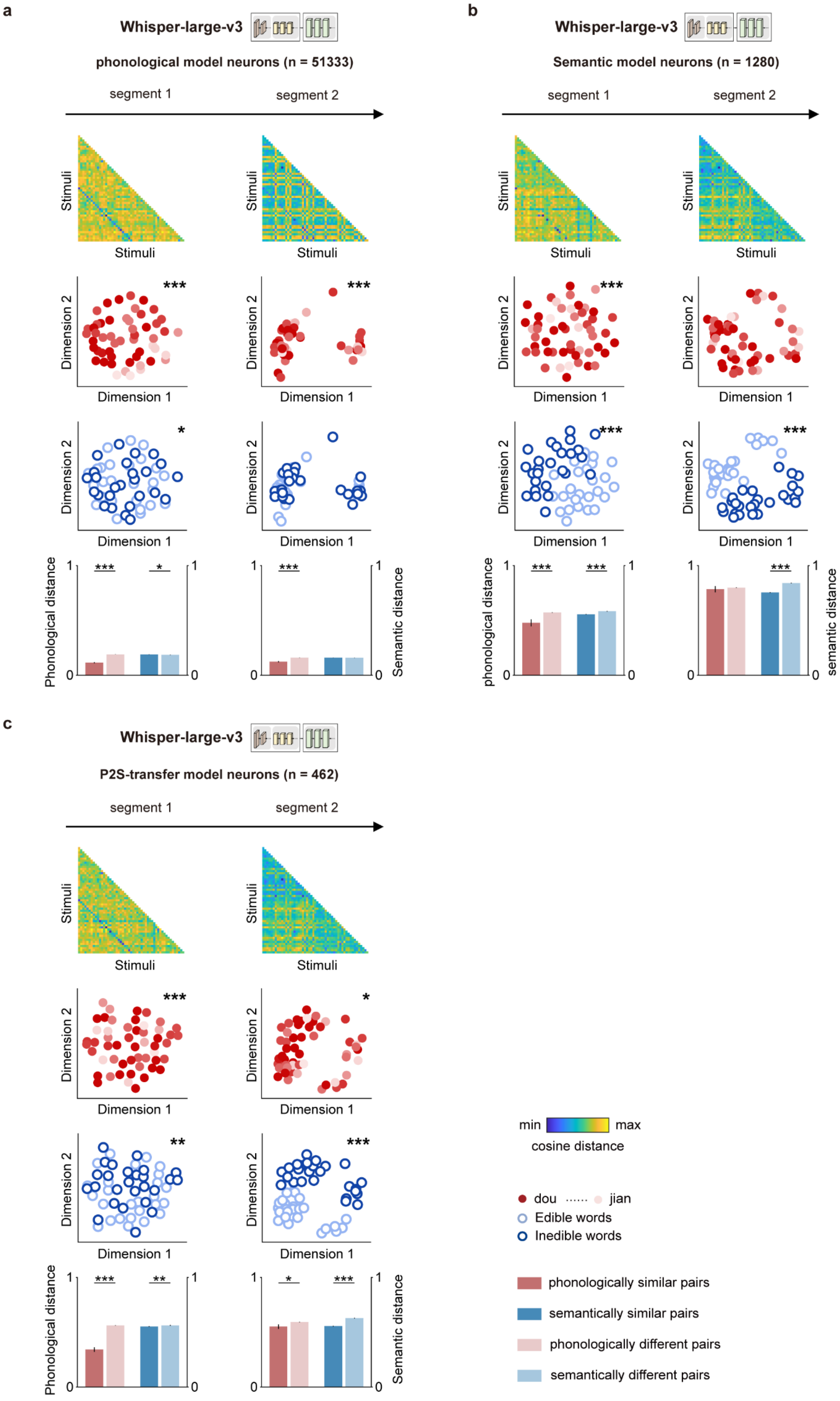

a
Whisper-large-v3
phonological model neurons (n = 51333)
segment 1
segment 2
Stimuli
Dimension 1
Dimension 2
Phonological distance
Semantic distance
b
Whisper-large-v3
Semantic model neurons (n = 1280)
segment 1
segment 2
phonological distance
semantic distance
c
Whisper-large-v3
P2S-transfer model neurons (n = 462)
segment 1
segment 2
min
max
cosine distance
dou
jian
Edible words
Inedible words
phonologically similar pairs
semantically similar pairs
phonologically different pairs
semantically different pairs

**Supplementary Fig. 9 | Representational geometry of phonological, semantic and P2S-transfer model neurons in Whisper-large-v3. a,** Visualizing representations of phonological model neurons in Whisper-large-v3 ($n$ = 51,333) across segment 1 and segment 2. For each sequence segment, RDMs (*top*), MDS projections in phonological and semantic spaces (*middle*), and bar plots of distances between phonologically similar and different pairs, and semantically similar and different pairs (*bottom*) are shown. Statistical significance was assessed by one-sided two-sample $t$-tests (*$P < 0.05$, **$P < 0.01$, ***$P < 0.001$). Error bars indicate SEM. **b,** Visualizing representations of semantic model neurons in Whisper-large-v3 ($n$ = 1,280) across segment 1 and segment 2, plotted as in a. **c,** Visualizing representations of P2S-transfer model neurons in Whisper-large-v3 ($n$ = 462) across segment 1 and segment 2, plotted as in a.

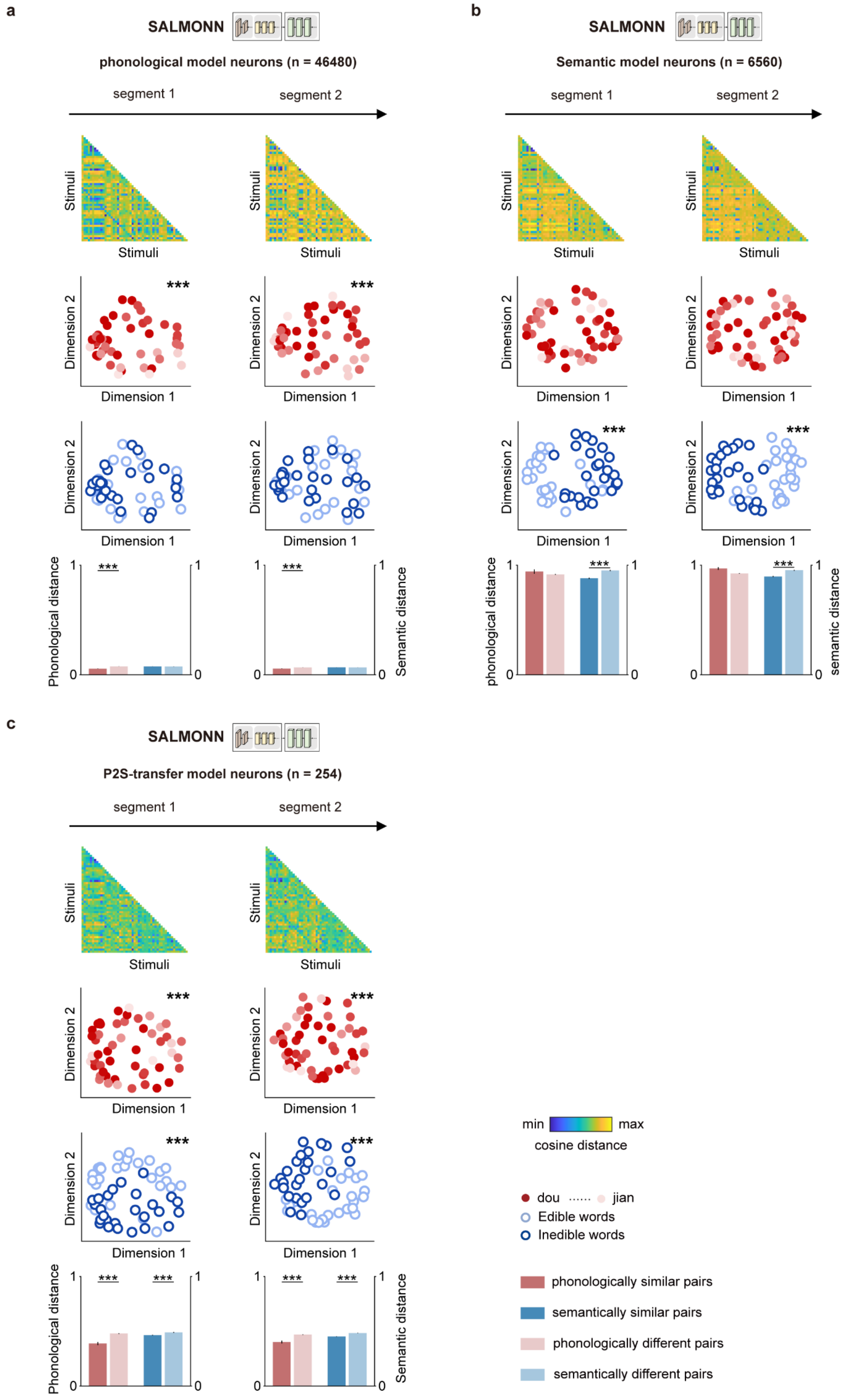
a
SALMONN
phonological model neurons (n = 46480)
segment 1
segment 2
Stimuli
Dimension 1
Dimension 2
***
Phonological distance
Semantic distance
b
SALMONN
Semantic model neurons (n = 6560)
phonological distance
semantic distance
c
SALMONN
P2S-transfer model neurons (n = 254)
min
max
cosine distance
dou
jian
Edible words
Inedible words
phonologically similar pairs
semantically similar pairs
phonologically different pairs
semantically different pairs

116

**Supplementary Fig. 10 | Representational geometry of phonological, semantic and P2S-transfer model neurons in SALMONN. a,** Visualizing representations of phonological model neurons in SALMONN ($n$ = 46,480) across segment 1 and segment 2. For each sequence segment, RDMs (*top*), MDS projections in phonological and semantic spaces (*middle*), and bar plots of distances between phonologically similar and different pairs, and semantically similar and different pairs (*bottom*) are shown. Statistical significance was assessed by one-sided two-sample $t$-tests (**$P < 0.01$, ***$P < 0.001$). Error bars indicate SEM. **b,** Visualizing representations of semantic model neurons in SALMONN ($n$ = 6,560) across segment 1 and segment 2, plotted as in a. **c,** Visualizing representations of P2S-transfer model neurons in SALMONN ($n$ = 254) across segment 1 and segment 2, plotted as in a.

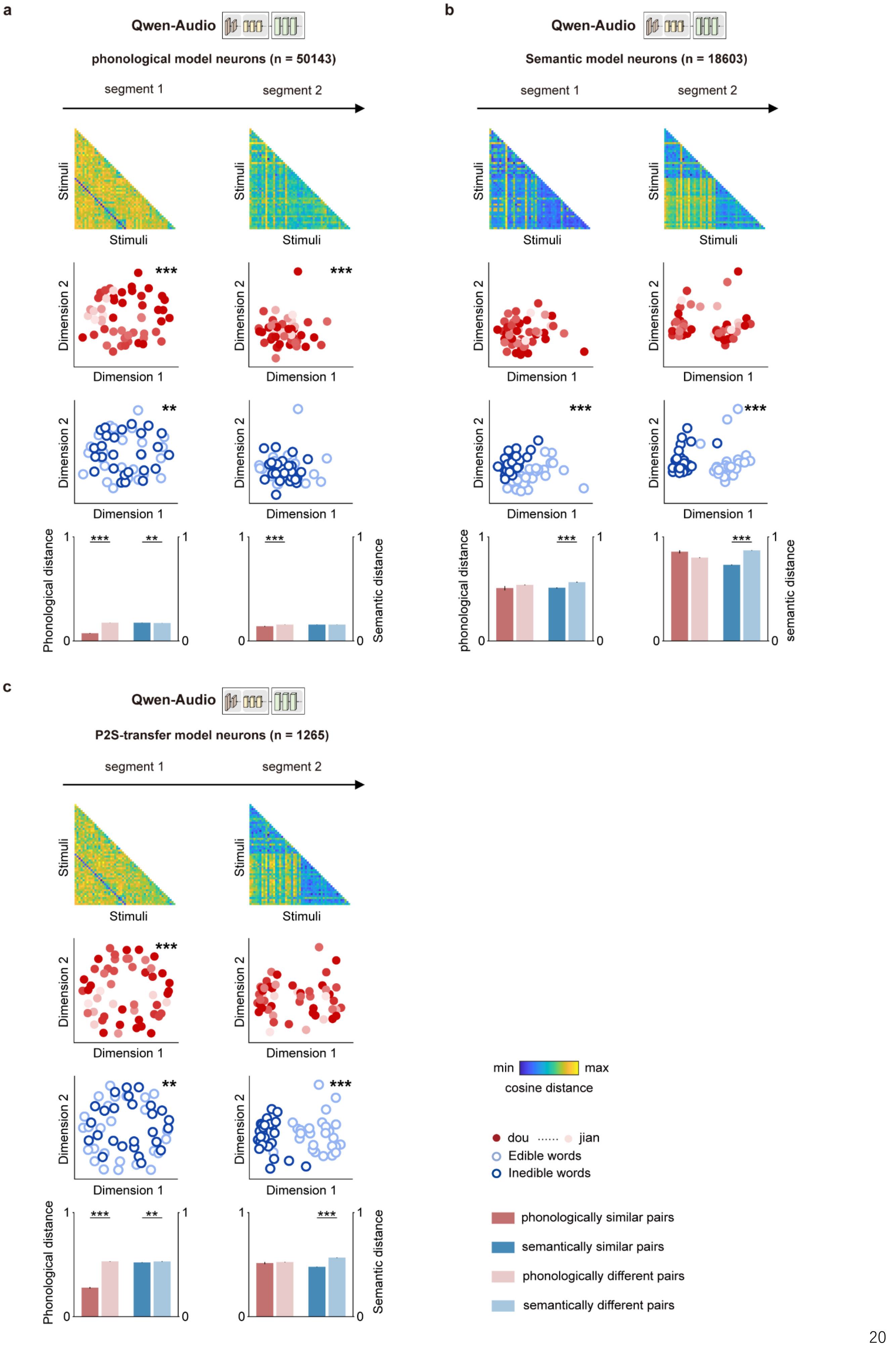

a
Qwen-Audio
phonological model neurons (n = 50143)
segment 1
segment 2
Stimuli
Dimension 1
Dimension 2
Phonological distance
Semantic distance
b
Qwen-Audio
Semantic model neurons (n = 18603)
segment 1
segment 2
phonological distance
semantic distance
c
Qwen-Audio
P2S-transfer model neurons (n = 1265)
segment 1
segment 2
min
max
cosine distance
dou
jian
Edible words
Inedible words
phonologically similar pairs
semantically similar pairs
phonologically different pairs
semantically different pairs


126

**Supplementary Fig. 11 | Representational geometry of phonological, semantic and P2S-transfer model neurons in Qwen-Audio. a,** Visualizing representations of phonological model neurons in Qwen-Audio ($n$ = 50,143) across segment 1 and segment 2. For each sequence segment, RDMs (*top*), MDS projections in phonological and semantic spaces (*middle*), and bar plots of distances between phonologically similar and different pairs, and semantically similar and different pairs (*bottom*) are shown. Statistical significance was assessed by one-sided two-sample $t$-tests (**$P < 0.01$, ***$P < 0.001$). Error bars indicate SEM. **b,** Visualizing representations of semantic model neurons in Qwen-Audio ($n$ = 18,603) across segment 1 and segment 2, plotted as in a. **c,** Visualizing representations of P2S-transfer model neurons in Qwen-Audio ($n$ = 1,265) across segment 1 and segment 2, plotted as in a.

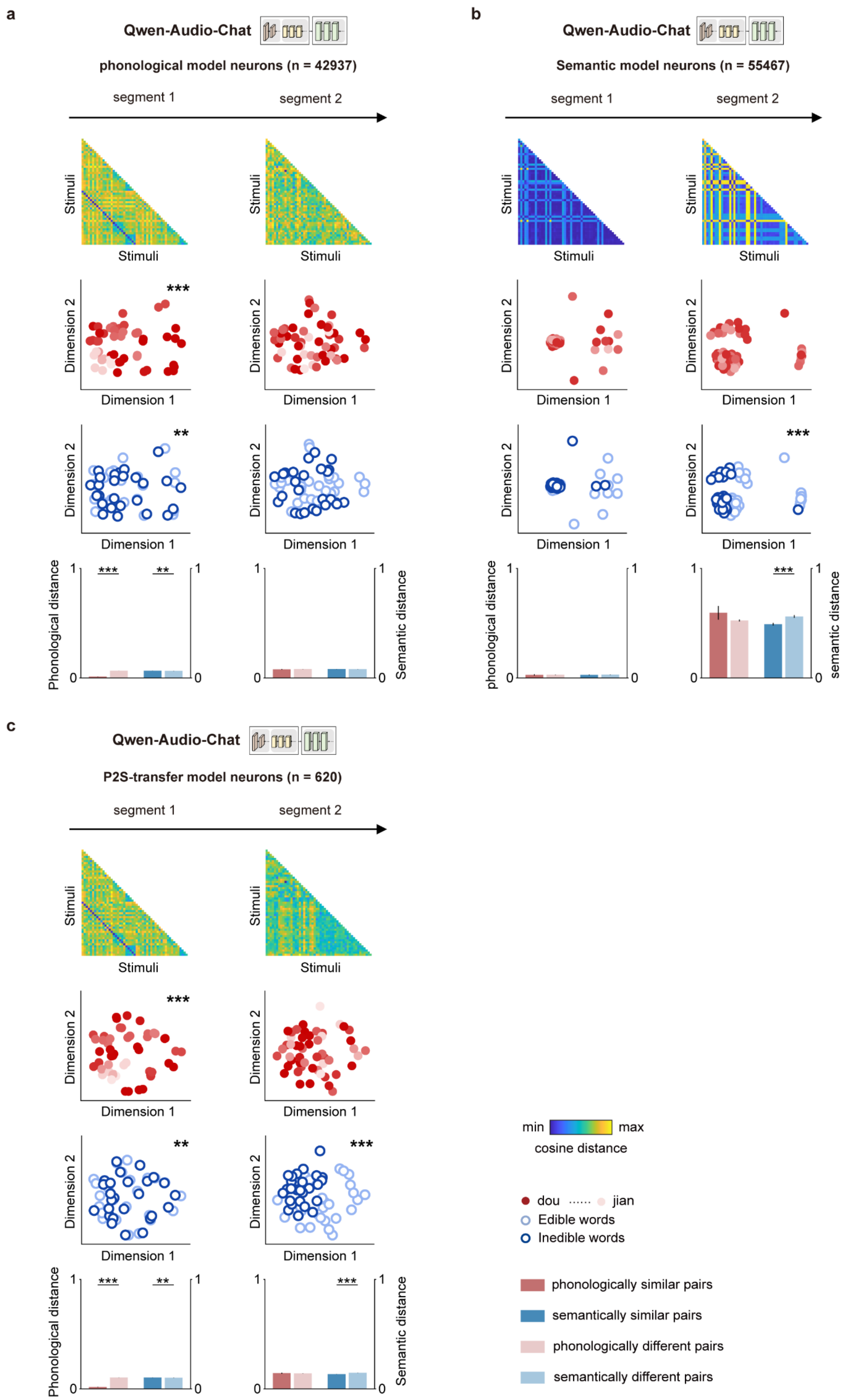

a
Qwen-Audio-Chat
phonological model neurons (n = 42937)
segment 1
segment 2
Stimuli
Dimension 1
Dimension 2
***
**
Phonological distance
Semantic distance
b
Qwen-Audio-Chat
Semantic model neurons (n = 55467)
segment 1
segment 2
phonological distance
semantic distance
c
Qwen-Audio-Chat
P2S-transfer model neurons (n = 620)
segment 1
segment 2
min
max
cosine distance
dou
jian
Edible words
Inedible words
phonologically similar pairs
semantically similar pairs
phonologically different pairs
semantically different pairs

**Supplementary Fig. 12 | Representational geometry of phonological, semantic and P2S-transfer model neurons in Qwen-Audio-Chat. a,** Visualizing representations of phonological model neurons in Qwen-Audio-Chat ($n$ = 42,937) across segment 1 and segment 2. For each sequence segment, RDMs (*top*), MDS projections in phonological and semantic spaces (*middle*), and bar plots of distances between phonologically similar and different pairs, and semantically similar and different pairs (*bottom*) are shown. Statistical significance was assessed by one-sided two-sample $t$-tests (**$P < 0.01$, ***$P < 0.001$). Error bars indicate SEM. **b,** Visualizing representations of semantic model neurons in Qwen-Audio-Chat ($n$ = 55,467) across segment 1 and segment 2, plotted as in a. **c,** Visualizing representations of P2S-transfer model neurons in Qwen-Audio-Chat ($n$ = 620) across segment 1 and segment 2, plotted as in a.

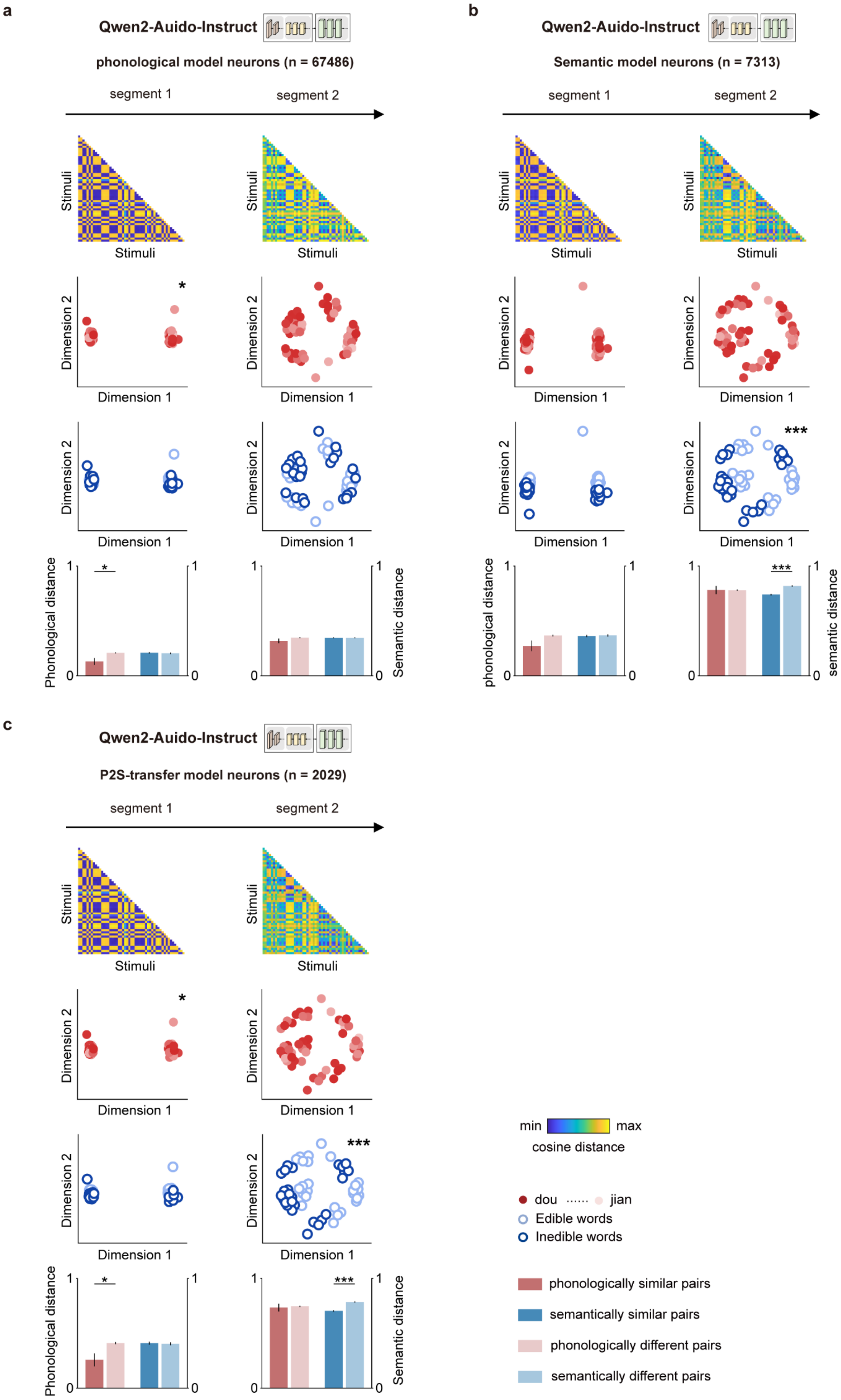
a
Qwen2-Auido-Instruct
phonological model neurons (n = 67486)
segment 1
segment 2
Stimuli
Dimension 1
Dimension 2
Phonological distance
Semantic distance
b
Qwen2-Auido-Instruct
Semantic model neurons (n = 7313)
segment 1
segment 2
phonological distance
semantic distance
c
Qwen2-Auido-Instruct
P2S-transfer model neurons (n = 2029)
segment 1
segment 2
min
max
cosine distance
dou
jian
Edible words
Inedible words
phonologically similar pairs
semantically similar pairs
phonologically different pairs
semantically different pairs

**Supplementary Fig. 13 | Representational geometry of phonological, semantic and P2S-transfer model neurons in Qwen2-Audio-Instruct. a,** Visualizing representations of phonological model neurons in Qwen2-Audio-Instruct ($n$ = 67,486) across segment 1 and segment 2. For each sequence segment, RDMs (*top*), MDS projections in phonological and semantic spaces (*middle*), and bar plots of distances between phonologically similar and different pairs, and semantically similar and different pairs (*bottom*) are shown. Statistical significance was assessed by one-sided two-sample $t$-tests (*$P < 0.05$, ***$P < 0.001$). Error bars indicate SEM. **b,** Visualizing representations of semantic model neurons in Qwen2-Audio-Instruct ($n$ = 7,313) across segment 1 and segment 2, plotted as in a. **c,** Visualizing representations of P2S-transfer model neurons in Qwen2-Audio-Instruct ($n$ = 2,029) across segment 1 and segment 2, plotted as in **a**.

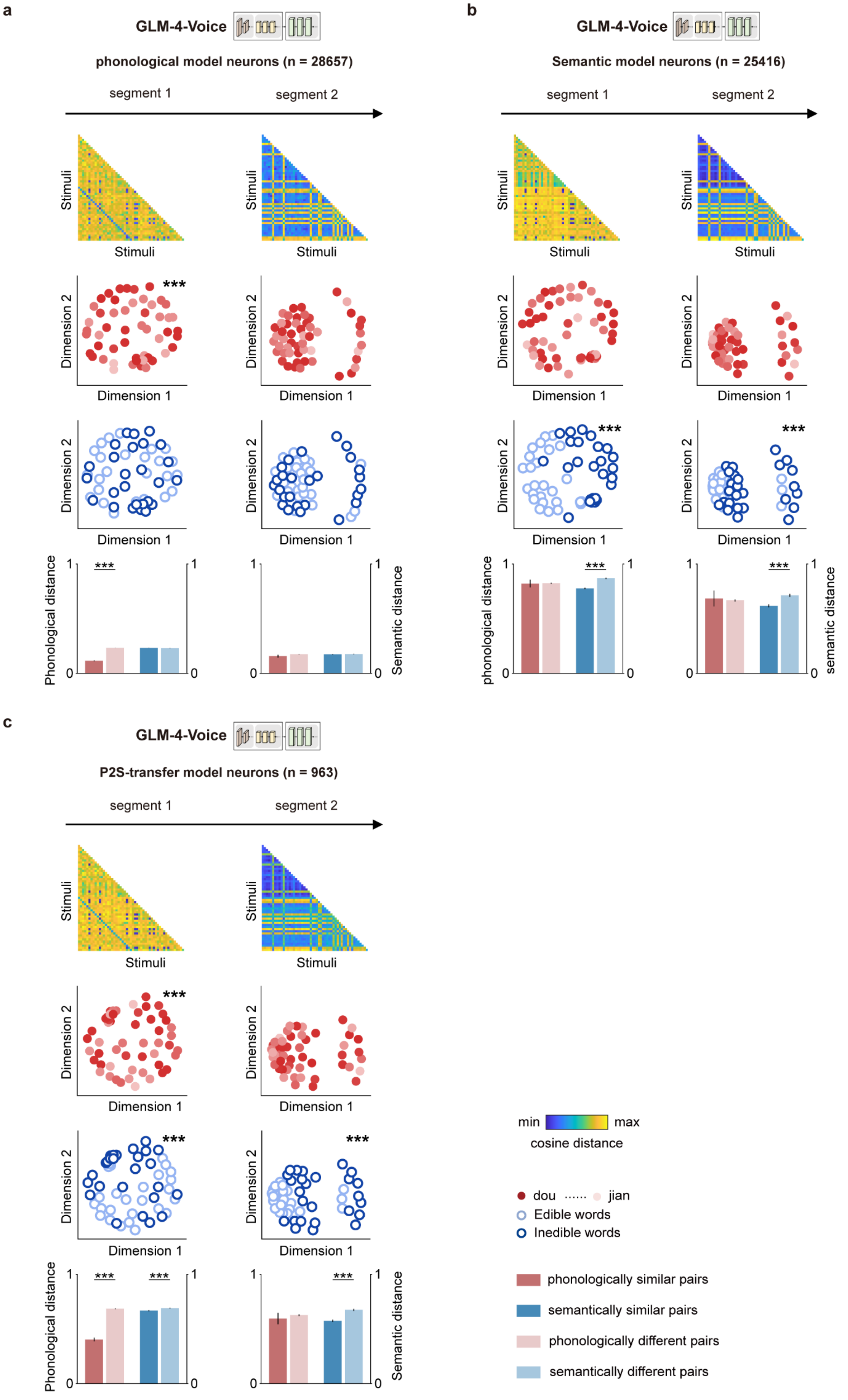

a
GLM-4-Voice
phonological model neurons (n = 28657)
segment 1
segment 2
Stimuli
Dimension 1
Dimension 2
***
Phonological distance
Semantic distance
b
GLM-4-Voice
Semantic model neurons (n = 25416)
segment 1
segment 2
Stimuli
Dimension 1
Dimension 2
***
phonological distance
semantic distance
c
GLM-4-Voice
P2S-transfer model neurons (n = 963)
segment 1
segment 2
Stimuli
Dimension 1
Dimension 2
***
Phonological distance
Semantic distance
0
1
min
max
cosine distance
dou
jian
Edible words
Inedible words
phonologically similar pairs
semantically similar pairs
phonologically different pairs
semantically different pairs

**Supplementary Fig. 14 | Representational geometry of phonological, semantic and P2S-transfer model neurons in GLM-4-Voice. a,** Visualizing representations of phonological model neurons in GLM-4-Voice ($n$ = 28,657) across segment 1 and segment 2. For each sequence segment, RDMs (*top*), MDS projections in phonological and semantic spaces (*middle*), and bar plots of distances between phonologically similar and different pairs, and semantically similar and different pairs (*bottom*) are shown. Statistical significance was assessed by one-sided two-sample $t$-tests (***$P < 0.001$). Error bars indicate SEM. **b,** Visualizing representations of semantic model neurons in GLM-4-Voice ($n$ = 25,416) across segment 1 and segment 2, plotted as in a. **c,** Visualizing representations of P2S-transfer model neurons in GLM-4-Voice ($n$ = 963) across segment 1 and segment 2, plotted as in a.

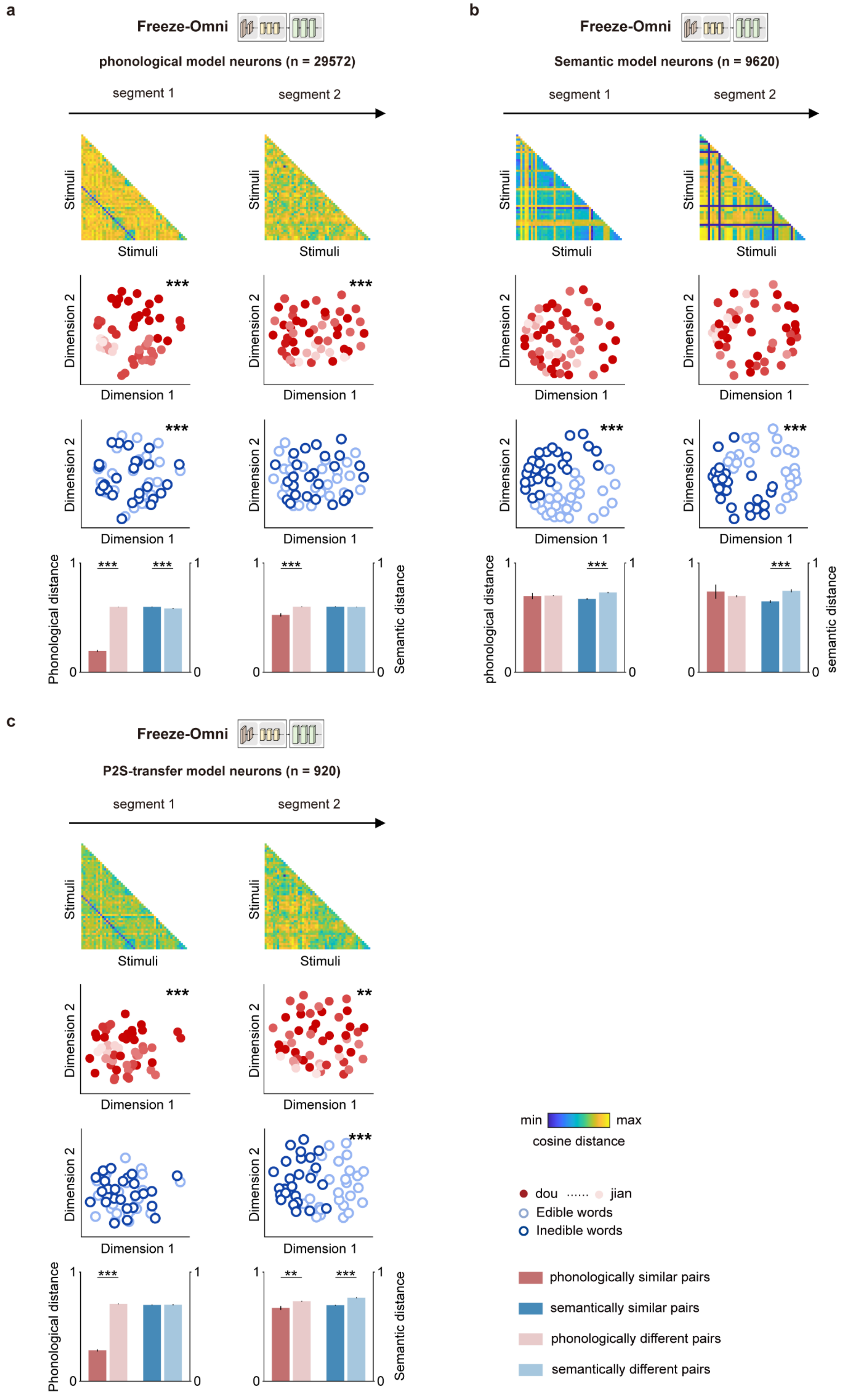

a
Freeze-Omni
phonological model neurons (n = 29572)
segment 1
segment 2
Stimuli
Dimension 1
Dimension 2
***
Phonological distance
Semantic distance
b
Freeze-Omni
Semantic model neurons (n = 9620)
segment 1
segment 2
phonological distance
semantic distance
c
Freeze-Omni
P2S-transfer model neurons (n = 920)
segment 1
segment 2
**
min
max
cosine distance
dou
jian
Edible words
Inedible words
phonologically similar pairs
semantically similar pairs
phonologically different pairs
semantically different pairs

**Supplementary Fig. 15 | Representational geometry of phonological, semantic and P2S-transfer model neurons in Freeze-Omni. a,** Visualizing representations of phonological model neurons in Freeze-Omni ($n$ = 29,572) across segment 1 and segment 2. For each sequence segment, RDMs (*top*), MDS projections in phonological and semantic spaces (*middle*), and bar plots of distances between phonologically similar and different pairs, and semantically similar and different pairs (*bottom*) are shown. Statistical significance was assessed by one-sided two-sample $t$-tests (**$P < 0.01$, ***$P < 0.001$). Error bars indicate SEM. **b,** Visualizing representations of semantic model neurons in Freeze-Omni ($n$ = 9,620) across segment 1 and segment 2, plotted as in a. **c,** Visualizing representations of P2S-transfer model neurons in Freeze-Omni ($n$ = 920) across segment 1 and segment 2, plotted as in a.

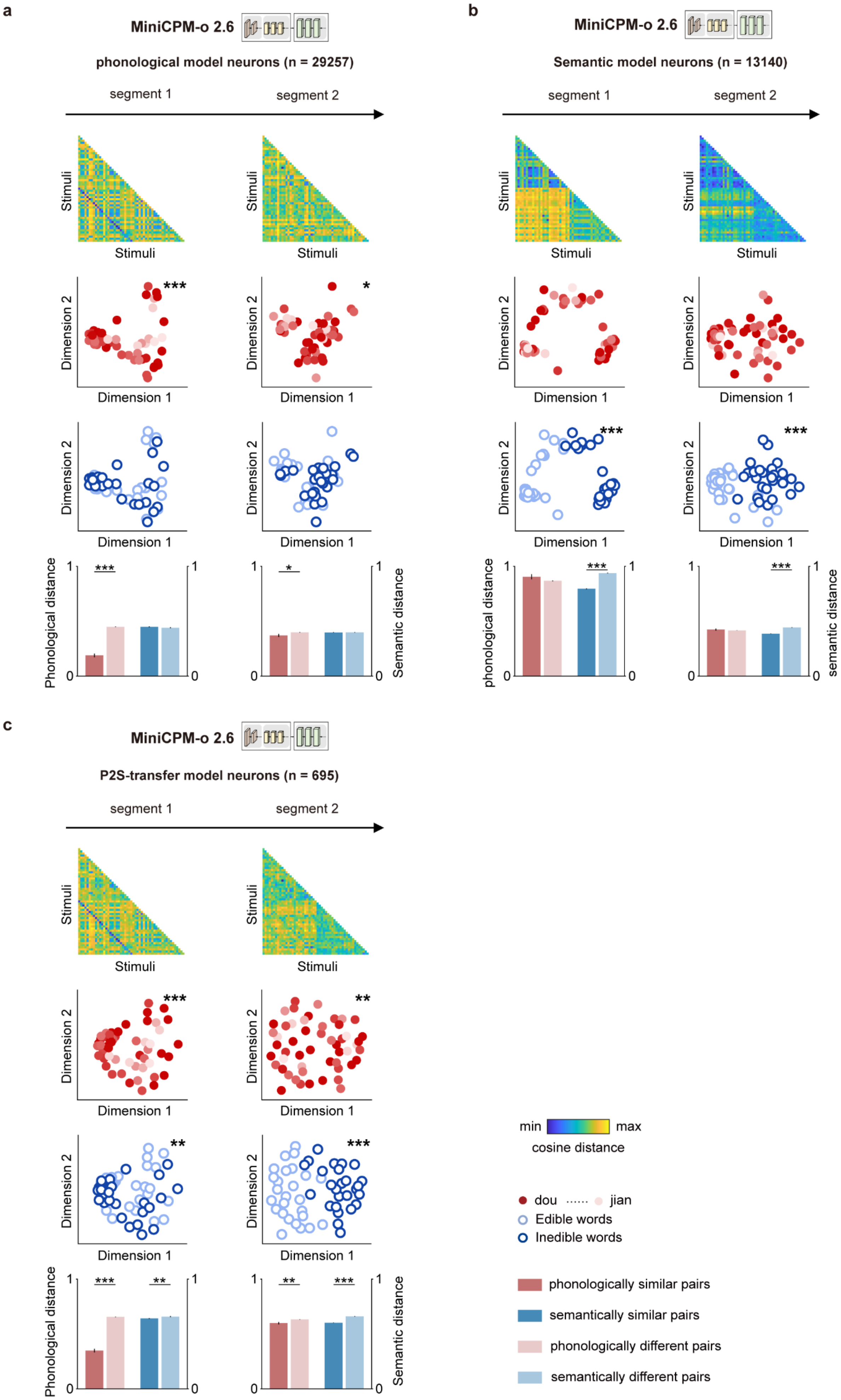

a
MiniCPM-o 2.6
phonological model neurons (n = 29257)
segment 1
segment 2
Stimuli
Dimension 1
Dimension 2
Phonological distance
Semantic distance
b
MiniCPM-o 2.6
Semantic model neurons (n = 13140)
segment 1
segment 2
Stimuli
Dimension 1
Dimension 2
phonological distance
semantic distance
c
MiniCPM-o 2.6
P2S-transfer model neurons (n = 695)
segment 1
segment 2
Stimuli
Dimension 1
Dimension 2
Phonological distance
Semantic distance
min
max
cosine distance
dou
jian
Edible words
Inedible words
phonologically similar pairs
semantically similar pairs
phonologically different pairs
semantically different pairs

**Supplementary Fig. 16 | Representational geometry of phonological, semantic and P2S-transfer model neurons in MiniCPM-o 2.6. a,** Visualizing representations of phonological model neurons in MiniCPM-o 2.6 ($n$ = 29,257) across segment 1 and segment 2. For each sequence segment, RDMs (*top*), MDS projections in phonological and semantic spaces (*middle*), and bar plots of distances between phonologically similar and different pairs, and semantically similar and different pairs (*bottom*) are shown. Statistical significance was assessed by one-sided two-sample $t$-tests (*$P < 0.05$, **$P < 0.01$, ***$P < 0.001$). Error bars indicate SEM. **b,** Visualizing representations of semantic model neurons in MiniCPM-o 2.6 ($n$ = 13,140) across segment 1 and segment 2, plotted as in a. **c,** Visualizing representations of P2S-transfer model neurons in MiniCPM-o 2.6 ($n$ = 695) across segment 1 and segment 2, plotted as in a.

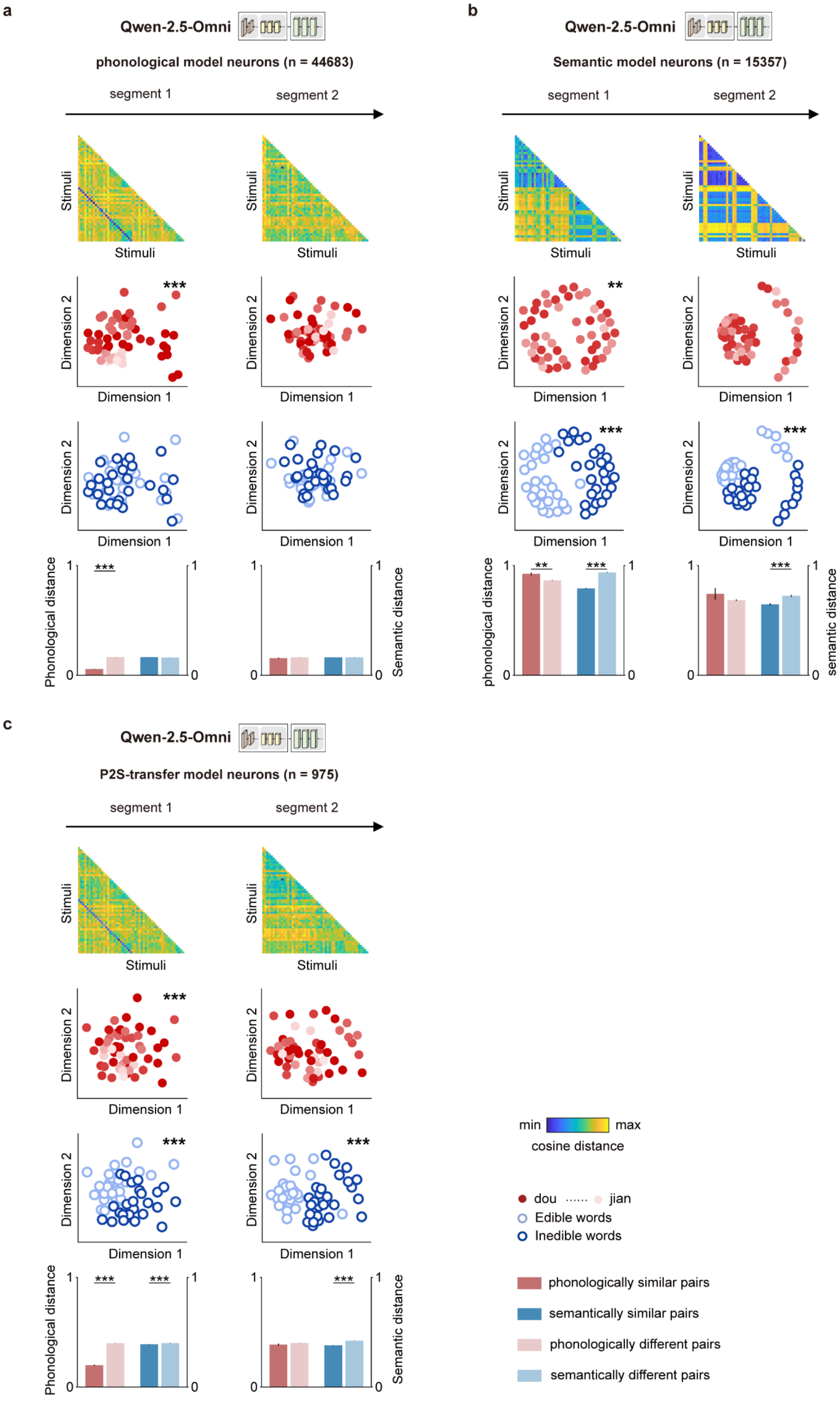
a
Qwen-2.5-Omni
phonological model neurons (n = 44683)
segment 1
segment 2
Stimuli
Dimension 1
Dimension 2
***
Phonological distance
Semantic distance
b
Qwen-2.5-Omni
Semantic model neurons (n = 15357)
segment 1
segment 2
Stimuli
Dimension 1
Dimension 2
**
***
phonological distance
semantic distance
c
Qwen-2.5-Omni
P2S-transfer model neurons (n = 975)
segment 1
segment 2
Stimuli
Dimension 1
Dimension 2
***
Phonological distance
Semantic distance
min
max
cosine distance
dou
jian
Edible words
Inedible words
phonologically similar pairs
semantically similar pairs
phonologically different pairs
semantically different pairs

**Supplementary Fig. 17 | Representational geometry of phonological, semantic and P2S-transfer model neurons in Qwen2.5-Omni. a,** Visualizing representations of phonological model neurons in Qwen2.5-Omni ($n$ = 44,683) across segment 1 and segment 2. For each sequence segment, RDMs (*top*), MDS projections in phonological and semantic spaces (*middle*), and bar plots of distances between phonologically similar and different pairs, and semantically similar and different pairs (*bottom*) are shown. Statistical significance was assessed by one-sided two-sample $t$-tests (**$P < 0.01$, ***$P < 0.001$). Error bars indicate SEM. **b,** Visualizing representations of semantic model neurons in Qwen2.5-Omni ($n$ = 15,357) across segment 1 and segment 2, plotted as in a. **c,** Visualizing representations of P2S-transfer model neurons in Qwen2.5-Omni ($n$ = 975) across segment 1 and segment 2, plotted as in a.

### Supplementary tables

**Supplementary Table 1 |** Demographic and experimental information.

| Patient No. | Age | Gender | Hemisphere | #Speech-responsive contacts |
|---|---|---|---|---|
| 01 | 41 | F | L | 91 |
| 02 | 22 | M | L | 75 |
| 03 | 37 | F | L, R | 49 |
| 04 | 21 | M | R | 90 |
| 05 | 26 | F | L, R | 54 |
| 06 | 25 | M | R | 38 |
| 07 | 16 | M | L | 54 |
| 08 | 31 | F | L, R | 107 |
| 09 | 32 | F | L | 48 |
| 10 | 25 | F | L | 29 |
| 11 | 30 | M | L | 44 |
| 12 | 29 | F | L, R | 55 |
| 13 | 13 | M | L, R | 30 |
| 14 | 42 | M | L | 30 |
| 15 | 41 | F | L | 42 |
| 16 | 32 | M | L | 61 |
| 17 | 27 | M | R | 65 |
| #Sum | | | | 962 |

**Note:** F, female; M, male; L, left hemisphere; R, right hemisphere; L, R, bilateral implantation. Speech-responsive contacts indicate the number of sEEG contacts showing significant speech-evoked high-γ responses.

**Supplementary Table 2 |** List of Mandarin disyllabic words used in the study.

| Word No. | Mandarin disyllabic words | Mandarin Chinese pinyin (Phonology) | First syllable | English translation | Semantic category |
|---|---|---|---|---|---|
| 1 | 煎饼 | *jian bing* | *jian* | pancake | edible |
| 2 | 西瓜 | *xi gua* | *xi* | watermelon | edible |
| 3 | 鸡蛋 | *ji dan* | *ji* | eggs | edible |
| 4 | 玉米 | *yu mi* | *yu* | corn | edible |
| 5 | 樱桃 | *ying tao* | *ying* | cherry | edible |
| 6 | 荔枝 | *li zhi* | *li* | lychee | edible |
| 7 | 绿豆 | *lv dou* | *lv* | mung bean | edible |
| 8 | 苹果 | *ping guo* | *ping* | apple | edible |
| 9 | 蜜桔 | *mi ju* | *mi* | honey tangerine | edible |
| 10 | 猪蹄 | *zhu ti* | *zhu* | pig's trotter | edible |
| 11 | 面包 | *mian bao* | *mian* | bread | edible |
| 12 | 薯条 | *shu tiao* | *shu* | French fries | edible |
| 13 | 萝卜 | *luo bo* | *luo* | radish | edible |
| 14 | 牛肉 | *niu rou* | *niu* | beef | edible |
| 15 | 香肠 | *xiang chang* | *xiang* | sausage | edible |
| 16 | 龙虾 | *long xia* | *long* | lobster | edible |
| 17 | 蘑菇 | *mo gu* | *mo* | mushroom | edible |
| 18 | 红薯 | *hong shu* | *hong* | sweet potato | edible |
| 19 | 软糖 | *ruan tang* | *ruan* | gummy candy | edible |
| 20 | 蛋糕 | *dan gao* | *dan* | cake | edible |
| 21 | 芒果 | *mang guo* | *mang* | mango | edible |
| 22 | 洋葱 | *yang cong* | *yang* | onion | edible |
| 23 | 带鱼 | *dai yu* | *dai* | hairtail | edible |
| 24 | 海带 | *hai dai* | *hai* | kelp | edible |
| 25 | 草莓 | *cao mei* | *cao* | strawberry | edible |
| 26 | 豆腐 | *dou fu* | *dou* | tofu | edible |
| 27 | 肩膀 | *jian bang* | *jian* | shoulders | inedible |
| 28 | 膝盖 | *xi gai* | *xi* | knee | inedible |
| 29 | 机会 | *ji hui* | *ji* | opportunity | inedible |
| 30 | 浴室 | *yu shi* | *yu* | bathroom | inedible |
| 31 | 英雄 | *ying xiong* | *ying* | hero | inedible |
| 32 | 力量 | *li liang* | *li* | power | inedible |
| 33 | 律师 | *lv shi* | *lv* | lawyer | inedible |
| 34 | 评委 | *ping wei* | *ping* | judges | inedible |
| 35 | 密码 | *mi ma* | *mi* | password | inedible |
| 36 | 珠宝 | *zhu bao* | *zhu* | jewelry | inedible |
| 37 | 面具 | *mian ju* | *mian* | mask | inedible |
| 38 | 暑假 | *shu jia* | *shu* | summer vacation | inedible |
| 39 | 螺旋 | *luo xuan* | *luo* | spiral | inedible |
| 40 | 牛仔 | *niu zai* | *niu* | cowboy | inedible |
| 41 | 乡村 | *xiang cun* | *xiang* | village | inedible |
| 42 | 笼子 | *long zi* | *long* | cage | inedible |
| 43 | 模特 | *mo te* | *mo* | model | inedible |
| 44 | 红灯 | *hong deng* | *hong* | red light | inedible |
| 45 | 软件 | *ruan jian* | *ruan* | software | inedible |
| 46 | 弹弓 | *dan gong* | *dan* | slingshot | inedible |
| 47 | 盲人 | *mang ren* | *mang* | blind person | inedible |
| 48 | 阳光 | *yang guang* | *yang* | sunshine | inedible |
| 49 | 代表 | *dai biao* | *dai* | representative | inedible |
| 50 | 海滩 | *hai tan* | *hai* | beach | inedible |
| 51 | 草坪 | *cao ping* | *cao* | lawn | inedible |
| 52 | 逗号 | *dou hao* | *dou* | comma | inedible |

**Supplementary Table 3** | SLMs used in this study, along with their performance and alignment indices across hierarchical and sequential dimensions.

| Models | Accuracy (Aishell-1) | Accuracy (Fleurs-zh) | Hierarchical alignment index | | | Sequential alignment index | | |
|---|---|---|---|---|---|---|---|---|
| | | | Phonological units | Semantic units | P2S-transfer units | Phonological units | Semantic units | P2S-transfer units |
| XLSR-53-ch | 79.74 | 76.15 | 0.1045 | 0.6755 | 0.0557 | 0.9355 | 0.5192 | 0.4622 |
| Whisper-large-v3 | 89.41 | 81.71 | 0.4680 | 0.5816 | 0.5971 | 0.8992 | 0.8391 | 0.6429 |
| LLaSM | 2.84 | 4.84 | 0.5069 | 0.7453 | -0.0300 | 0.9030 | 0.9520 | 0.2033 |
| SALMONN | 43.01 | 52.61 | 0.5076 | 0.6196 | -0.0637 | 0.9031 | 0.9507 | 0.2996 |
| Qwen-Audio | 98.51 | 84.33 | 0.5067 | 0.5053 | 0.2010 | 0.9328 | 0.8922 | 0.6513 |
| Qwen-Audio-Chat | 98.43 | 91.14 | 0.5010 | 0.6579 | -0.0907 | 0.9319 | 0.8033 | 0.4575 |
| Qwen2-Audio | 96.20 | 90.90 | 0.5630 | 0.7417 | 0.4681 | 0.8872 | 0.8453 | 0.8862 |
| Qwen2-Audio-Instruct | 97.11 | 89.86 | 0.5816 | 0.8007 | 0.4866 | 0.9010 | 0.5862 | 0.8327 |
| GLM-4-Voice | 81.54 | 74.49 | 0.4739 | 0.4251 | -0.1639 | 0.9340 | 0.9525 | 0.7406 |
| Freeze-Omni | 65.93 | 68.37 | 0.5190 | 0.6056 | -0.1411 | 0.9412 | 0.9399 | 0.4798 |
| MiniCPM-o 2.6 | 97.79 | 93.22 | 0.5061 | 0.5407 | -0.1366 | 0.9339 | 0.9312 | 0.6020 |
| Qwen2.5-Omni | 97.20 | 93.69 | 0.4011 | 0.6254 | -0.3799 | 0.9311 | 0.9280 | 0.5462 |

**Supplementary Table 4** | Example model speech transcriptions with different lesion types.

| Lesion type | Model transcription | | Ground-truth transcription |
|---|---|---|---|
| | Qwen2-Audio | Qwen-Audio | |
| Intact model | 因此，铅笔一问世就成了许多人的心头好 *English translation:* Therefore, "pencil" became a favorite of many people as soon as it was introduced. | 因此，铅笔一问世就成了许多人的心头好 *English translation:* Therefore, "pencil" became a favorite of many people as soon as it was introduced. | 因此，铅笔一问世就成了许多人的心头好 *English translation:* Therefore, "pencil" became a favorite of many people as soon as it was introduced. |
| Phonological units | 因此，铅笔一问世就成了许多人的心头好 *English Translation:* Therefore, "pencil" became a favorite of many people as soon as it was introduced. | 因此，铅笔一卫士就成了许多人的心头好 *English translation:* Therefore, "pencil" "one Wei Shi" became a favorite of many people as soon as it was introduced. | |
| Semantic units | 因此，铅笔一问世就成了许多人的心头好 *English translation:* Therefore, "pencil" became a favorite of many people as soon as it was introduced. | 因此，铅笔一问世就成了许多人的心头好 *English translation:* Therefore, "pencil" became a favorite of many people as soon as it was introduced. | |
| P2S-transfer units | 因此，铅笔一问世就成了许多人的新头号 *English translation:* Therefore, "pencil" became a favorite of many people's new top favorite as soon as it was introduced. | 因此，千笔一问式就成了许多人的心头号 *English translation:* Therefore, "Qian Bi Yi Wen Shi" became a favorite of many people's hearts. | |
